# Discovery and characterisation of gene by environment and epistatic genetic effects in a vertebrate model

**DOI:** 10.1101/2025.04.24.650462

**Authors:** Bettina Welz, Saul Pierotti, Tomas Fitzgerald, Thomas Thumberger, Risa Suzuki, Philip Watson, Jana Fuss, Tiago Cordeiro da Trindade, Fanny Defranoux, Marcio Ferreira, Kiyoshi Naruse, Felix Loosli, Jakob Gierten, Joachim Wittbrodt, Ewan Birney

## Abstract

Phenotypic variation arises from the interplay between genetic and environmental factors. However, disentangling these interactions for complex traits remains challenging in observational cohorts such as human biobanks. Instead, model organisms where genetic (G) and environmental (E) variation can be controlled offer a valuable complement to human studies in the analysis of higher-order genetic effects such as GxE interactions, dominance, and epistasis. Here, we utilized 76 medaka strains of the Medaka Inbred Kiyosu-Karlsruhe (MIKK) panel, to compare heart rate plasticity across temperatures. An F2 segregation analysis identified 16 quantitative trait loci (QTLs), with many exhibiting dominance, GxE, GxG, and GxGxE interactions. We experimentally validated four candidate genes using gene editing, revealing their temperature-sensitive impact on heart function. Finally, we devised simulations to assess how GWAS discovery power is influenced by the choice of statistical models, showing that the apparent additivity in human GWAS is to be expected given study design and sample sizes of current studies. This work demonstrates the value of controlled model organism studies for dissecting the genetics of complex traits and provides guidance on the design of genetic association studies.

## Introduction

Understanding the causes of phenotypic variation is a fundamental question in genetics, past (Fisher, 1919) and present (Herrera-Luis et al., 2024). While statistical arguments (Hill et al., 2008) and practical observations (Hivert et al., 2021; Hu et al., 2025; Yang et al., 2010) suggest that additive genetic effects can explain a large part of the heritability for many phenotypes in humans, ample evidence in model organisms (Baud et al., 2017; Bloom et al., 2013; Chen et al., 2023; Lachance et al., 2013; T. F. C. Mackay, 2014; T. F. C. Mackay & Anholt, 2024; Mott et al., 2014; Tonnelé & Baud, 2022; Wright et al., 2022), plant breeding (Anshori et al., 2024, 2024; Cooper & Messina, 2021; Napier et al., 2023), and animal breeding (Copley et al., 2024; Fennewald et al., 2018; Hammond, 1947; Hasan et al., 2024; Hayes et al., 2016; Kebede et al., 2024; Sae-Lim et al., 2025) suggests that non-additive effects such as gene-by-environment (G×E) and epistatic (G×G) interactions are critical for understanding the genetic architecture underlying complex traits. These effects are also important for other tasks in genetics, such as accurate phenotype prediction. Nonetheless, even in human genetics, the importance of non-additive effects for realising the goals of precision health is well recognised (Bakermans-Kranenburg & IJzendoorn, 2015; Motsinger-Reif et al., 2024) and is at the heart of the field of pharmacogenomics (Russell et al., 2021). Moreover, several examples of environmentally-dependent effects in humans have been reported (Bononi et al., 2020; Caspi et al., 2003; Hawn et al., 2018; Johnson et al., 2010; Modafferi et al., 2021; Polimanti et al., 2018; Rampersaud et al., 2008), as well as other forms of non-additivity such as dominance (Palmer et al., 2023) and epistatic effects (Currant et al., 2023).

Genome-Wide Association Studies (GWAS) in humans have been a resounding success (Abdellaoui et al., 2023), fuelled by the establishment of large-scale cohorts for genetic research (Bick et al., 2024; Bycroft et al., 2018; Gaziano et al., 2016; Kurki et al., 2023; Leitsalu & Metspalu, 2017; Nagai et al., 2017; Sijtsma et al., 2022). However, human studies are limited in most cases to an observational approach, with major confounding deriving from population structure and genotype-environment correlations (Abdellaoui et al., 2022; Sul et al., 2018; Vilhjálmsson & Nordborg, 2013). In contrast, the literature on model organisms and plant or animal breeding genetics has a long history of leveraging controlled experimental settings to dissect the origins of phenotypic variation. Notable mentions are Multi-parent Advanced Generation Intercross (MAGIC) populations in plants (Kover et al., 2009; I. J. Mackay et al., 2014; Scott et al., 2020, 2021), laboratory and wild strains in yeast (Brem et al., 2002; Ho & Zhang, 2014; Jelier et al., 2011; Ohya et al., 2005; Smith & Kruglyak, 2008), structured populations in worms (Evans et al., 2021; Noble et al., 2017; Snoek et al., 2019; Widmayer et al., 2022), the Drosophila Genetic Reference Panel (DGPR) (T. F. C. Mackay et al., 2012; T. F. C. Mackay & Huang, 2018), the mouse Collaborative Cross (Churchill et al., 2004; Collaborative Cross Consortium, 2012; Srivastava et al., 2017), and heterogenous stocks in rodents (Baud et al., 2013; Mott et al., 2000).

Studies specifically designed to identify non-linear effects and to account for environmental influences on complex traits in model organisms indicate great complementary potential (Rohde et al., 2019; Huang et al., 2020a; Maazi et al., 2019; Willis-Owen & Valdar, 2009). Mice in particular, as mammals, have a high translational relevance for human biology due to genetic, anatomical and physiological similarities. However, the genetic variation that is covered by the available laboratory mouse strains (The Complex Trait Consortium, 2004; Svenson et al., 2012; Saul et al., 2019; Ashbrook et al., 2021; Dumont et al., 2024), with complex domestication histories (Morse, 2012; Wade & Daly, 2005), does not reflect genetic variability in the wild. Thus, extending the analysis of complex traits to a vertebrate system that reflects the genetic diversity of a wild population promises to provide further insights on natural genetic variation. It remains an open question how much the acknowledged deficiencies of using additive models for genetic discovery impact on our understanding of human biology.

In this study, we used medaka (*Oryzias latipes*), a well-established vertebrate model, to investigate gene-by-environment (G×E) and epistatic (G×G) interactions affecting embryonic heart rate in the context of wild-derived haplotypes (**Figure 1**). Medaka has long been used as model organism in research, with numerous wild-derived inbred strains and molecular tools available (Aida, 1921; Wittbrodt et al., 2002; Spivakov et al., 2014; Kirchmaier et al., 2015). Its high fecundity, *ex-utero* embryonic development, short generation time, small 700 Mb genome, and cost-effective husbandry make it particularly suitable for large-scale genetic studies (Furutani-Seiki & Wittbrodt, 2004; Kasahara et al., 2007). Here, we leveraged the Medaka Inbred Kiyosu-Karlsruhe (MIKK) panel — a collection of 80 inbred strains derived from a wild population (Fitzgerald et al., 2022) — to study naturally occurring genetic and phenotypic variation in embryonic heart rate under controlled temperature conditions.

**Figure 1.**
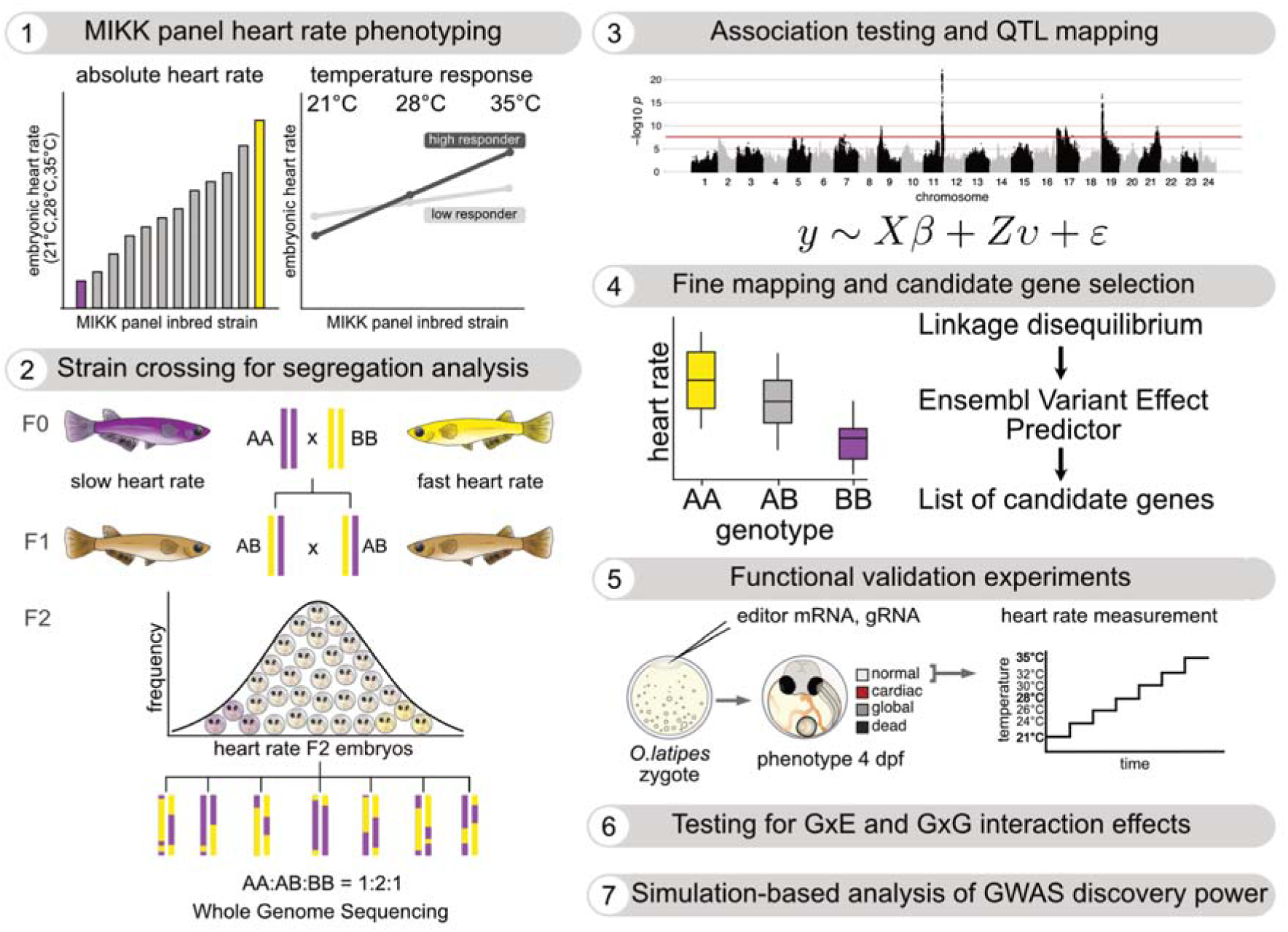
Study design for the identification and validation of genetic variants underlying embryonic heart rate differences at changing temperature conditions. (1) The heart rate spectrum of genetically diverse MIKK panel inbred strains was assessed 4 days post fertilization (dpf) at 21°C, 28°C and 35°C. (2) Eight strains exhibiting contrasting baseline heart rates and differences in heart rate changes upon temperature treatment were crossed to each other to segregate phenotypic differences within a F2 mapping population. (3) Association testing and quantitative trait loci (QTL) mapping were performed to map genetic variants linked to the observed divergence in heart rate. (4) Fine mapping aided the identification of candidate genes harbouring genetic variants with a predicted effect on protein function. (5) To experimentally validate the identified candidate genes, different gene editing tools were applied to assess functional consequences on heart rate in a temperature-dependent manner. (6) Gene-by-environment (G×E) and epistatic (G×G) interaction effects were tested for their potential contribution to heart rate variation across the measured temperatures. (7) Simulations based on the experimental data set reveal insights on the discovery power of association studies under model misspecification.

Embryonic heart rate in fish is a readily measurable physiological trait influenced by temperature and thus enables the functional examination of a conserved organ system (Jensen et al., 2013; Gierten et al., 2020). We examined heart rate genetics at three ecologically relevant temperatures — two moderate (21°C, 28°C) and one extreme but still within regularly recorded ecological limits (35°C) — using whole-genome sequencing data from 2,209 F2 embryos derived from 11 crosses among 8 MIKK panel inbred strains. We identified 16 genetic loci driving heart rate differences and used them to explore G×E and G×G interactions. Using gene editing technologies, we further experimentally validated the effects of identified candidate genes on heart rate across temperature conditions.

Finally, we used simulations informed by our experimental results to address the impact of the statistical model chosen for discovery, and to make recommendations on the key factors driving discoverability in human genetic association studies. We confirm that additive models are adequate for the discovery of the majority of loci in environmentally heterogeneous observational studies as long as the causal variant is directly used in the discovery and there is a reasonable measurement of the environment. We confirm that with the observed effect sizes of non-additive effects from this medaka study we would not be able to discover interaction effects in human GWAS study designs. Finally, we also highlight scenarios where additive models are not adequate and the challenges in the interpretation of the resulting discovered loci when additive models are used. Together, this work highlights the power of medaka as a model system for genetic research, sheds light on the genetic architecture of complex traits using the example of heart rate and provides guidance for the design and analysis of future genetic association efforts.

## Results

### Temperature-dependent heart rate variation of MIKK panel inbred strains is under genetic control and can be dissected by a segregation analysis

We first assessed how genetic factors affect heart rate variation in the context of ecologically-relevant temperature conditions, by measuring embryonic heart rates of 76 wild-derived medaka inbred strains in response to successive treatments at moderate (21°C, 28°C) and more extreme (35°C) temperature conditions (**Figure 2A, 2B**); 4 MIKK strains were not fecund enough to support this assessment. Heart rate measurements of 1,671 embryos were acquired 4 days post-fertilization (dpf) in a highly randomized and high-throughput set up, with a minimum of five individuals and up to 47 individuals phenotyped per strain. We observed that the median heart rate of the strains differed by as much as 26 beats per minute (bpm) (31% difference) at 21 °C, 39 bpm (27% difference) at 28 °C and 58 bpm (29% difference) at 35 °C. In addition to strain-dependent absolute heart rate differences at the tested temperature conditions, the degree of heart rate change according to the temperature treatment varied across the strains (**Figure S1A**). The comparison of the median heart rate change between 21°C and 35°C revealed clear strain-dependent differences. While an increase in heart rate of only 106 bpm was measured in the strain with the lowest response (22-1), an increase in heart rate to the temperature change (21°C vs 35°C) of up to 155 bpm was observed in the strain with the strongest response (7-2). To test for significance of the genetic strain-dependent effect on heart rate, we ran Kruskal-Wallis tests stratified by temperature treatment. For all the three temperatures, we observed a Kruskal-Wallis *p*-value smaller than 2.2 * 10^-16^.

**Figure 2.**
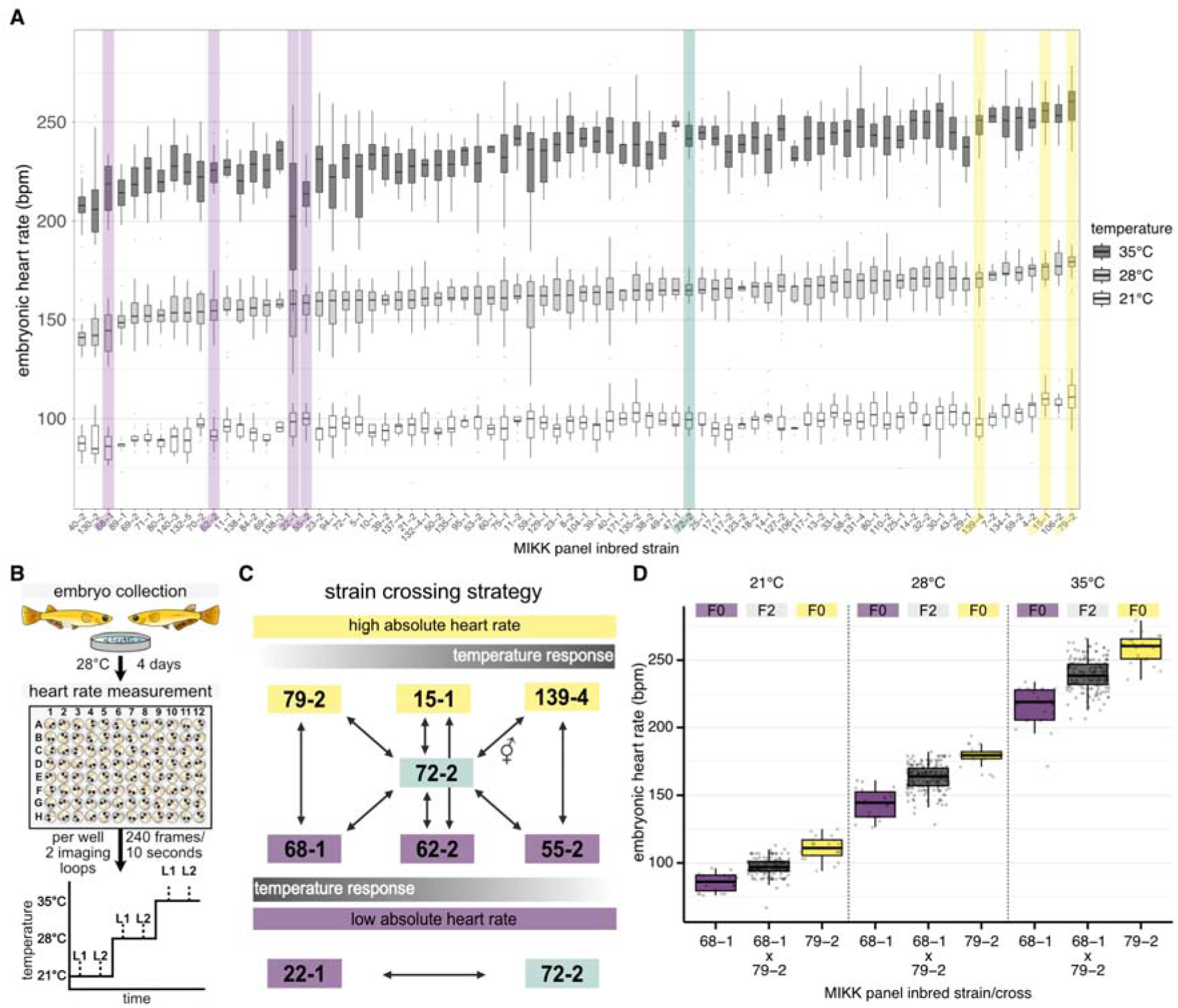
Embryonic heart rate spectrum of MIKK panel strains at 21°C, 28°C, 35°C and heart rate segregation through strategic crossing. **A:** Strain-dependent heart rate differences in beats per minute (bpm) for 76 MIKK inbred strains 4 days post fertilization (dpf) at 21°C, 28°C and 35°C. Strains were ranked based on their median baseline heart rate at the standard temperature 28°C (n = 5 to 47 embryos). Selected strains for segregation analysis are highlighted (slow heart rate, varying temperature response: purple; average heart rate: green, fast heart rate, varying temperature response: yellow). Data is displayed as box plots (median line and box showing the interquartile range between the 25th and 75th percentiles). **B:** Experimental workflow of the embryonic heart rate assessment 4 dpf. Image sequences were acquired after a 20 min acclimatization period at the respective temperature condition within two consecutive imaging loops (L1, L2) from which the average heart rate has been calculated. **C:** Strategic crossing strategy of eight MIKK panel strains with different heart rate phenotypes at the changing temperature conditions as basis for segregation analysis and genetic mapping. **D:** Heart rate of recombinant F2 embryos (grey) spreads between the heart rate of parental strains (F0; purple and yellow) across the measured temperatures, shown for one representative cross (F2 heart rates for all crosses cf. **Figure S1B**). Data is displayed as box plots (median line and box showing the interquartile range between the 25th and 75th percentiles) and overlaid scatter plots of individual heart rate measurements in beats per minute (bpm). Sample sizes (n) for 21 °C, 28 °C and 35 °C: 68-1 x 79-2 F2 embryos n = 180, 181, 179; 68-1 F0 n = 20, 20, 21; 79-2 F0 n = 21, 22, 22.

To estimate the broad-sense heritability of the phenotype at each temperature treatment ANOVA models were fitted to this dataset, stratified by temperature. The estimated heritability was 0.42 (95% C.I.: 0.40 to 0.49) at 21 °C, 0.45 (95% C.I.: 0.43 to 0.51) at 28 °C and 0.44 (95% C.I.: 0.43 to 0.51) at 35 °C. Although this assessment was clear evidence of genetic as well as G×E effects, it cannot reveal the specific loci underlying these phenomena. To identify the genetic variants contributing to the observed trait differences we conducted a segregation analysis as a basis for genetic mapping. We selected eight isogenic MIKK panel strains with different heart rate and temperature response characteristics (**Figure 2A, S1A**). In total, we performed eleven cross combinations (**Figure 2C**); seven MIKK panel strains with extreme heart rates and temperature response phenotypes were crossed with the average-phenotype MIKK panel strain 72-2. One of these crosses was performed reciprocally, by respectively using maternal and paternal fish from the two founding strains. In addition, three crosses among strains with the greatest phenotypic contrast along the heart rate and temperature response axes were performed. From each cross we set up mating groups with the heterozygous F1 offspring in order to establish F2 populations. These were used to assay embryonic heart rates at 21°C, 28°C and 35°C and we generated shallow whole genome sequencing (WGS) data for 2,209 F2 individuals.

We previously described the whole genome sequencing of this F2 population in the context of optimising population design for genotype imputation from shallow WGS data (Pierotti et al., 2024). We were able to impute a set of 3.2 million SNPs with a mean per-sample squared Pearson correlation of 0.996 on high-coverage validation samples (see **Methods**). The majority of F2 individuals exhibited heart rate values that are intermediate compared to the heart rate of the respective cross founder strains (**Figure 2D and S1B**). Nevertheless, a subset of crosses gave rise to F2 individuals with more extreme heart rates than those of the cross founders. As expected, we observed that F2 individuals deriving from the same founder strains exhibited higher degrees of genetic relatedness compared to F2 individuals from independent crosses (**Figure S2**). Within the reciprocal cross (72-2x139-4 and 139-4x72-2) we observed a sub-clustering of the F2 individuals, indicating some remaining degree of heterozygosity in the fish used to set up the crosses.

Consistent with our F0 inbred strain assessment, mixed model-based heritability estimates for heart rate were determined to be 0.35 at 21 °C (95% C.I.: 0.28 to 0.42), 0.46 at 28 °C (95% C.I.: 0.40 to 0.53), and 0.45 at 35 °C (95% C.I.: 0.38 to 0.52). Heritability for the difference in heart rate across temperatures (which we call the temperature response phenotype) was 0.31 between 28 and 21 °C (95% C.I.: 0.25 to 0.38), 0.34 between 35 and 21 °C (95% C.I.: 0.28 to 0.41), and 0.33 between 35 and 28 °C (95% C.I.: 0.27 to 0.40).

### Association testing reveals 16 QTLs linked to heart rate and underlying pervasive G×E and G×G interaction effects

A linear mixed model GWAS analysis of the combined genotype/phenotype F2 dataset revealed the presence of 16 quantitative trait loci (QTLs) that passed the phenotype-specific significance threshold (set by permutations) in at least one of the measured temperature conditions (**Figure 3A and 3B**, **Table 1**). Additionally, we ran the GWAS analysis on the difference in heart rate for each F2 embryo following temperature change (**Figure S3**). In these temperature response phenotypes, 7 of the 16 QTLs that we observed in the stratified analysis remained detectable, while 9 were not. No novel loci were identified in the response phenotypes that had not already been observed in the stratified analysis. However, certain loci (e.g., *chr15_qtl*, *chr2_qtl*) exhibited stronger association signals for the response phenotype compared to the single-temperature heart rate analysis, improving fine mapping potential. Given the large magnitude of the effect observed by a locus on chromosome 15 (*chr15_qtl*), we decided to include the lead SNP as a covariate before performing further analyses (**Figure 3B and S3B**). This also allowed for a more clear identification of the remaining loci. Overall, our complex F2 cross structure proved to be well-powered to identify genetic associations with heart rate. This is particularly true for low-frequency, high-effect alleles, which are theoretically enriched to a 50% frequency in F2 crosses when homozygous divergent in the founder strains. However, due to long F2 haplotypes in the cross from different F0 founders, our F2 population is not designed for high resolution fine-mapping of association signals.

**Figure 3.**
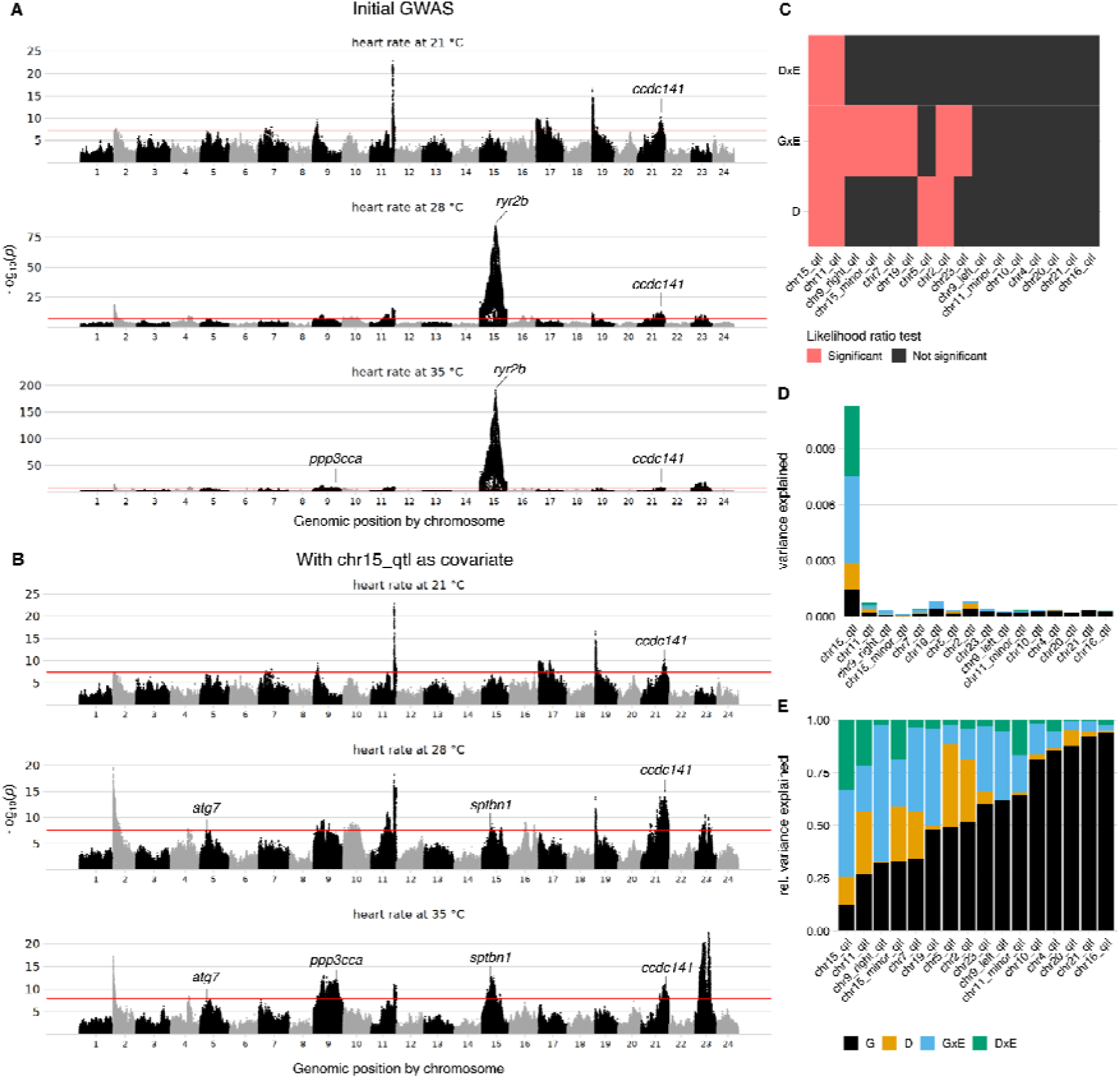
GWAS results reveal 16 QTLs and G×G/G×E interaction effects underlying heart rate variance. **A:** Manhattan plot for the association of heart rate with genetics, stratified by environment (temperature). The significance threshold (red line) corresponds to the minimum *p*-value achieved over 100 permutations. The genes selected for experimental validation are indicated. **B:** Same as panel A, but with the chromosome 15 locus included as a covariate in the model to unmask weaker association signals. **C:** Heatmap of significant test results for dominance (D), gene-by-environment interactions (G×E), and dominance-by-environment interactions (D×E) terms, for each of the discovered loci. **D:** Percentage of variance explained by genetic terms (G; black), dominance terms (D; orange), and gene-by-environment (G×E; blue), and dominance-by-environment (D×E; green) terms, for each of the 16 discovered loci. **E:** Same as panel D, but showing the relative variance explained as a proportion of the total variance attributable to the G, D, G×E, and D×E terms.

**Table 1.**
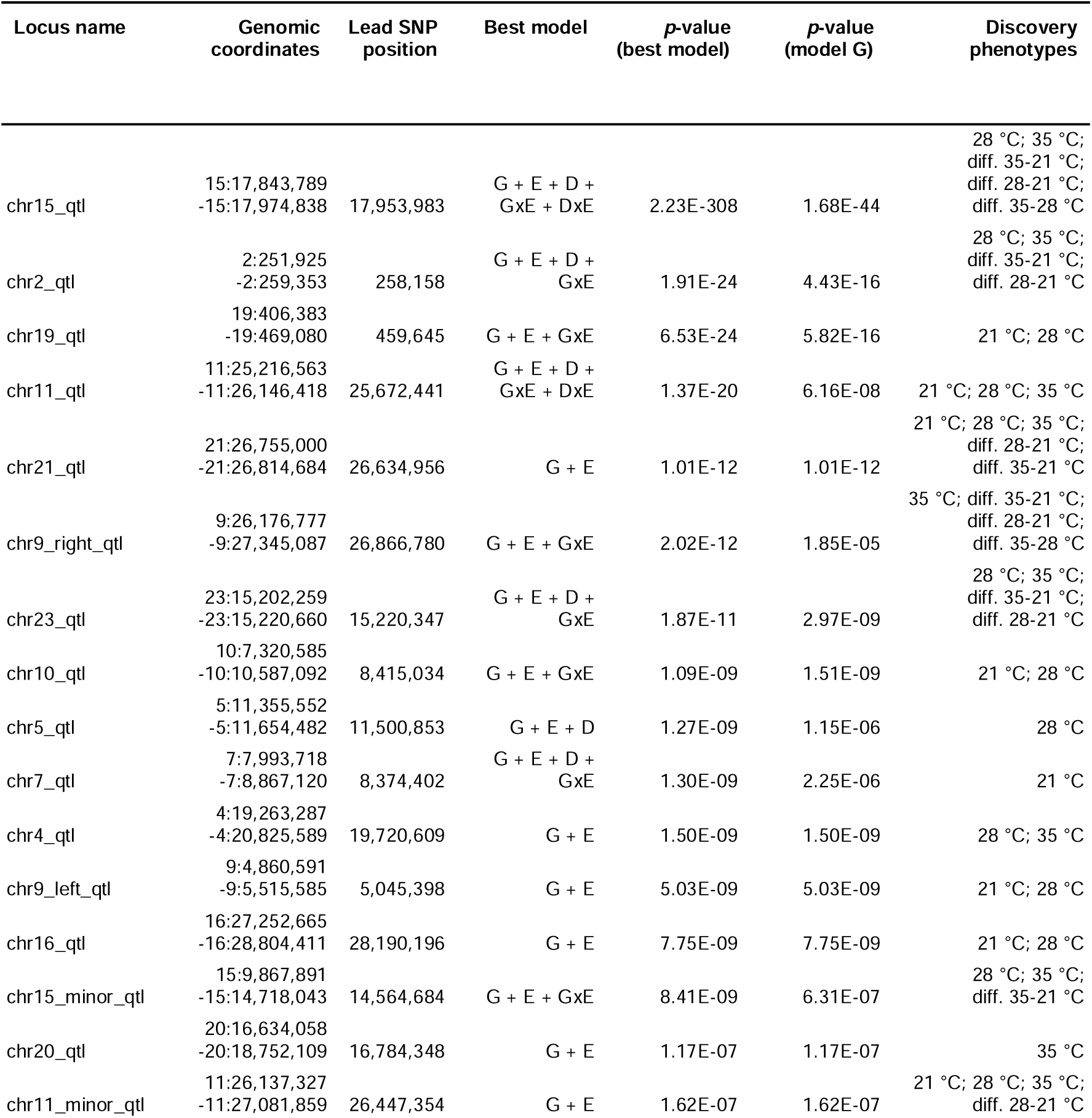
Overview of the 16 mapped quantitative trait loci (QTLs). The best model is the one with the best fit in likelihood ratio tests against a covariate-only model (while accounting for the increase in degrees of freedom in more complex models with the choice of the appropriate *x*^2^ distribution for testing the LRT statistics). In addition to the covariates, the G + E model contains the lead SNP for the locus additively encoded; the G + E + G×E model contains the SNP and its interaction with temperature; the G + E + D model includes the SNP and a dominance term; the G + E + D + G×E model includes the SNP, a dominance term, and the interaction of the SNP with temperature; the G + E + D + G×E + D×E model includes the SNP, a dominance term, and both of their interactions with temperature. Locus boundaries were manually determined by inspecting the association profiles. The lead SNP was usually selected as the one with the strongest association, but in some cases a different SNP was selected if the strongest associated SNP was missing a homozygous state. Note: the *p*-values are not extracted from the whole-genome discovery run, but they were obtained from models fit to all the temperatures simultaneously (see **Methods**). The “Discovery phenotypes” column indicates the phenotypes where the QTL was determined to be significant in the whole-genome discovery run for each temperature (21 °C, 28 °C, 35 °C) and comparing heart rate changes from measurements at different (diff.) temperatures (35-21 °C; 28-21 °C; 35-21 °C). For all the loci except for chr15_qtl, the lead SNP of chr15_qtl and its interactions with temperature were included as covariates.

| Locus name | Genomic coordinates | Lead SNP position | Best model | p-value (best model) | p-value (model G) | Discovery phenotypes |
| --- | --- | --- | --- | --- | --- | --- |
| chr15_qtl | 15:17,843,789<br>-15:17,974,838 | 17,953,983 | G + E + D +<br>GxE + Dx E | 2.23E-308 | 1.68E-44 | 28 °C; 35 °C;<br>diff. 35-21 °C;<br>diff. 28-21 °C;<br>diff. 35-28 °C |
| chr2_qtl | 2:251,925<br>-2:259,353 | 258,158 | G + E + D +<br>GxE | 1.91E-24 | 4.43E-16 | 28 °C; 35 °C;<br>diff. 35-21 °C;<br>diff. 28-21 °C |
| chr19_qtl | 19:406,383<br>-19:469,080 | 459,645 | G + E + GxE | 6.53E-24 | 5.82E-16 | 21 °C; 28 °C |
| chr11_qtl | 11:25,216,563<br>-11:26,146,418 | 25,672,441 | G + E + D +<br>GxE + Dx E | 1.37E-20 | 6.16E-08 | 21 °C; 28 °C; 35 °C |
| chr21_qtl | 21:26,755,000<br>-21:26,814,684 | 26,634,956 | G + E | 1.01E-12 | 1.01E-12 | 21 °C; 28 °C; 35 °C;<br>diff. 28-21 °C;<br>diff. 35-21 °C |
| chr9_right_qtl | 9:26,176,777<br>-9:27,345,087 | 26,866,780 | G + E + GxE | 2.02E-12 | 1.85E-05 | 35 °C; diff. 35-21 °C;<br>diff. 28-21 °C;<br>diff. 35-28 °C |
| chr23_qtl | 23:15,202,259<br>-23:15,220,660 | 15,220,347 | G + E + D +<br>GxE | 1.87E-11 | 2.97E-09 | 28 °C; 35 °C;<br>diff. 35-21 °C;<br>diff. 28-21 °C |
| chr10_qtl | 10:7,320,585<br>-10:10,587,092 | 8,415,034 | G + E + GxE | 1.09E-09 | 1.51E-09 | 21 °C; 28 °C |
| chr5_qtl | 5:11,355,552<br>-5:11,654,482 | 11,500,853 | G + E + D | 1.27E-09 | 1.15E-06 | 28 °C |
| chr7_qtl | 7:7,993,718<br>-7:8,867,120 | 8,374,402 | G + E + D +<br>GxE | 1.30E-09 | 2.25E-06 | 21 °C |
| chr4_qtl | 4:19,263,287<br>-4:20,825,589 | 19,720,609 | G + E | 1.50E-09 | 1.50E-09 | 28 °C; 35 °C |
| chr9_left_qtl | 9:4,860,591<br>-9:5,515,585 | 5,045,398 | G + E | 5.03E-09 | 5.03E-09 | 21 °C; 28 °C |
| chr16_qtl | 16:27,252,665<br>-16:28,804,411 | 28,190,196 | G + E | 7.75E-09 | 7.75E-09 | 21 °C; 28 °C |
| chr15_minor_qtl | 15:9,867,891<br>-15:14,718,043 | 14,564,684 | G + E + GxE | 8.41E-09 | 6.31E-07 | 28 °C; 35 °C;<br>diff. 35-21 °C |
| chr20_qtl | 20:16,634,058<br>-20:18,752,109 | 16,784,348 | G + E | 1.17E-07 | 1.17E-07 | 35 °C |
| chr11_minor_qtl | 11:26,137,327<br>-11:27,081,859 | 26,447,354 | G + E | 1.62E-07 | 1.62E-07 | 21 °C; 28 °C; 35 °C;<br>diff. 28-21 °C |

Our experimental design, which evaluated the heart rate of the same individuals across multiple temperature environments, allowed for direct investigation of gene-by-environment (G×E) interactions. Indeed, the identification of loci significantly associated with temperature response phenotypes, coupled with distinct association patterns for heart rate at different temperatures, provides strong evidence for G×E interactions in our population. A qualitative example is the QTL detected on chromosome 15 (*chr15_qtl*). While this locus showed no association with heart rate at 21 °C, it was significantly associated with heart rate at 28 °C and 35 °C. As expected, given this striking difference, *chr15_qtl* exhibited highly significant associations with all response phenotypes. To quantitatively assess interaction effects and further characterise the genetic architecture of the heart rate phenotype, we formally modelled the significance and fraction of variance explained by various interaction terms in our dataset. Overall, we observed that 8 of our 16 QTLs exhibit significant G×E effects, and 4 exhibit significant dominance effects (**Figure 3C**). In addition, 2 loci are significant for interactions between the dominance term and the temperature (which we call DxE). Of the 16 loci, only 7 are not significant for any interaction term (including dominance) and are thus fitting a purely additive model. In terms of variance explained, we notice a large difference among loci in the fraction of the overall phenotypic variance explained by the locus that is of purely additive origin (**Figure 3D and 3E**). *chr15_qtl* is the locus with the least linear effect, with less than 13% of the variance of additive origin. On the contrary, *chr16_qtl* is the locus that most closely resembles pure additivity, with almost 94% of the variance of additive origin.

Finally, we tested whether the paternal versus maternal founding strain status had an effect on the detected QTLs using the reciprocal crosses (72-2x139-4 and 139-4x72-2, n = 302). We found no evidence of interaction between the reciprocal cross and genotype at any of the 16 QTLs after Bonferroni correction (**Data S9**). On the contrary, when we looked into the marginal effect of the reciprocal cross status independent of genetics, we were able to detect significant effects on heart rate after Bonferroni correction at all the temperature treatments (21 °C: p = 8.06e-03; 28 °C: p = 7.29e-08; 35 °C: p = 5.18e-04).

Besides looking at dominance and G×E interactions, we also assessed how different loci interact with each other in terms of epistatic (G×G) interactions (**Figure 5A**); the sample size of the multi-way F2 cross is powered to see major G×G effects between specific loci. In the ideal scenario of 2 interacting G×G loci, both at 50% allele frequency in the same F2 cross and no linkage, one would expect 6.25% of samples in each of the minority classes (double homozygous genotypes). For our average F2 cross of 220 samples, this translates to approximately 14 samples in the smallest genotype class. To control for the multiple testing burden, we decided to only look at interactions between the 16 QTLs that we discovered. After multiple testing corrections, we observed that 21 of 120 possible QTL pairs are significant for either G×G or G×G×E effects. Only 4 QTL pairs are significant uniquely for G×G effects, while the majority of the G×G interactions are themselves temperature-dependent (G×G×E). 12 loci pairs are uniquely significant for G×G×E interactions but not G×G, while 5 loci pairs are significant for both G×G and G×G×E interactions. In this case, a locus being significant for G×G×E but not for G×G indicates that the marginal G×G effect is undetectable on its own, but the epistatic interaction becomes evident when the environmental context is considered.

### Experimental validation of identified candidate genes by gene editing phenocopies heart rate differences across temperatures

To experimentally test whether the identified variants are causal and influence heart rate in response to temperature, we selected five candidate gene*s: atg7, ccdc141, ppp3cca, ryr2b,* and *sptbn1* for functional validation experiments (**Table 2**). For the candidate gene selection, we focused on genetic variants that are in strong linkage with the lead SNP and have effect predictions in the coding regions with severe consequences (e.g. premature stop codon or frameshift mutations) and examined related phenotypic descriptions (see Methods “Candidate gene selection”). The *ccdc141*, *ryr2b* and *sptbn1* genes are linked to cardiac traits (see **Discussion**). We used CRISPR/Cas9 or a cytosine base editor for gene perturbations by microinjecting the editor mRNA together with a gRNA into 1-cell stage embryos of the Cab medaka strain, which is commonly used as wildtype reference. The choice of gene editing tool (base editor or CRISPR/Cas9) for functional validation experiments was made in a locus-specific manner. For genes harbouring a genetic variant with a stop gained predicted effect, we used the cytosine base editor evoBE4max to introduce a premature stop codon in close proximity to the originally observed variant (*ryr2b* and *atg7*). The CRISPR/Cas9 system was applied to investigate variants that resulted in a frameshift mutation (*ccdc141*), or when narrowing down to a single putative causal variant was not straightforward (*ppp3cca*, *sptbn1*). Hereafter, we refer to the embryos edited using CRISPR/Cas9 as "crispants", and to the embryos edited using a base editor as "editants". In a two-step phenotyping process of the edited embryos, we examined consequences on embryo morphology and heart rate 4 days post injection. Editing of the selected candidate genes resulted 4 days post injection in an increased proportion of embryos with heart-specific morphological effects (9-30%) and global phenotypes (16-45%) compared to the mock-injected control group (heart-specific morphological effects: 5%, global phenotypes: 2%) (**Figure S4A and S4C**). The cardiac affected embryos showed looping defects, pericardial edema, resized atrial or ventricular chambers, arrhythmia and dextrocardia (**Figure S4B**). To determine whether the heart rate profiles of the edited embryos are altered in a temperature-dependent context, we exposed injected embryos to an expanded temperature ramp (21°C, 24°C, 26°C, 28°C, 30°C, 32°C, 35°C). We assessed heart rate changes in morphologically wild type crispants and editants (and did not consider cardiac and global affected embryos). Editing of the selected candidate genes revealed significant heart rate effects for four of the five tested loci (**Figure 4B**). While editing of three genes (*ccdc141*, *ppp3cca* and *sptbn1*) led to a heart rate increase, a heart rate decrease was observed for the *ryr2b* editants. For the *ryr2b* and the *ccdc141* genes, we had identified a putative causal allele (a stop gain and frameshift). In these two cases, the gene editing effect recapitulated the effect direction expected from the QTL discovery phase. We detected a temperature-independent genetic effect for *ryr2b* (chr15_qtl), *ccdc141* (chr21_qtl), *sptbn1* (chr15_minor_qtl), and temperature-dependent genetic effects for *ryr2b, ccdc141, ppp3cca, sptbn1* at a 5% false discovery rate. Some degree of inter-individual variability in the effect size among the crispants and editants (injected generation) was observed for all of the candidate genes. This is likely due to genetic mosaicism in the injected generation, and it was statistically accounted for as a random effect for the individual embryo.

**Figure 4.**
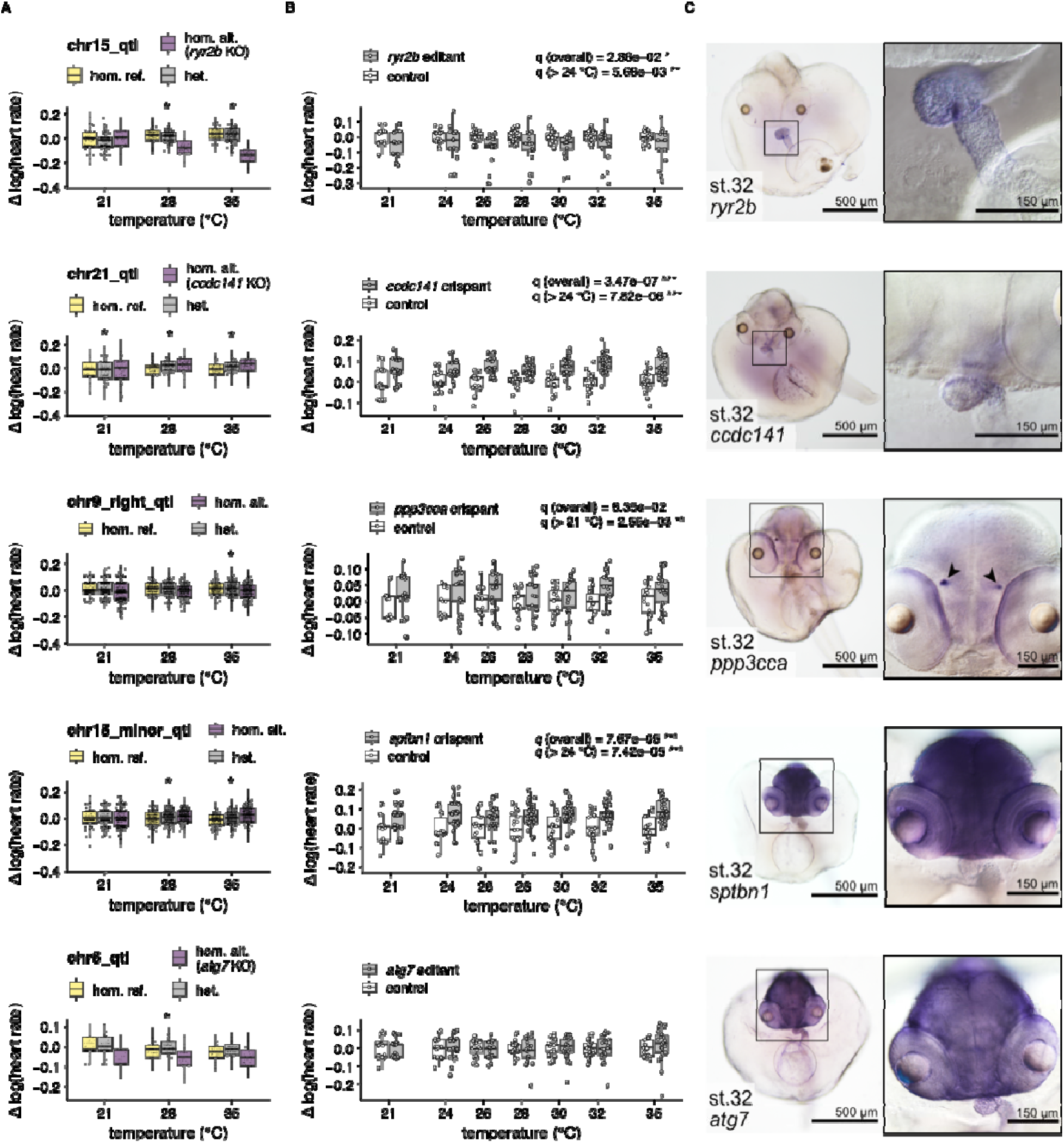
Experimental characterization of candidate genes confirms temperature-dependent heart rate differences through gene editing and reveals their expression domains. **A:** Boxplot showing the genotype effect on heart rate at different temperatures across the F2 embryos for 5 selected QTLs. Heart rate of F2 individuals was compared based on the genotype state (homozygous wildtype reference, yellow; heterozygous, grey; homozygous alternate, purple). For ease of visualisation, the effect of temperature was removed by subtracting the mean phenotype at any given temperature. A log transformation is used to stabilise the variance. **B:** Heart rate assessment 4 days post injection of normally developed crispants (grey; n = 21-38; cf. Data S5, Data S6) and editants (grey, n = 17-23; cf. Data S5, Data S6) in comparison to mock-injected control embryos (white; n = 15-24; cf. Data S5, Data S6) at temperatures ranging from 21°C to 35°C. *p*-values are reported for the presence of a temperature-independent (G), and temperature-dependent (G×E) effect (see **Methods**). The *y*-axis is log-transformed and centred on the mean of the control samples for each temperature. Data is displayed as box plots (median +/- interquartile range between the 25th and 75th percentiles) and overlaid scatter plots of individual heart rate measurements. **C:** Wholemount *in situ* hybridization displays expression patterns of the identified candidate genes at stage 32 in the albinism Heino strain.

**Table 2.** Overview of the five candidate gene(s) selected for experimental validation, QTL, variant(s), tissue of expression, and wild allele frequencies. Candidate genes and causal variants were manually selected according to their position, pattern across the mapping population, and known effect or functionality.

| Candidate gene | Locus name | Putative causal variant position | VEP consequence | Tissue of expression | Wild allele frequency causal variant | MIKK panel allele frequency |
| --- | --- | --- | --- | --- | --- | --- |
| *ryr2b* | chr15_qtl | 18,044,484;<br>18,043,338;<br>18,006,656 | stop gained;<br>frameshift;<br>stop gained | heart | 0.030;<br>not observed;<br>0.008 | 0.027;<br>0.007;<br>0.02 |
| *ccdc141* | chr21_qtl | 26,749,290 | frameshift | heart | 0.005 | 0.05 |
| *ppp3cca* | chr9_right_qtl | - | - | retinal pigment epithelium;<br>habenula | - | - |
| *sptbn1* | chr15_minor_qtl | - | - | ubiquitous except heart | - | - |
| *atg7* | chr5_qtl | 9,430,242 | stop gained | ubiquitous | 0.202 | 0.14 |

For *ryr2b* we detected a gene perturbation effect on the heart rate conditional on the temperature being above 24 °C, consistent with what we observed in the F2 GWAS data. This temperature-dependent effect was larger than the marginal gene editing effect. For the *ccdc141* crispants we were also able to detect a temperature-independent heart rate effect (mimicking the discovery results). For *ppp3cca* we were not able to detect an overall effect, but a heart rate effect was detected conditional on the temperature being above 21 °C. This differs from the discovery GWAS, where the QTL was detected as significant only for the heart rate measured at 35 °C. Nonetheless, in the discovery GWAS also at 28 °C a comparable, though not above our discovery threshold, an association profile is observable for the locus (**Figure 4A**). For *sptbn1*, the measured effect was more significant when conditioning on the temperature being above 24°C, consistent with the discovery GWAS. Finally, for the *atg7* editants we were not able to detect an effect on heart rate across the different temperatures.

In summary, we experimentally confirmed functional consequences on the heart rate in response to temperature for four of the five investigated candidate genes; in each of the four cases the effect validated by gene editing was broadly consistent with our F2 cross discovery GWAS.

To further characterize the candidate genes associated with heart rate in medaka, we analysed their expression patterns by wholemount *in situ* hybridization at the embryonic stage 32, where heart rate was also measured. While some of the candidate genes were specifically expressed in the heart (*ccdc141*, *ryr2b*) indicating a local mechanism of action, others did not show a cardiac expression (*sptbn1, ppp3cca)* (**Figure 4C**). The gene *atg7* showed a ubiquitous expression pattern, whereas *sptbn1* was ubiquitously expressed except in heart tissue. Interestingly, the candidate gene *ppp3cca* was expressed in the habenula, a part of the brain, and in the retinal pigment epithelium in the eyes, both suggesting a secondary effect on heart rate. The different expression domains of the candidates identified by genetic mapping suggest diverse cellular and molecular mechanisms influencing cardiac function across the MIKK panel strains.

Since heart rate is a complex trait that can be influenced by multiple factors, we next sought to experimentally investigate the phenotypic consequences of an identified variant, taking into account possible genetic interactions. To experimentally investigate how epistatic (G×G) interactions might influence phenotypic penetrance we targeted in different genetic backgrounds the candidate gene *ccdc141*, for which an environmentally-dependent G×G interaction (G×G×E) was detected in the F2 GWAS data (see **Figure 5A**). To achieve this, we injected the CRISPR/Cas9 reagents into different medaka strains (Cab, MIKK-panel strains 14-1, 72-2 and 106-2). While editing of *ccdc141* robustly increased the heart rate in the Cab reference strain and in the MIKK-panel strain 72-2, we were unable to detect a significant heart rate change upon editing of *ccdc141* in the MIKK-panel strains 14-1 and 106-2 (**Figure 5C**). The strain-dependence of the genetic effect is statistically significant (p = 8.98 * 10^-3^) and it provides evidence for the presence of a G×G interaction between the *ccdc141* edit and the overall genetic background of the strains.

**Figure 5.**
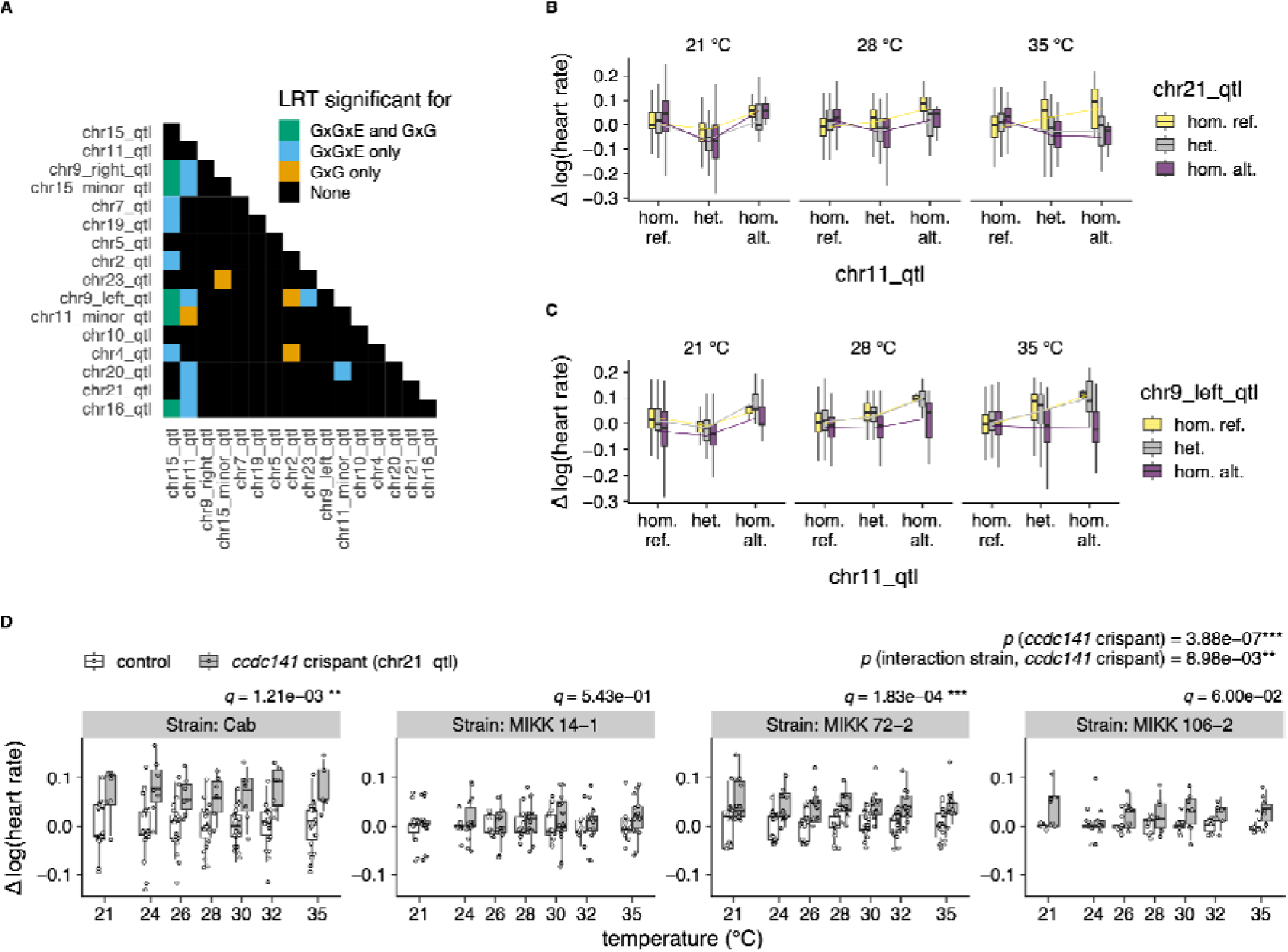
Detection and experimental investigation of G×G and G×G×E interaction effects. **A:** Heatmap of pairwise G×G and G×G×E interactions among the discovered loci. For all the loci pairs, the lead SNP at *chr15_qtl* was included as a covariate to account for its large effect. LRT: Likelihood Ratio Test. **B:** Interaction among *chr11_qtl* and *chr21_qtl*. The effect of *chr21_qtl* changes direction depending on the genotype of *chr11_qtl,* but only at 28 and 35 °C. **C:** Interaction among *chr11_qtl* and *chr9_left_qtl*. The effect of *chr9_left_qtl* is suppressed by the genotype of chr11_qtl being homozygous reference, particularly at higher temperatures. **D:** The heart rate of *ccdc141* crispants (putative candidate for locus *chr21_qtl;* n = 6-15 crispants per strain; cf. **Data S5, Data S6**) compared to mock-injected control embryos (n = 7-23 embryos per strain; cf. **Data S5, Data S6**) differs depending on the temperature and the genetic background of the injected individuals (Cab and MIKK panel strains). The y-axis is log-transformed and centred on the mean of the control samples for each temperature. Data is displayed as box plots (median +/- interquartile range between the 25th and 75th percentiles) and overlaid scatter plots of individual heart rate measurements. *p*-values are reported for a general *ccdc141* editing effect on heart rate as well as within and between strains across the tested temperatures.

Having functionally validated the selected candidate genes, we next investigated whether the genetic variants identified by association mapping are present in the wild. We sampled medaka fish from natural locations in close proximity to the original MIKK panel sampling site. For the two loci with putative causal alleles giving rise to disabled genes (*ryr2b* and *ccdc141*) confirmed with equivalent gene editing, we can observe disabling variants in the wild at the same location as the F2 cross in both cases (**Table 2**). This confirmed that the mapped MIKK panel variants still occur naturally and are of ecological relevance.

### Simulations highlight the effect of model misspecification on GWAS discovery power

The pervasive presence of interaction effects in our dataset prompted us to investigate what would be the consequences of ignoring these terms in a setting akin to a human GWAS. To this end, we simulated synthetic SNPs and environmental terms with additive, dominance and interaction effect sizes estimated from our well-powered medaka genotype and phenotype data, but in the context of an unrelated outbred population. We tested generative models both with and without epistatic (G×G and G×G×E) effects. G×G effects were evaluated with respect to two loci — chr15_qtl and chr21_qtl — for which we confirmed the frequency of the causal variant in the wild. Consequently, the simulation was performed only for the remaining 14 loci. Using this synthetic dataset, we determined the consequence of performing a GWAS study using statistical models with varying degrees of mis-specification. These discovery models ranged from the original data generating model with additive genetic (G), dominance (D), gene-by-environment (G×E), dominance-by-environment (D×E), epistatic (G×G), and epistatic-by-environment (G×G×E) interactions included, to a simple additive-only SNP model with no knowledge of the environmental covariates.

In addition, we investigated the effect of uncertainty in the environmental measurement, and we simulated the commonplace scenario of not having direct access to the causal genetic variant, but only to a tagging SNP with a predetermined level of correlation to the causal variant.

For all of these conditions, we determined the minimum sample size required for the simulated locus to be discoverable at the 5* 10^-8^ significance threshold that is commonly used in human genetics (Pe’er et al., 2008). We deem a simulated locus as discoverable under a given model if the required minimum sample size is smaller than 500K samples (approximative size of large human cohorts such as the UK Biobank) for a simulated minor allele frequency smaller than 1%. Our choice of 1% allele frequency is motivated by the observation that most variants of large effect tend to be rare (Bomba et al., 2017; Yengo et al., 2022), and the QTLs we discovered with our F2 cross setup can be considered large effect variants (per-locus variance explained by the 16 QTLs 0.33% to 22.9%, c.f. maximum variance explained by a single locus 0.38%; Yengo et al., 2022). The results of our simulation are presented in **Figure 6** and summarised in **Table S3**.

**Figure 6:**
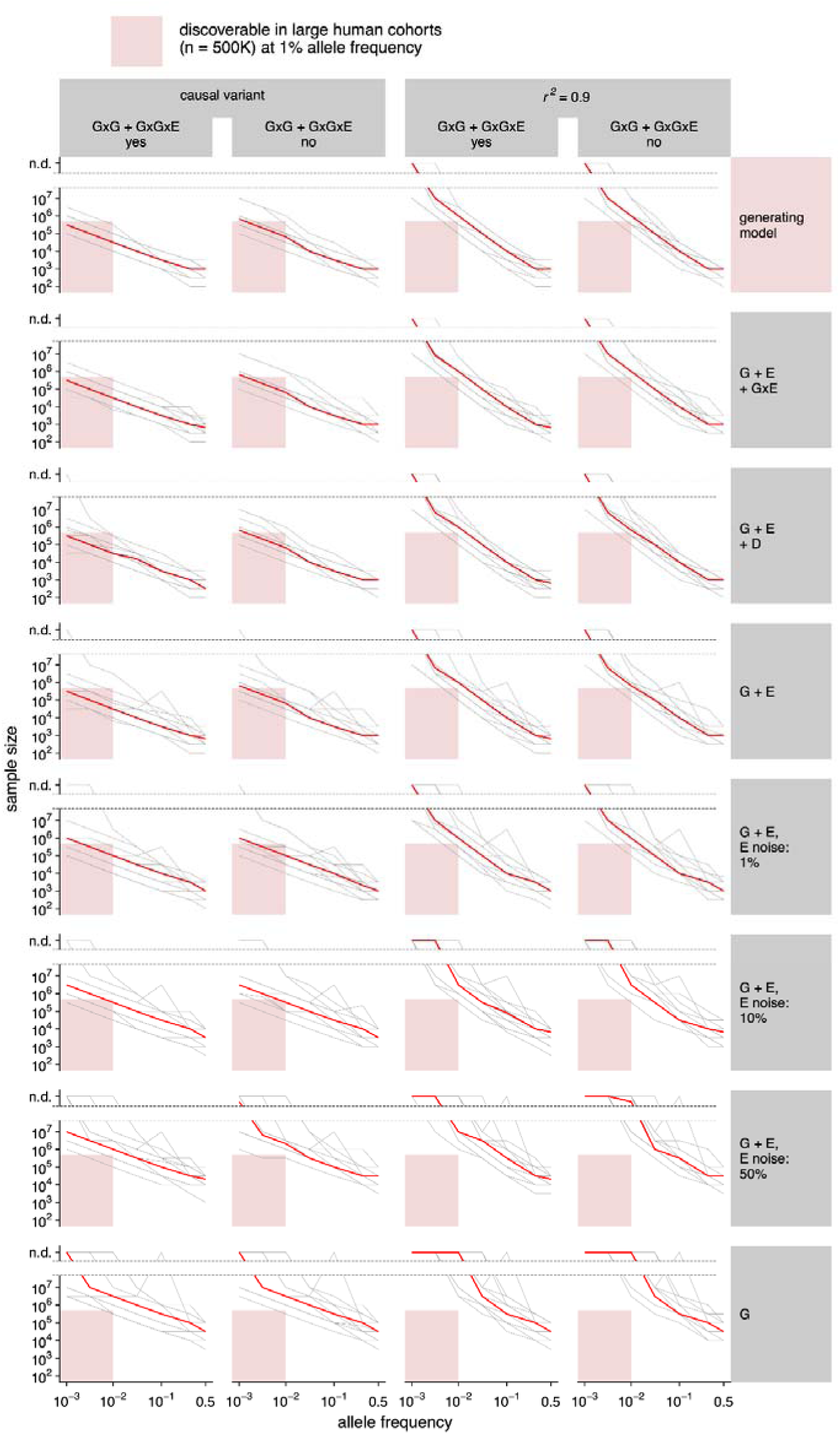
Simulation of the factors affecting discovery power in outbred GWAS and of the consequence of model mis-specification. Minimum sample size required for discovery ( ) of a locus with the same architecture (effect sizes) as the ones that we discovered in medaka, using a variety of generating models (GxG yes/no, see main text), discovery models (genetics only, linear, interactions, full model), levels of environmental (E) measurement noise (no noise, 1%, 10%, 50%), and at a range of allele frequencies (10^-3^ to 0.5) and levels of correlation strength between the genotyped and causal variant (causal variant, ). The allele frequency is expressed in terms of the most recessive allele. Each grey line represents the sample size median across 100 replicates for each medaka QTL, and the red line represents the median across the QTL medians (n = 14).

First of all, we observe that directly genotyping the causal variant is critical for discovery. Of the 14 simulated loci, only 1 is discoverable with a G + E model when the genetic variant tested has a 0.9 *r*^2^ correlation with the causal variant. On the contrary, when the discovery model has direct access to the causal variant 12 of the 14 loci are discoverable. For higher allele frequencies, such as 10%, tagging approaches would discover the majority of loci. Measuring the environment also proved to be a large determinant of discovery power in our simulations: a model with no access to the environment cannot discover any of the tested loci. Even imperfect access to the environmental covariate, in the form of an environment measurement with 1% added noise, allows for the discovery of 11 of the 14 loci.

On the contrary, we notice that interaction effects can in most cases be safely omitted from discovery modelling without an appreciable increase in sample size requirements. Using the full model that generated the simulation, which includes G×E, D, D×E, G×G, and G×G×E terms, does not allow for the recovery of any additional locus compared to a G + E + G×E model. The inclusion of D or G×E effects plays an important role in reducing sample size requirements for only 2 of the 14 loci. This effect is particularly evident for common genetic variants, while for rare alleles the benefit of including dominance terms in the discovery process is reduced. Dominant loci have a discoverability profile that is asymmetric with respect to allele frequency. That is, the required sample size is larger at a given allele frequency when the rare allele is recessive, and smaller when it is dominant (**Figure S5**). An interesting observation can be made when loci with a strong dominance component are incorrectly modelled with only additive effects. When this is the case, the resulting additive effect size estimate is dependent on allele frequency. In the extreme scenario of overdominance - which we clearly observe in 2 of the 16 medaka QTLs - this can lead to reversal of the effect direction or the complete non-discoverability of the locus. A clear example of this scenario is *chr5_qtl,* which at an allele frequency of 1% exhibits a sharp increase in required sample size. This is due to the dominance (overdominance in this case) and additive effect almost perfectly cancelling out at this specific allele frequency, leading to an estimated effect size close to 0 for an additive-only model (**Figure S6**). The G×G and G×G×E terms in the simulation, instead, do not seem to have a large effect on discoverability.

Non-additive effects, such as dominance and G×E interactions, were substantially more difficult to detect than additive effects in our simulations (**Figure S7**). Dominance was undetectable for the median locus at a 1% allele frequency, even with a sample size of 10 million. G×E interactions were highly sensitive to measurement noise in the environmental variable: with 10% environmental noise, the median locus at 1% allele frequency remained undetectable even with 10 million samples. In contrast, with perfect environmental information, the G×E effect became detectable with fewer than 1 million samples. This implies that even in ideal situations with current large human cohort sizes between 500,000 to 1 million that we should not expect to be able to discover the presence of dominance for most loci, even if present, and interactions with the environment are only detectable when the causal variant is genotyped and the environment of interest is accurately measured.

## Discussion

Our study explored the genetic architecture of heart rate in the vertebrate model medaka fish in the context of experimentally controlled temperature variation (21°C - 35°C). We leveraged 76 wild-derived inbred strains from the MIKK panel (Fitzgerald et al., 2022) to detect strain-dependent heart rate differences as well as differences in the heart rate response to temperature changes. The estimated trait heritability for the heart rate differences suggested an underlying genetic basis. By performing phenotype-genotype segregation experiments with eight founding strains we discovered 16 QTLs affecting heart rate. For 4 of these QTLs we were able to fine-map the causal gene and experimentally validate their cardiac relevance using gene editing tools and to characterise their spatial expression patterns. Out of the 16 QTLs, 8 exhibit clear interaction effects with temperatures (G×E), 4 exhibit dominance effects (D) and 2 display third order interactions between dominance and temperature (D×E). We detected 21 epistatic (G×G) interactions among the QTLs, 17 of which exhibited significant third-order G×G×E effects with temperature. Using simulations, based on our temperature-dependent heart rate QTL data set in medaka, we investigated the genetic discovery power of an additive statistical model in the presence of widespread interaction effects and showed that it still works adequately for genetic discovery. However, our simulations demonstrated that genotyping the causal variant is a key determinant of discovery power in genetic association studies and measuring the environment as precisely as possible is crucial for detecting genetic associations.

The estimated broad-sense heritability of the MIKK panel heart rate phenotypes in our study (0.42 to 0.44) aligns well with heritability estimates for physiological traits in other animal populations. For instance, it is similar to the heritability of cardiac performance traits (0.30 to 0.56) (Saha et al., 2022) or alcohol sensitivity (0.38 to 0.42) (Morozova et al., 2015) in the *Drosophila melanogaster* Genetic Reference Panel (DGRP), but is higher than recent estimates for resting heart rate heritability in humans (van de Vegte et al., 2023; Xhaard et al., 2021). This difference is likely due to the presence of non-additive effects not accounted for in human studies, as well as higher levels of confounding and non-measured environments in human cohorts. The comparison of the heart rate spectrum of the MIKK panel strains with existing heart rate measures of classical medaka strains (Gierten et al., 2025), showed that the MIKK panel harbours higher phenotypic diversity. The 16 QTLs that we discovered aided the identification and experimental validation of four genes underlying heart rate differences in wild-derived medaka (*ryr2b, ppp3cca, ccdc141, sptbn1*) which exhibit high levels of evolutionary conservation and have respective human orthologs or co-orthologs (**Figure S8**). *ryr2b* is co-orthologous to human RYR2, a classic cardiac calcium release channel which is known to play a fundamental role in heart function (Steinberg et al., 2023). Genetic variants in ryanodine receptors (RyRs) are linked to many human diseases and *RYR2* variants have been described to cause catecholaminergic polymorphic ventricular tachycardia, that can lead to ventricular arrhythmias during physical or emotional stress and even result in sudden cardiac arrest (Priori et al., 2002; Campbell et al., 2015; Sleiman et al., 2021). The channel conductance of cardiac sheep RyR has been shown to be temperature-dependent (Sitsapesan et al., 1991; Shiels & Sitsapesan, 2015). Our data shows that this also applies to heart rate of *ryr2b* variant carrier strains from the MIKK panel, with the identified temperature-sensitive *ryr2b* variants also occurring in wild-caught medaka. *ccdc141* is orthologous to human CCDC141, also called CAMDI. This gene has been shown to play a role in neuronal migration in mice (Fukuda et al., 2010) and it has been linked with heart rate variation in both humans (den Hoed et al., 2013; van den Berg et al., 2017) and medaka (Hammouda et al., 2021). *ppp3cca* is co-orthologous to the human calcineurin catalytic isozyme PPP3CC, which has been associated to blood pressure variation (Keaton et al., 2024; Kichaev et al., 2019), with calcineurin playing a key role in heart function (Parra & Rothermel, 2017). While we found *ppp3cca* specifically expressed in the habenula brain structure and the retinal pigment epithelium of medaka, the human co-ortholog is expressed in the retina and the skeletal muscle (Karlsson et al., 2021; *The Human Protein Atlas*, n.d.). In zebrafish *ppp3cca* expression was upregulated upon rewarming of hatchlings that have experienced cold stress (Ren et al., 2021). Together with our validation observations, in which a heart rate effect of *ppp3cca* crispants was only detected conditional on the temperature being above 21°C, this indicates that *ppp3cca* in fish acts in a temperature-dependent manner. Finally, *sptbn1* is co-orthologous to human SPTBN1, which has been associated with electrocardiogram features (Ntalla et al., 2020; Verweij et al., 2020), mitral valve prolapse (Roselli et al., 2022), hypoplastic left heart syndrome (HLHS) (Birker et al., 2023), and cardiovascular age (Shah et al., 2023). In HLHS probands, SPTBN1 came into focus when the oligogenic disease ethology was investigated by testing for genetic interactions (Birker et al., 2023), which we also detected for the QTL on which *sptbn1* is located in our dataset. Collectively, the analysed candidate genes validate the use of the MIKK panel and our study approach for the identification of human-relevant genetic variants associated with heart function and disease. Our experimental results showed that the phenotypic penetrance and expressivity of the edited candidate genes depended on both the temperature condition under which the phenotype was measured and the genetic background into which it was introduced. Moreover, the success of our CRISPR validation experiments is unlikely to be attributable to random chance in selecting genes with heart rate effects. In previous work, it has been shown that randomly selecting 10 genes from the medaka genome resulted in only two genes affecting heart rate upon editing with CRISPR/Cas9, and both of these genes had been linked to heart function before (Hammouda et al., 2021). Furthermore, it has been shown that embryonic heart rate in medaka can serve as an indicator of physical fitness and cardiac function in adulthood (Gierten et al., 2025), suggesting that the identified genetic variants are potentially relevant throughout life.

Our multi-parental F2 cross approach is well-suited for discovering novel associations, as it allows for achieving high allele frequencies, even for large-effect alleles that are typically rare in outbred populations. For example, the *ryr2b* variant that we discovered would have required a population size roughly six times larger to be detectable at the observed wild allele frequency of 0.03. For the *ccdc141* variant, which has an allele frequency in the wild of 0.005, the sample size would need to be 35 times larger (more than 65k samples). Predictably, fine mapping proved to be challenging in our F2 population due to the expected haplotype length. Potential strategies to improve fine mapping include integrating our results with additional genetic and phenotypic data from outbred populations, increasing sample size, and extending the number of breeding generations. Among our 16 identified QTLs, 50% exhibit significant G×E interactions, a proportion comparable to estimates in yeast for G×E effects on gene expression (Smith & Kruglyak, 2008), and higher than estimates in flies (Huang et al., 2020b). In humans, a variance QTL study on cardiac phenotypes detected G×E effects in 22% of the loci tested (Westerman et al., 2022). As for epistatic interactions, we observe significant effects in 17.5% of locus pairs, and a pervasive presence of G×G×E effects. This differs from reports in yeast, where G×G×E has been described as playing a less prominent role (Costanzo et al., 2021). To caution against equating statistical and biological interactions, we emphasize that G×E interactions can depend on the phenotypic scale and may reflect non-linear relationships between phenotype and mediator variables rather than true mechanistic interactions (Westerman & Sofer, 2024). In our study, we mitigated such artifacts by applying a variance-stabilizing transformation to the phenotype and designing the experiment to prevent hidden genotype-environment correlations. However, we cannot entirely rule out the possibility that some detected interactions arise from alternative scale related mechanisms rather than direct biological effects. Nonetheless, even scale-dependent interactions may be meaningful, particularly when they align with the primary phenotypic scale of interest (VanderWeele & Knol, 2014).

Our simulation results suggest that, as long as the causal variant is directly tested, an additive discovery model is generally appropriate for genetic association studies for rare alleles (below 1% allele frequency). This holds even in the presence of widespread interaction effects, particularly if the environment is accurately measured. However, using a proxy locus (“SNP tagging”) or measuring the environment with >10% noise significantly impacts the discovery of loci at 1% allele frequency, though loci at 10% allele frequency are more discoverable. Furthermore, we caution that in the presence of dominance, mis-specifying the model as additive can introduce implicit dependencies between effect size, allele frequency, and discoverability. The simulations identified two key determinants of statistical power: (1) reasonably accurate (<10% additional variance) measurement of environmental variables and (2) direct genotyping of causal variants through whole-genome sequencing rather than relying on tagging variants. As a consequence, we suggest prioritizing these aspects when designing GWAS cohorts, particularly in human studies, where environmental control is inherently more limited than in model organism research.

As expected from theoretical considerations and previous work, non-additive effects were notably difficult to detect in our simulations, particularly dominance, which often required sample sizes in the multiple millions at a 1% allele frequency. These findings align with previous work underscoring the limited contribution of non-additive effects to complex trait variation at the population level (Hill et al., 2008), as well as recent studies examining the role of dominance in human cohorts (Palmer et al., 2023). In our view, this study and the accompanying simulations indicate that there is no fundamental difference between human and non-human vertebrate trait architectures. While human studies have primarily identified and modelled additive effects, research in other species has leveraged controlled environments and breeding to uncover a broader range of trait-associated interactions. Although non-additive effects can be ignored for discovery or some population modelling, this does not mean they do not have important roles in individual predictions, in particular at phenotypic extremes, as is already the case for the need to consider dominance/recessive effects in many rare disease diagnoses. Furthermore, as the environment can often be a modifiable factor, discovery of GxE interactions can provide not just improved prediction but also open up other intervention pathways. Our simulation suggests though that gaining discovery power for interactions just through sample size is challenging, and other approaches may be better, for example, sampling at reasonably large-scale bottle neck populations, where stronger drift effects will increase the allele frequency of some alleles.

One of the main advantages of working with model organisms such as medaka is the ability to control both environmental and genetic variation, which is crucial in dissecting the dynamics of G×E effects. For example, in this study we leveraged the possibility to finely vary the temperature and genetically engineer medaka to demonstrate how the effect of the genetic variants affecting the *ryr2b* gene materialises only above a temperature threshold of 24 °C. More broadly, medaka sits at an optimal intermediate level of complexity: as a vertebrate, it shares key physiological and genetic similarities with humans, yet it is more cost-effective on a large scale than mammalian models. Moreover, its high tolerance to inbreeding allows for the direct establishment of inbred strains from the wild. Studies in medaka are thus complementary to those in the more closely related to human and extensively studied laboratory mouse. The MIKK panel we previously developed and this work exemplify how these advantages can be harnessed in practice. In the future, our approach to dissecting G×E and G×G interactions can be extended to more phenotypes relevant to human physiology, other environmental effects, such as exposure to chemical toxins or drugs, and to other forms of genetic variation besides small nucleotide variation.

In summary, our results lay the groundwork for future research into the genetic determinants of complex phenotypes, reinforce the value of medaka as a powerful model system, and provide guidance for the design of cohort studies that capture the full spectrum of variation — from environmental factors to genetic diversity.

## Materials and methods

### Fish maintenance

All medaka (*Oryzias latipes*) strains were maintained at Heidelberg University and Karlsruhe Institute of Technology (fish husbandry, permit numbers 35–9185.64/BH Wittbrodt, AZ35-9185.64/BH KIT) in accordance to local animal welfare standards (Tierschutzgesetz §11, Abs. 1, Nr. 1) and with European Union animal welfare guidelines (Bert et al., 2016). Fish were maintained as closed stocks in constant recirculating systems at water temperatures between 24 °C - 26 °C with a 14 hr light/10 hr dark cycle. The following medaka strains were used in this study: MIKK panel strains (Fitzgerald et al., 2022), the Cab strain and the Heino strain (Loosli et al., 2000). Both sexes were used for experiments. The fish facility is under the supervision of the local representative of the animal welfare agency.

### Wild catches

To estimate the allele frequency of the mapped genetic variants in wild medaka populations, fish were sampled 2023 in Kiyosu and in Maeshiba near Toyohashi as previously described (Spivakov et al., 2014; Fitzgerald et al., 2022). For whole genome sequencing, fin clips from 183 wild-derived medaka were collected at NIBB, Okazaki and sent to Heidelberg for extraction of genomic DNA with phenol-chloroform as described earlier (Pierotti et al., 2024). Following DNA extraction, samples were sequenced at 16.8x to 33.2x coverage at the Wellcome Trust Sanger Institute as described earlier (Fitzgerald et al., 2022).

### Automated image acquisition and heart rate detection

F0 and F2 embryos were reared in hatching medium (2 mg/l Methylene Blue Trihydrate (Sigma-Aldrich) in 1x Embryo Rearing Medium (ERM) (17 mM sodium chloride, 0.4 mM potassium chloride, 0.27 mM calcium chloride dihydrate and 0.66 mM magnesium sulphate heptahydrate at pH 7)) at 28 °C for four days and medium was changed every other day. For imaging, medaka embryos (stage 32, according to (Iwamatsu, 2004)) were rolled on sandpaper to remove the outer surface hairs of the chorion, then transferred in 150 μl 1x ERM to a 96-U bottom well plate and then sealed using a semi-permeable foil. To acquire the heart rate information at controlled temperature conditions, automated imaging was performed in a 96-well plate format using the ACQUIFER imaging machine. Embryos were exposed to temperatures ranging from 21 °C to 35 °C. After an acclimation time of at least 20 min for each tested temperature, 240 images were acquired with 24 fps. The heart rate was measured within two consecutive imaging loops per temperature condition and the heart rate average of the loops was used for further analysis. For all datasets, the heart rate information was extracted using the FEHAT detection tool (Ferreira et al., 2024). FEHAT measurements below 50 bpm were not considered because they likely result from masked heart regions during image acquisition.

Automated heart rate acquisition for the crispant and editant embryos was carried out 4 days post injection in an extended temperature ramp with seven consecutive temperature treatments (21°C, 24°C, 26°C, 28°C, 30°C, 32°C, 35°C) using the ACQUIFER imaging machine. The following amendments have been made to the previously described imaging protocol: injected embryos were not rolled on sandpaper and to prevent large temporal shifts in extended temperature experiments only one imaging loop was acquired and analysed per temperature condition.

### Crossing of phenotypic contrasting MIKK panel strains for segregation analysis

Eight MIKK panel strains with different temperature-dependent heart rate properties were selected to set up 11 different crosses (**Table S1**). The F1 generation was established from cross-mating one male with up to 5 female fish and the F2 embryos were collected from F1 families containing four to 40 individuals. In total 2667 F2 embryos were phenotyped and whole genome sequencing data were obtained for 2209 F2 embryos.

### Genomic DNA extraction and sequencing library preparation of F1 and F2 individuals

Genomic DNA extraction of F1 and F2 individuals and the sequencing library preparation was performed as previously described (Pierotti et al., 2024).

### Whole-genome sequencing and genotype imputation

The F2 medaka embryos were sequenced and their genotype was imputed with the *birneylab/stitchimpute* pipeline as previously described (Pierotti et al., 2024). Briefly, 150 bp paired-end Illumina short-read sequencing was performed on a NextSeq2000 machine (https://www.illumina.com/). We sequenced a variable number of samples per flow cell, achieving an average sequencing depth of 1.4x. Reads were aligned to the Hdr-R medaka reference genome (ENSEMBL ID: ASM223467v1) using the *nf-core/sarek* pipeline (Hanssen et al., 2023). As an aligner we used *bwa-mem2* (Vasimuddin et al., 2019) and reads were deduplicated with *GATK MarkDuplicates* (Van der Auwera & O’Connor, 2020).

One F1 sample per cross and 2 F2 samples from the 72-2 x 55-2 cross were sequenced at higher depth (12 samples in total, obtaining a sequencing depth of 33x to 61x). These samples were used as a ground truth for validating the imputation process. These samples were also processed with the *nf-core/sarek* pipeline. As an aligner we used *bwa-mem2* and reads were deduplicated with *GATK MarkDuplicates*. Genotypes were obtained with *GATK via* joint germline variant calling (Poplin et al., 2018).

We ran the genotype imputation on the low-coverage F2 samples and high coverage F1 and F2 samples jointly. The high coverage samples were downsampled to a sequencing depth of 0.5x before being used for the imputation. Imputation accuracy was evaluated as the squared Pearson correlation (*r*^2^) between the imputed genotypes of the downsampled high-coverage samples and the genotype calls obtained from the same samples with GATK.

The initial set of genetic variants used for the imputation process was obtained from the variants called by GATK on the high-coverage samples, and consisted of 6.2 million SNPs. We filtered this variant set with `bcftools` `view` (Danecek et al., 2021) using the following filtering criteria: `-i ‘CHROM!=“MT”’, -i ‘TYPE==“snp”’, -i ‘N_ALT >= 1’, -i ‘MAC >= 1’`. This variant set was then refined iteratively using the “`snp_set_refinement`” mode of the *birneylab/stitchimpute* pipeline. We performed 5 iterations and at each iteration retained only the SNPs that passed a certain *r*^2^ threshold. We used the following filter values: 0.5, 0.5, 0.75, 0.9, and 0.9. Our final set of SNPs consisted of 3.2 million variants.

Imputation parameters K and nGen were optimised using the “`grid_search`” mode of the *birneylab/stitchimpute* pipeline. We selected `K=16` and `nGen=2`. In addition, we used the following parameters: `expRate⍰=⍰2, niterations⍰=⍰100, shuffleHaplotypeIterations⍰=⍰seq(4, 88, 4), refillIterations⍰=⍰c(6, 10,14, 18), shuffle_bin_radius =⍰1000`.

Overall, the imputed genotypes proved to be very reliable, with and average sample-wise *r*^2^ value of 0.996 between the genotypes called on the high coverage samples using GATK and the imputation obtained from the same samples with reads downsampled to 0.5x depth.

### Linear mixed model genetic association analysis

The association between genetic variants and the heart rate phenotype was analysed using a mixed linear model, implemented in the *birneylab/flexlmm* pipeline (Pierotti et al., 2025). The association analysis was run separately for the heart rate measurements taken at different temperatures. Additional phenotypes consisting of the heart rate difference in the same samples across temperatures were also used in the association analysis. All the phenotypes were quantile normalised to a normal distribution. We used a random effect proportional to the genetic relatedness matrix of the samples, and fixed covariates corresponding to intercept, phenotyping plate ID and F2 cross ID. Significance was evaluated with a likelihood ratio test between a covariate-only model and a model that in addition to the covariates contained a term for the linear encoding of the genetic variant (homozygous reference encoded as 0, heterozygous encoded as 1, homozygous alternate encoded as 2) and a dominance term (heterozygous encoded as 1, any homozygous encoded as 0). Genome-wide significance thresholds were established as the minimum *p*-value obtained across all the SNPs tested over 100 permutations of the residuals (see (Pierotti et al., 2025) for methodological details).

### Locus definition and fine mapping

The boundaries of the association regions were defined manually by visual inspection of the association profile and linkage disequilibrium structure to the lead SNP. The lead SNP was usually taken to be the variant with the smallest *p*-value in the locus, but excluding variants missing completely one homozygous state, or with impossible segregation patterns across the different F2 crosses.

### Candidate gene selection

Candidate genes (**Table S2**) for gene editing validation were selected within the association regions of each locus according to the presence of variants disrupting protein function that are also strongly linked to the lead variant for the locus, and previously reported cardiac effects associated with the gene. To check for previously reported cardiovascular effects for a gene, we used the phenotype tab of each gene’s page on the ENSEMBL genome browser (Cunningham et al., 2021). We also performed manual literature searches. To detect variants with a consequence on protein function that are in strong linkage with a lead variant for a locus, we used the *birneylab/varexplore* pipeline that we describe in the next section.

### The *birneylab/varexplore* pipeline

We reasoned that the causal mutation for an observed association signal may not be included in the set of genetic variants used for association testing. This may be because, for example, the causal variant is an insertion or deletion, while the set of variants that we used in the association testing were exclusively SNPs (due to limitations of the STITCH imputation software used).

To address this issue and detect putative causal variants that are not part of the tested set, we developed the *birneylab/varexplore* (https://github.com/birneylab/varexplore) Nextflow (Di Tommaso et al., 2017) pipeline, which we also make publicly available with associated documentation for other researchers to use. *birneylab/varexplore* can take advantage of linkage disequilibrium around a variant of interest to infer the presence of additional linked genetic variants. It requires in input aligned (low-depth) sequencing reads for a set of samples, a file containing a set of variants of interest (for example lead SNPs from a GWAS analysis), and called genotypes for the same samples in *vcf* format (for example deriving from an imputation process). The pipeline extracts genotypes from the *vcf* file at the variants of interest, and clusters the samples according to their genotype at such variants (this is performed in parallel as separate processes for each variant). The sequencing reads are clumped together using *samtools* (Danecek et al., 2021) according to the cluster to which each sample belongs (again, in parallel different clusterings are performed for different variants), and processed as meta-samples in downstream steps. These meta-sample reads are processed with *GATK* performing germline joint calling (Van der Auwera & O’Connor, 2020). Variants in strong linkage disequilibrium with the grouping variants used, which may have been absent in the variant set used for the original GWAS, are expected to be called with different genotypes in the different meta-samples. On the contrary, variants that are not linked to the grouping variant will appear as heterozygous in all the meta-samples. The effects of the newly discovered variants are then predicted using ENSEMBL Variant Effect Predictor (VEP) (McLaren et al., 2016), and the output is reformatted so that the variants and their effects can be directly loaded to the Integrative Genomics Viewer (IGV) (Robinson et al., 2011) for manual exploration. Other outputs produced by the pipeline for further analysis are the meta-sample grouped reads in cram format, the meta-sample variant calls in *vcf* format, and the ENSEMBL VEP predictions.

To aid in filtering out variants that are not linked to the variants of interest, we also provide an R script (R Core Team, 2023), `filter_variants.R,` within the pipeline repository that can parse the various outputs of the pipeline, and retain only variants that follow specific criteria in terms of genotype distribution across the meta-samples. For example, using this script it is possible to retain only variants that are called as opposite homozygous genotypes for the meta-sample corresponding to opposite homozygous genotypes at the grouping variant, and heterozygous for the meta-sample heterozygous at the grouping variant. More complex configurations are also possible.

For a more complete description of the pipeline usage and capabilities, the reader is encouraged to explore the *birneylab/varexplore* documentation at https://github.com/birneylab/varexplore.

### gRNA design

The gRNA target site was selected in close proximity to the mapped genetic variant and the gRNAs were designed as previously described with the target prediction tools CCTop (Stemmer et al., 2017) and ACEofBAses (Cornean et al., 2022). Locus-specific crRNAs and the tracrRNA backbone were ordered via the Integrated DNA Technologies (IDT) synthesis service. The crRNA and the tracrRNA were both diluted in nuclease-free duplex buffer (IDT) to a final concentration of 40 µM and incubated at 95°C for 5 min to form a functional gRNA duplex (**Table S2**).

### *In vitro* transcription of mRNA

To generate the Cas9 mRNA and evoBE4max mRNA, pCS2+ (heiCas9) (Thumberger et al., 2022) and pCS2+ (evoBE4max) (Cornean et al., 2022) were linearized by a NotI digest and the mRNA was transcribed using the mMESSAGE mMACHINE SP6 or T7 Transcription Kit (Thermo Fisher Scientific). RNA purification was performed with the RNeasy Mini Kit (Qiagen) according to manufacturer’s instructions.

### Microinjections

For gene-editing, microinjections were performed at the 1-cell stage. The injection solution contained 150 ng/µl editor mRNA (either Cas9 or evoBE4max), 4 pmol gRNA and 10 ng/µl GFP mRNA as injection tracer. As a mock-control 10 ng/µl GFP mRNA was injected. Injected embryos were kept at 28°C in medaka embryo rearing medium (ERM, 17 mM NaCl, 40 mM KCl, 0.27 mM CaCl2, 0.66 mM MgSO4, 17 mM Hepes) and selected for GFP signal 7 hours post injection. Phenotyping was done 4 days post fertilization.

### Genotyping of crispants and editants

Single embryos were lysed in DNA extraction buffer (0.4 M Tris/HCl pH 8.0, 0.15 M NaCl, 0.1% SDS, 5 mM EDTA pH 8.0; 1 mg/ml proteinase K) at 60°C overnight. Nuclease-free water was added 1:2 and the proteinase K was inactivated for 20 min at 95°C. To precipitate the genomic DNA, 300 mM sodium acetate and 3x vol. absolute ethanol were added, followed by centrifugation at 20,000 x g at 4°C. The precipitated DNA was resuspended in TE buffer (10 mM Tris pH 8.0, 1 mM EDTA in RNAse-free water) and 1 µl used as input material for the genotyping-PCR reaction: 1 x Q5 reaction buffer, 200 µM dNTPs, 200 µM forward and reverse primer (for locus-specific primer pairs see **Table S2**) and 0.3 U Q5 polymerase (NEB). The following PCR cycling conditions were used: 2 min at 98°C, 30 cycles of 30 sec at 98°C, primer annealing for 20 - 30 sec at 60 - 70°C, 30 sec at 72°C, and a final extension of 5 min at 72 °C. PCR products were purified via agarose gel electrophoresis and extracted using the Monarch DNA Gel Extraction Kit (NEB) before forwarding to Sanger sequencing (Eurofins Genomics). Sanger sequencing results were visualized and analysed using Geneious Prime (2019.2.3, BioMatters).

### Microscopy

*In vivo* images of phenotypic-representative embryos were acquired 4 days post-injection with a Nikon SMZ18 equipped with Nikon DS-Ri1 and DS-Fi2 cameras. For imaging embryos were mounted into injection molds (1.5 % (w/v) agarose in ERM). All images were processed using ImageJ (Schindelin et al., 2012). ImageJ was used for image cropping and to adjust brightness and contrast settings.

### Wholemount *in situ* hybridisation

Wholemount in *situ* hybridisation using BCIP-NBT as chromogenic substrates was performed as previously described (Loosli et al., 1998). Stained embryos were mounted in 87% Glycerol on a microscope slide and imaged in a brightfield microscope (Nikon Eclipse 80i microscope equipped with a pco.panda 4.2 camera). The RNA probes were synthesized from plasmids that derived from an in-house cDNA library (Souren et al., 2009) or were ordered from Biocat and transcribed with the T7 polymerase.

### Detection of G×E and G×G effects

To detect the effect size and statistical significance of interaction terms, additional Nextflow pipelines were set up. These are available at https://github.com/birneylab/heart_rate_tmp_gxe and https://github.com/birneylab/heart_rate_tmp_pairwise_gxg. We performed both analyses uniquely on the marker SNPs for each locus. Differently than for the discovery GWAS, in this case we run the analysis jointly on the heart rate measured at different temperatures. The lead SNP of the *chr15_qtl* locus and its interaction with the temperature were included as a covariate in all the cases where *chr15_qtl* was not the target locus itself.

For the G×E analysis, we used a linear mixed model with 3 random effects covariance matrices. One covariance matrix was designed to correspond to a binary specification of the sample identity (1 for pairs of measurements relating to the same sample and 0 otherwise, to account for the repeated measurements of the same sample across temperatures). The second covariance matrix was derived from the genetic relatedness matrix and expanded to account for the repeated presence of the same samples at 3 different temperatures (it was computed as the Kronecker product of the GRM and a 3x3 matrix full of 1s). This second matrix modelled the relationship between samples across different temperatures as well as within the same temperature, and thus served to estimate a genome-wide temperature-independent covariance. The third matrix was computed as the Kronecker product of the GRM and a 3x3 identity matrix. This last matrix served to encode the relatedness among samples within the same temperature, while the covariance across measurements taken at different temperatures was always set to 0. Thus, it estimated genetically-determined covariance among samples within but not between temperature treatments. Together, the second and third matrix define a compound symmetry environment model where the variance and covariance among environments are constant (but possibly different from each other). The same fixed effects covariates used in the discovery GWAS were adopted. The GRMs were computed in a Leave-One-Chromosome-Out (LOCO) fashion, excluding the chromosome where the currently tested locus resided when calculating pairwise sample relatedness. The variance components were estimated using the *gaston* R package (Claire & Hervé, 2018). The variance components were estimated once per locus, using a fixed effect structure containing only an additive encoding for the locus and no dominance or G×E terms. The temperature was treated as a categorical variable, while a log-transformation of the heart rate was used as a phenotype for stabilising the residual variance. Phenotyping batch and F2 cross were used as fixed-effect covariates. After estimation of the variance components, the random effects were regressed out in the same way described in (Pierotti et al., 2025). Linear models were then fitted to the decorrelated data to estimate effect sizes. To assess statistical significance of interaction terms and dominance (**Figure 3C**), likelihood ratio tests were performed. The *p*-values were then Bonferroni-corrected and assessed at a 0.05 threshold. To calculate the variance explained by the additive genetic (G), dominance (D), gene-by-environment (G×E), and dominance-by-environment (D×E) terms (**Figure 3D**), an ANOVA model (including covariates) was fitted on the decorrelated data. The sum of squares for each term was then divided by the total sum of squares for the model. To calculate the variance explained as a proportion of the overall variance explained by genetic terms (**Figure 3E**), the sum of squares for each term was divided by the total sum of squares for the G, D, G×E, and D×E terms.

The G×G analysis (**Figure 5A**) was performed similarly to the G×E analysis, using the same structure for the random effects. In addition to the covariates used in the G×E analysis, the marginal effect of the interacting loci tested and their marginal G×E effect were included as covariates. The GRMs were computed excluding 2 chromosomes at a time (for the 2 interacting loci modelled). Significance was determined via likelihood ratio tests and Bonferroni-corrected to account for the number of tests performed (120).

### Calculation of variance explained by single loci

The variance explained estimates reported in **Figure 3D**, **Figure 3E** and **Data S7** were derived from the locus-specific linear model fits used in the G×E analysis described in the previous section. For each locus, variance explained was calculated as the partial *η*^2^ for the genetic terms—defined as the ratio of the sum of squares attributable to the genetic effects (including interactions) to the residual sum of squares from the ANOVA.

### Statistical analysis of gene editing results

Gene editing validation results (**Figure 4B**) were modelled with a mixed effect model using the *lmerTest* R package (Kuznetsova et al., 2017). Random intercepts and random slopes with respect to temperature were included to account for repeated measurements of the same samples across different temperatures. The heart rate phenotype was log-transformed to stabilise residual variance, and temperature was also log-transformed to linearise its relationship with the transformed heart rate (as in the simulations described in the next section, we used *log(t - 6)*, where *t* is the temperature in degrees Celsius). All *p*-values were FDR-corrected.

For plotting only (but not for statistical analysis), phenotypes were mean-centred based on the mean of the control samples, and temperature was shown on the original scale.

Data points were considered strong outliers and excluded from the analysis following Tukey’s definition (Tukey, 1977) if they fell outside the range:

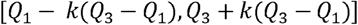

With *Q*_1_ and *Q*_3_ the first and third quartiles respectively. The value of *k* was conservatively set to 4, marking only exceptionally strong outliers (by comparison, *k*□=□1.5 defines standard Tukey outliers, and *k*□=□3 marks points that are “far out”).

The data presented in **Figure 5D** were analysed similarly. Separate mixed models were fit, stratified by background medaka strain (with results shown in each subplot), and the resulting *p*-values were FDR-corrected. A joint mixed model was also fit to the data, treating strain and its interaction with gene editing status as categorical fixed effects (results shown in the top right of Figure 5D).

### Simulation of the effects of model misspecification

To assess the detectability of QTLs with effect sizes similar to those observed in the medaka dataset under varying levels of model mis-specification, we developed a simulation framework. The corresponding simulation code is available as a Nextflow pipeline at: https://github.com/birneylab/heart_rate_tmp_simulation_gxg.

For each medaka QTL, we simulated the same locus at a range of allele frequencies and sample sizes, with 100 replicates per combination. QTL genotypes were generated by sampling from a binomial distribution with two trials and success probability equal to the target allele frequency. Environmental data (temperature) were sampled with replacement from outdoor water temperature measurements collected in the MIKK panel’s native geographic region in Japan (courtesy of Dr. Nakayama, Nagoya University and used also in (Nakayama et al., 2023)). These measurements were recorded from the middle layer of a 60L water container every hour over two years (1 October 2015 – 15 October 2017). We restricted our sampling to values between 21–35□°C during the medaka reproductive season, avoiding extrapolation beyond the range directly observed in the GWAS.

With the exception of *chr15_qtl* and *chr21_qtl* (discussed below), we fitted a linear mixed model to the medaka GWAS data for each locus, using the same model framework described previously in the context of estimating G×E and dominance effects. Effect sizes were estimated under two models: a G + E + D + G×E + D×E model, and an extended model that also included G×G and G×G×E terms (labelled “GxG + GxGxE no” and “GxG + GxGxE yes”, respectively, in **Figure 6**). G×G and G×G×E interactions were estimated between each focal locus and the lead SNPs at *chr15_qtl* and *chr21_qtl*, for which causal variants had been identified and their frequencies in the wild determined. As a consequence, the *chr15_qtl* and *chr21_qtl* loci themselves were not used as focal loci in simulations. The *chr15_qtl* and *chr21_qtl* loci were always simulated at their wild allele frequency under a binomial distribution.

The marginal effect of *chr15_qtl* and *chr21_qtl* and their interaction with temperature were always included as a fixed effect covariates, both in the “GxG and GxGxE yes” and “GxG and GxGxE no” models.

Unlike the approach used for G×E and dominance effect detection described earlier, here we treated temperature as a continuous variable to support simulations across a broader, realistic range of values—not just 21, 28, and 35□°C. We log-transformed the heart rate phenotype to stabilise residual variance. To linearise the relationship between temperature and log-transformed heart rate in the 21–35□°C range, we also applied a log-based transformation to the temperature (we used *log(t - 6)* where *t* is the temperature in degrees Celsius). These transformations were consistently applied to both the medaka data (for effect size estimation) and the simulated data (for discovery).

Synthetic phenotypes were generated by summing the products of the simulated values of each term (intercept, G, E, D, G×E, D×E, G×G, G×G×E) and the effect sizes estimated from medaka. For the “GxG and GxGxE no” column in **Figure 6** the epistatic terms were omitted. Gaussian noise with variance equal to the residual variance of the corresponding medaka locus was then added. We did not include covariates or random effects in the simulations, as these were specific to our experimental setup and population.

We conducted QTL discovery on the simulated data using models of increasing complexity. In each case, we performed a likelihood ratio test comparing the full model to a reduced model that included only temperature, intercept, *chr15_qtl* and *chr21_qtl*, and their interactions with temperature. In the case of the G-only model (without environment), the comparison was made against a model with just the intercept, *chr15_qtl* and *chr21_qtl*. To evaluate the impact of environmental noise, we introduced Gaussian noise to the temperature variable such that noise accounted for 1%, 10%, or 50% of the total temperature variance. The “generating model” row in **Figure 6** refers to simulations evaluated under models that either included (for the “GxG and GxGxE yes” column) or excluded (for the “GxG and GxGxE no” column) the G×G and G×G×E terms.

We also evaluated the effect of using tagging rather than causal SNPs by replacing the causal variant used to generate the phenotype with a SNP with known linkage (e.g., *r²* = 0.9) to the causal variant, and recomputing all interaction terms accordingly (**Figure 6**, “causal variant” vs “*r²* = 0.9” columns).

Each simulation condition (combination of QTL, sample size, allele frequency, epistasis model, and tagging level) was repeated 100 times. For each replicate, we identified the minimum sample size at which genome-wide significance (*p*□<□5×10□□) was achieved. Median minimum sample sizes across replicates are shown in grey in **Figure 6**, with the median across QTLs plotted as a red line.

In Figure S5, we present QTLs ordered by their degree of non-additivity. We quantified non-additivity using a dominance ratio:

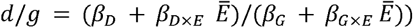

where *β*_*G*_, *β*_*G×E*_, *β*_*D*_, and *β*_*D×E*_ are the effect sizes for a given QTL, and *Ē* is the mean environmental value. This analysis used estimates from the "GxG and GxGxE no" models. The absolute value of the dominance ratio represents the degree of non-additivity; loci with a positive ratio are classified as dominant, and those with a negative ratio as recessive. In **Figure S5**, loci are colour-coded according to dominance/recessiveness. To visualise the reciprocal condition, we also plot each locus with the reference and alternate alleles swapped.

In **Figure S7** we instead show the sample size required to discover the non-additive effects, specifically dominance (D) and GxE interactions. In this analysis, likelihood ratio tests were conducted against a null model that included the additive effect of the locus, the environment, *chr15_qtl*, *chr21_qtl*, and the intercept.

### Construction of gene trees for confirmed genes

The trees shown in **Figure S8** were built with *iqtree2* (Minh et al., 2020) based on protein multiple sequence alignment performed with *t-coffee* (Notredame et al., 2000), both run with default parameters. Visualisations were produced with *iTOL* (Letunic & Bork, 2024), and homologues were retrieved from *OrthoDB* (Tegenfeldt et al., 2025). The set of species included in each tree varied depending on the availability of annotated homologues, but human, mouse, chicken, and medaka sequences were always included. The brownbanded bamboo shark (*Chiloscyllium punctatum*) or the roundworm (*Caenorhabditis elegans*) was used as an outgroup to root each tree. Very short or poorly aligned sequences were excluded from the analysis.

### Statistical tests for reciprocal cross effect

To test whether the maternal versus paternal origin of the founding strain influenced QTL effects, we focused on reciprocal F2 crosses (72-2×139-4 and 139-4×72-2; *n*□=□302), in which the parental strains were swapped in maternal and paternal roles. We further restricted the analysis to QTLs for which each reciprocal cross included at least 10 samples in the least frequent genotype. Analyses were stratified by temperature treatment and QTL, using an inverse quantile normalisation of the heart rate phenotype. For each combination, we fitted a null linear model including phenotyping plate as a covariate, as well as the main effects of QTL and reciprocal cross. This was compared via Likelihood Ratio Tests (LRTs) to an alternative model that additionally included the QTL × reciprocal cross interaction. Resulting *p*-values were corrected using both FDR and Bonferroni adjustments (**Data S9**).

Separately, we tested for a marginal effect of reciprocal cross status by comparing a model with only phenotyping plate as a covariate to one that included both plate and reciprocal cross (without any genetic terms). As before, significance was assessed using LRTs and corrected with a Bonferroni correction.

## Acknowledgments

We would like to thank all members of the Birney and Wittbrodt research group for their valuable input to the project and the manuscript. We thank for Tinatini Tavhelidse-Suck for her contribution to the COS-MIKK propagation campaign and Rachel Müller, Tanja Kellner, Beate Wittbrodt for expert technical assistance, Rie Ajioka, Yukari Koike, Yuko Teshima, Nadeshda Wolf, Natalja Kusminski, Thomas Seitz, Marzena Liv, Erik Leist and Antonino Sarazeno for animal husbandry. The authors thank Kouko Yamazaki for sampling wild medaka. We are indebted to Jochen Gehrig and Laurent Thomas (ACQUIFER, Bruker) for their technical advice on high-throughput imaging. The authors acknowledge Vladimir Benes, Ferris Jung, Mireia Osuna-López, and all members of the EMBL Genomics Core Facility (GeneCore) as well as the Wellcome Trust Sanger Institute for their support in the library preparation and the whole-genome sequencing. We would like to thank the National Bioresource Project (NBRP) medaka Japan and NIBB Individual collaborative research projects (24NIBB313) for the access to the wild-caught medaka fish.

This research was supported by the European Research Council Synergy Grant IndiGene (number 810172) and the NIH (National Institutes of Health, number R01ES029917).

## Supplementary Tables

**Table S1:**
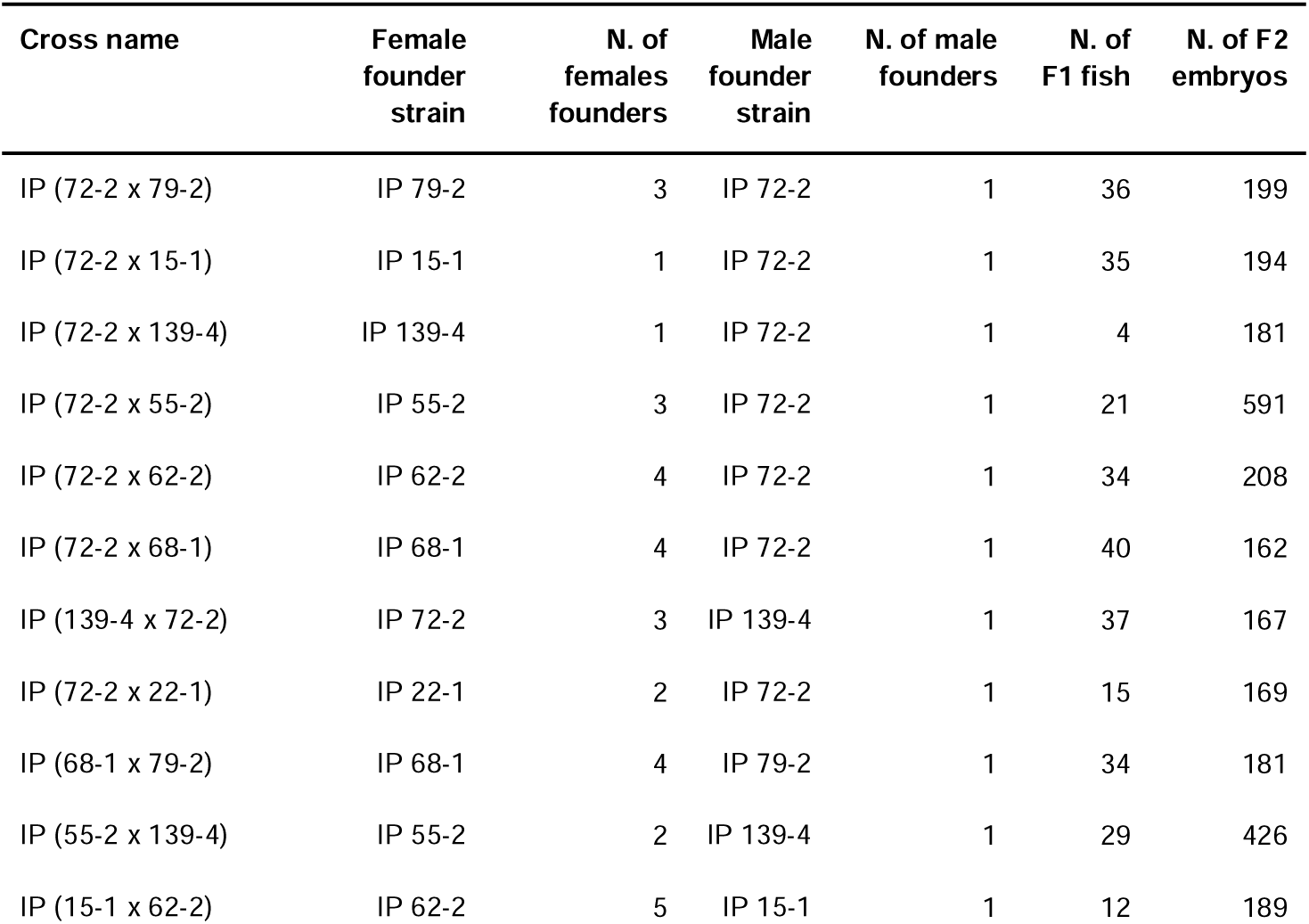
Overview of the MIKK panel strain crosses used for segregation analysis. Number (N.) of cross founder individuals, F1 mating groups sizes and number of F2 embryos heart rate phenotyped per cross. IP: Inbred Panel.

| Cross name | Female founder strain | N. of females founders | Male founder strain | N. of male founders | N. of F1 fish | N. of F2 embryos |
| --- | --- | --- | --- | --- | --- | --- |
| IP (72-2 x 79-2) | IP 79-2 | 3 | IP 72-2 | 1 | 36 | 199 |
| IP (72-2 x 15-1) | IP 15-1 | 1 | IP 72-2 | 1 | 35 | 194 |
| IP (72-2 x 139-4) | IP 139-4 | 1 | IP 72-2 | 1 | 4 | 181 |
| IP (72-2 x 55-2) | IP 55-2 | 3 | IP 72-2 | 1 | 21 | 591 |
| IP (72-2 x 62-2) | IP 62-2 | 4 | IP 72-2 | 1 | 34 | 208 |
| IP (72-2 x 68-1) | IP 68-1 | 4 | IP 72-2 | 1 | 40 | 162 |
| IP (139-4 x 72-2) | IP 72-2 | 3 | IP 139-4 | 1 | 37 | 167 |
| IP (72-2 x 22-1) | IP 22-1 | 2 | IP 72-2 | 1 | 15 | 169 |
| IP (68-1 x 79-2) | IP 68-1 | 4 | IP 79-2 | 1 | 34 | 181 |
| IP (55-2 x 139-4) | IP 55-2 | 2 | IP 139-4 | 1 | 29 | 426 |
| IP (15-1 x 62-2) | IP 62-2 | 5 | IP 15-1 | 1 | 12 | 189 |

**Table S2:**
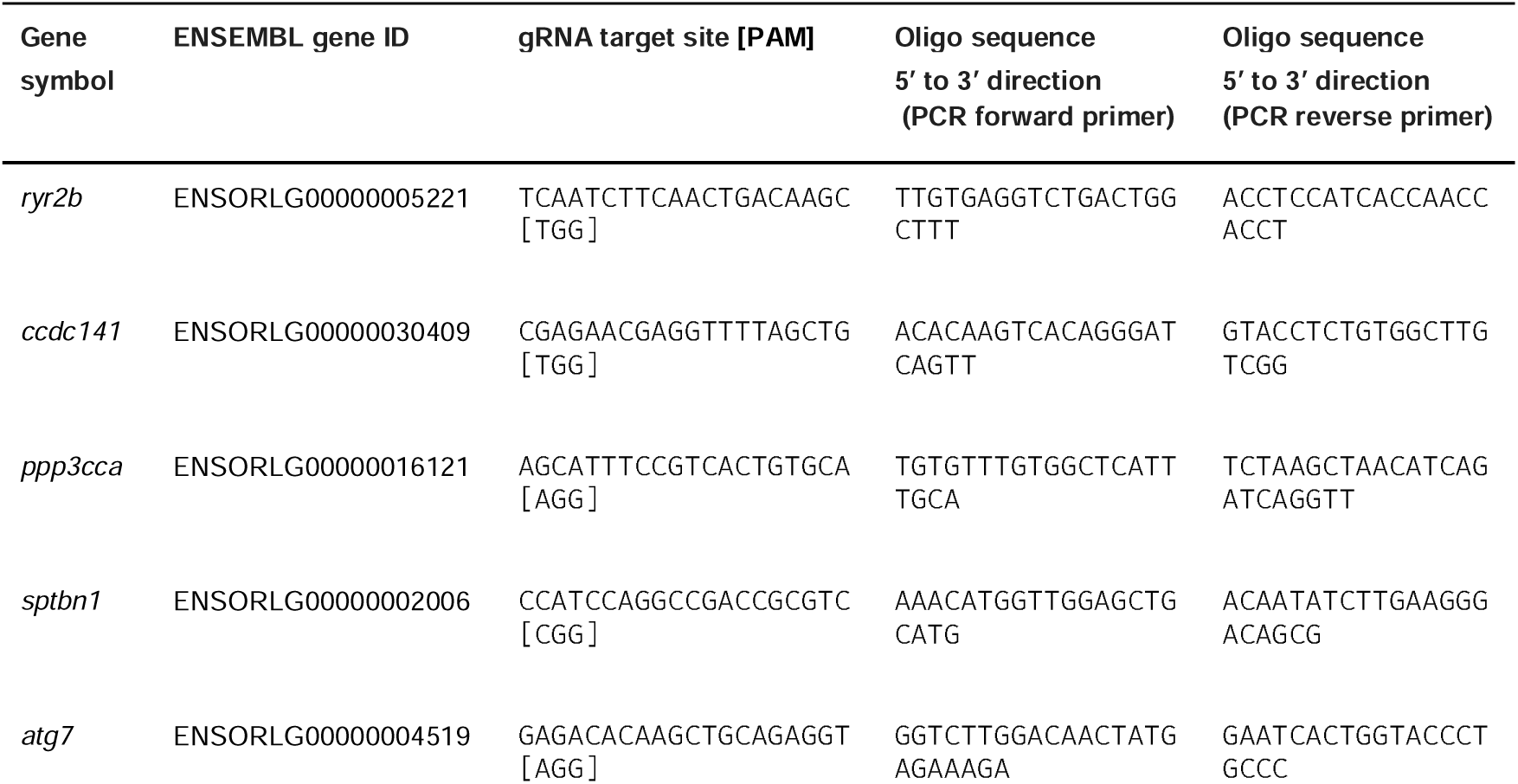
Locus-specific CRISPR/Cas9 and base editor target sites with the PAM of the gRNA sequence in brackets and oligonucleotides used for target site amplification via PCR.

**Table S3.**
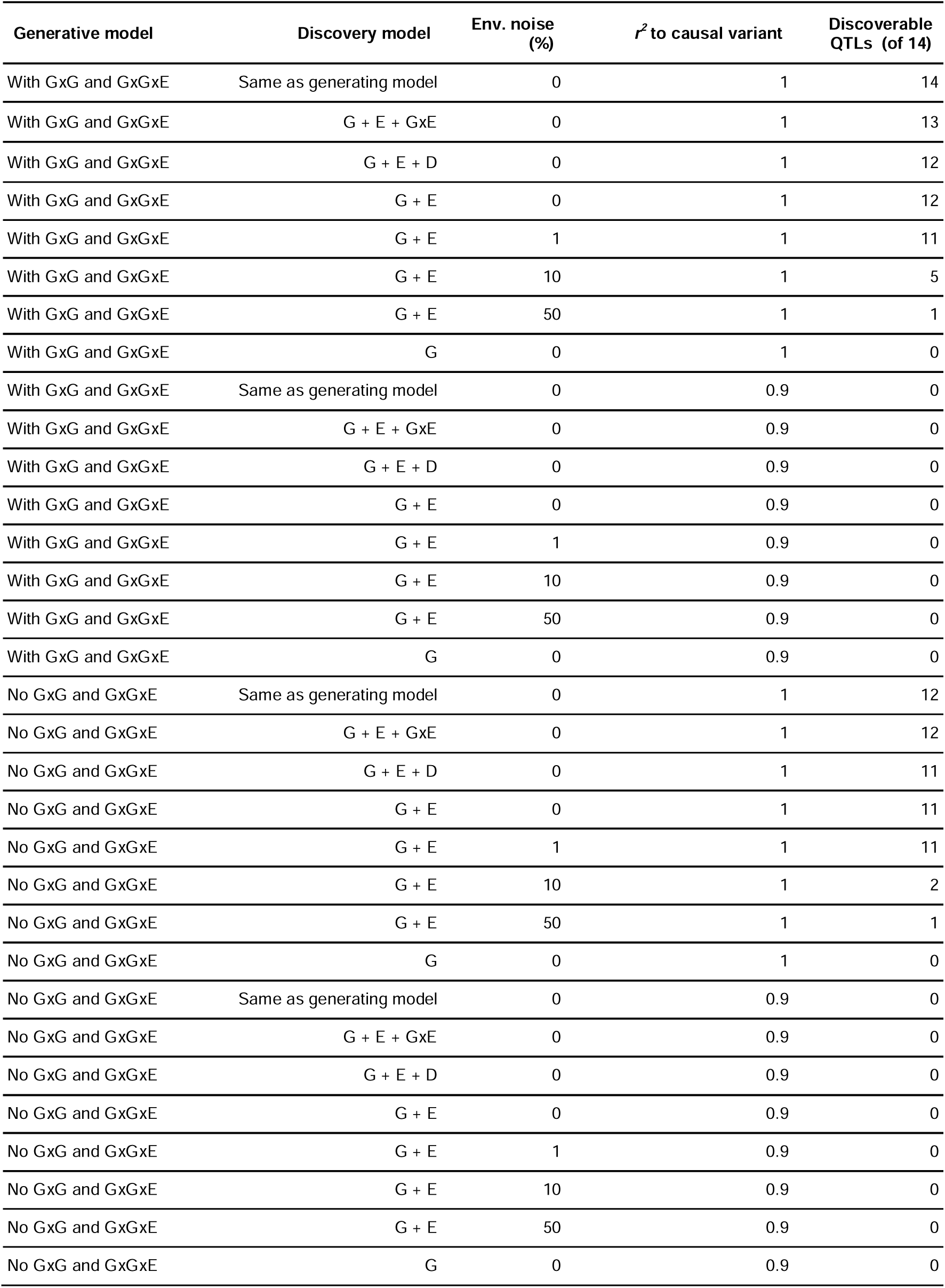
Simulation summary. Summary of the simulation results described in **Figure 6**. A QTL is deemed discoverable if the minimum sample size required to discover it at p < 5 * 10^-8^ is smaller than 500K for an allele frequency smaller than 1%.

| Generative model | Discovery model | Env. noise (%) | $r^2$ to causal variant | Discoverable QTLs (of 14) |
| --- | --- | --- | --- | --- |
| With GxG and GxGxE | Same as generating model | 0 | 1 | 14 |
| With GxG and GxGxE | G + E + GxE | 0 | 1 | 13 |
| With GxG and GxGxE | G + E + D | 0 | 1 | 12 |
| With GxG and GxGxE | G + E | 0 | 1 | 12 |
| With GxG and GxGxE | G + E | 1 | 1 | 11 |
| With GxG and GxGxE | G + E | 10 | 1 | 5 |
| With GxG and GxGxE | G + E | 50 | 1 | 1 |
| With GxG and GxGxE | G | 0 | 1 | 0 |
| With GxG and GxGxE | Same as generating model | 0 | 0.9 | 0 |
| With GxG and GxGxE | G + E + GxE | 0 | 0.9 | 0 |
| With GxG and GxGxE | G + E + D | 0 | 0.9 | 0 |
| With GxG and GxGxE | G + E | 0 | 0.9 | 0 |
| With GxG and GxGxE | G + E | 1 | 0.9 | 0 |
| With GxG and GxGxE | G + E | 10 | 0.9 | 0 |
| With GxG and GxGxE | G + E | 50 | 0.9 | 0 |
| With GxG and GxGxE | G | 0 | 0.9 | 0 |
| No GxG and GxGxE | Same as generating model | 0 | 1 | 12 |
| No GxG and GxGxE | G + E + GxE | 0 | 1 | 12 |
| No GxG and GxGxE | G + E + D | 0 | 1 | 11 |
| No GxG and GxGxE | G + E | 0 | 1 | 11 |
| No GxG and GxGxE | G + E | 1 | 1 | 11 |
| No GxG and GxGxE | G + E | 10 | 1 | 2 |
| No GxG and GxGxE | G + E | 50 | 1 | 1 |
| No GxG and GxGxE | G | 0 | 1 | 0 |
| No GxG and GxGxE | Same as generating model | 0 | 0.9 | 0 |
| No GxG and GxGxE | G + E + GxE | 0 | 0.9 | 0 |
| No GxG and GxGxE | G + E + D | 0 | 0.9 | 0 |
| No GxG and GxGxE | G + E | 0 | 0.9 | 0 |
| No GxG and GxGxE | G + E | 1 | 0.9 | 0 |
| No GxG and GxGxE | G + E | 10 | 0.9 | 0 |
| No GxG and GxGxE | G + E | 50 | 0.9 | 0 |
| No GxG and GxGxE | G | 0 | 0.9 | 0 |

## Supplementary Figures

**Figure S1.**
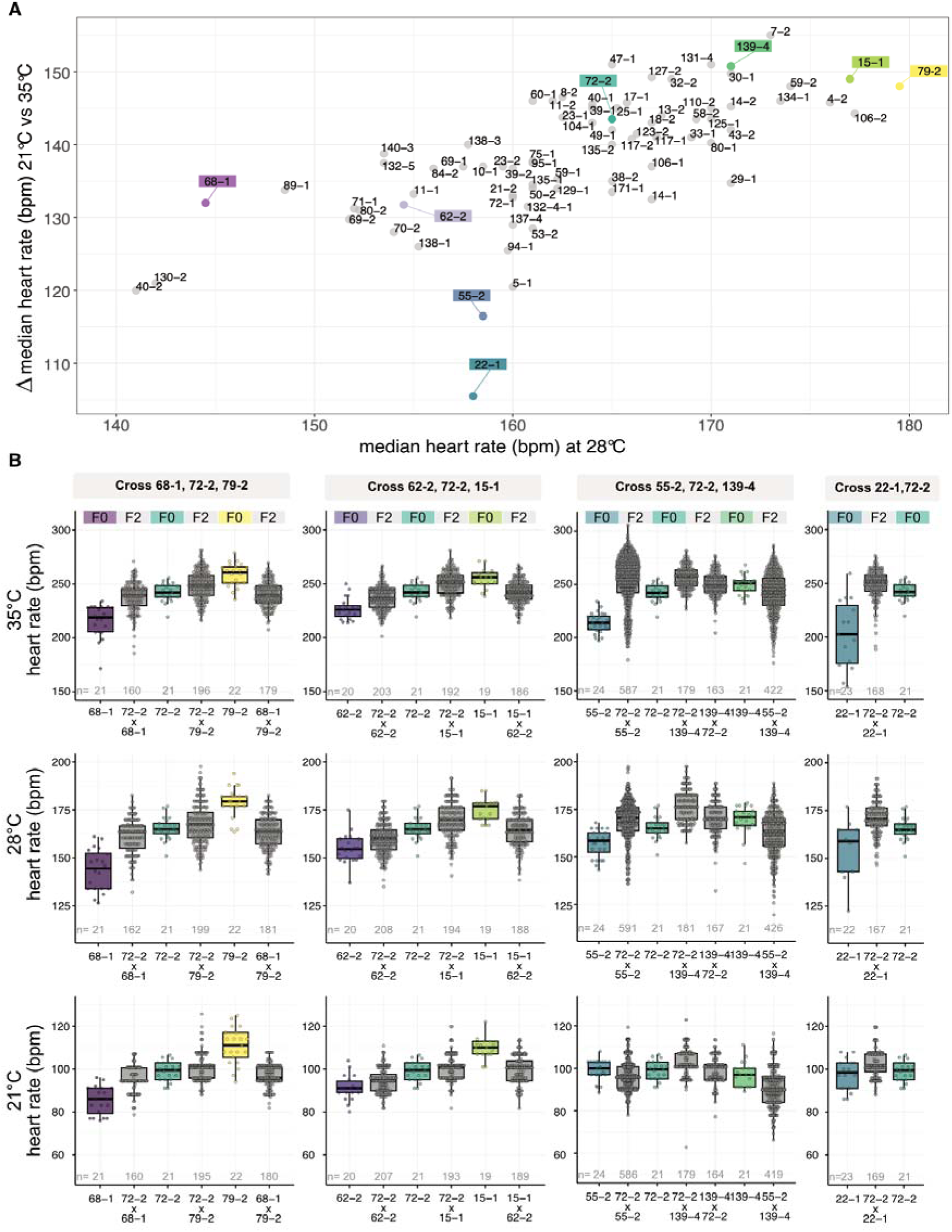
Strain-dependent heart rate change in the MIKK panel in response to temperature and subsequent heart rate segregation by 11 different crosses. **A:** Heart rate change from 21°C to 35°C and baseline heart rate at 28 °C varies across the MIKK panel strains, indicating different temperature-response characteristics. Strains selected for segregation analysis are highlighted. **B:** Heart rate of recombinant F2 embryos (grey) in comparison to the heart rate of the parental strains (F0) across the measured temperatures. Data is displayed as box plots (median +/- interquartile range between the 25th and 75th percentiles) and overlaid scatter plots of individual heart rate measurements (n) in beats per minute (bpm).

**Figure S2.**
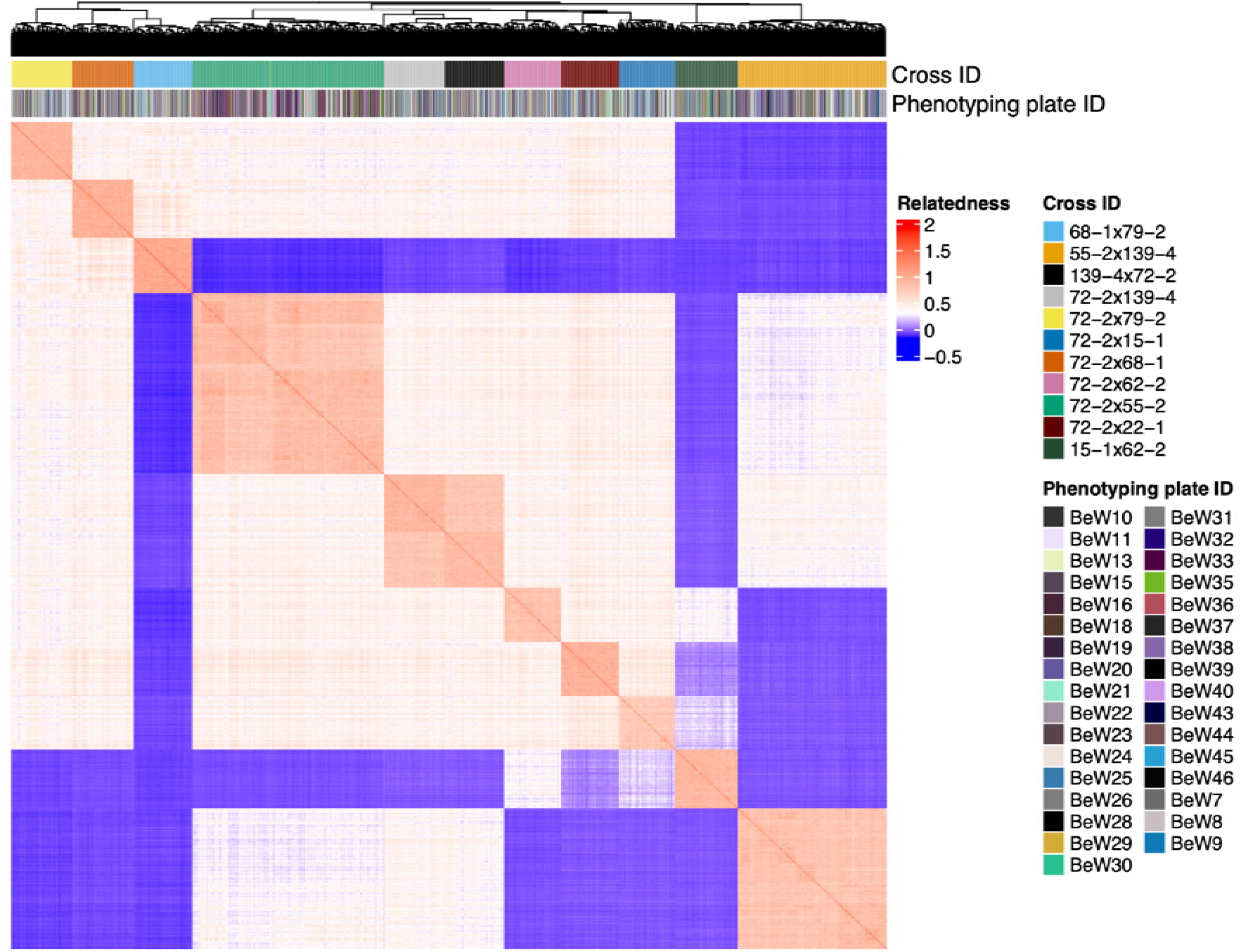
Genetic relatedness matrix of F2 samples. Heatmap of pairwise genetic relatedness among the samples of the multi-parental F2 population used in the GWAS analysis described in this work. Rows and columns are clustered by genetic relatedness, demonstrating spontaneous clustering of the samples belonging to the same F2 cross (“cross ID” annotation). The “Phenotyping plate ID” annotation demonstrates the randomisation achieved between phenotyping batches and population structure. Note the sub-clustering within the reciprocal cross (72-2 x 139-4 and 139-4 x 72-2), highlighting shared parental genetic contributions.

**Figure S3.**
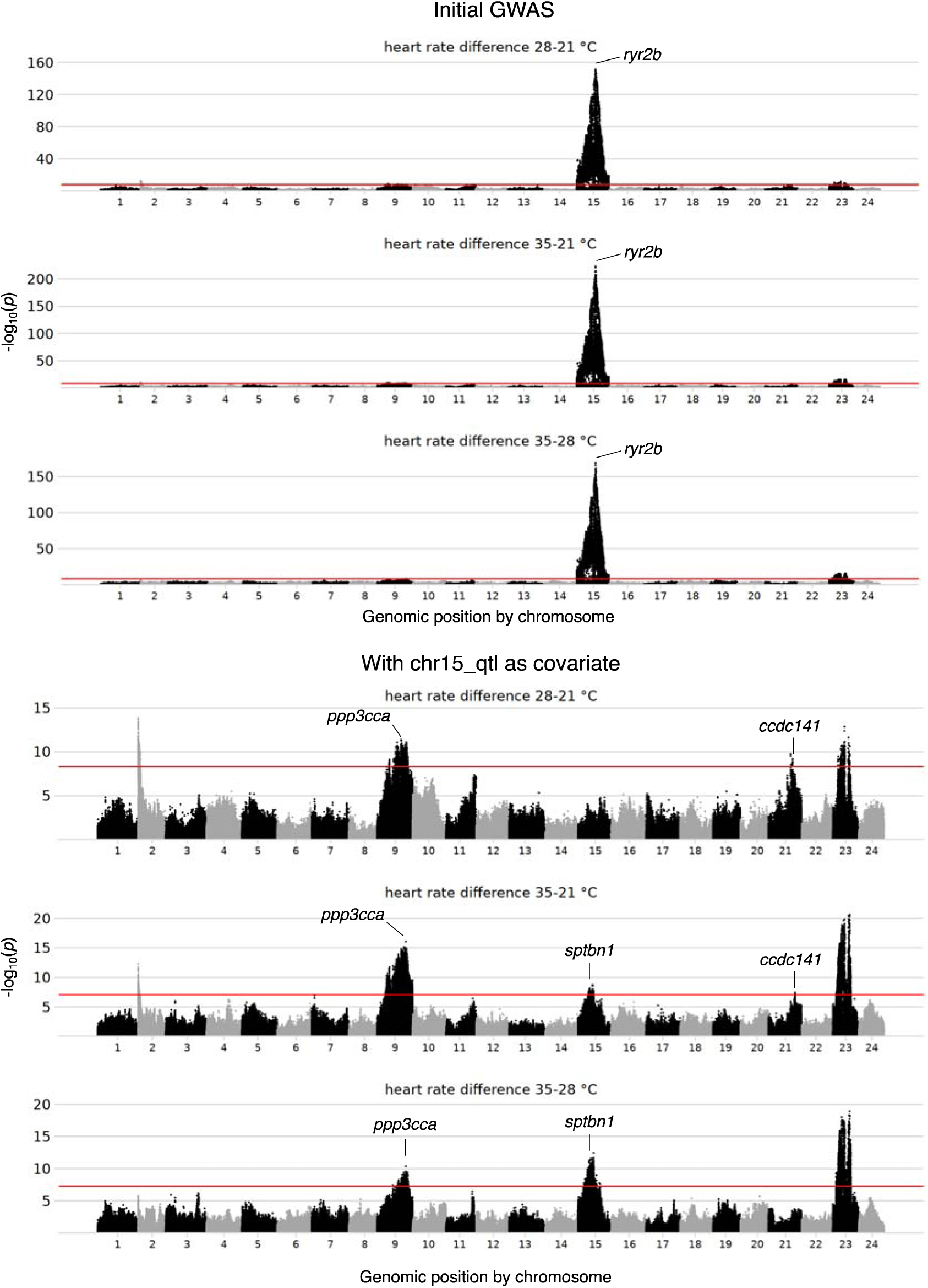
Manhattan plots of the heart rate temperature response phenotypes. Manhattan plots for the association of the difference in heart rate across temperature treatments with genetics. The significance threshold (red line) corresponds to the minimum *p*-value achieved over 100 permutations. The genes selected for experimental validation are indicated. In the bottom three panels, the strong locus at chromosome 15 was included as a covariate to unmask weaker effects.

**Figure S4.**
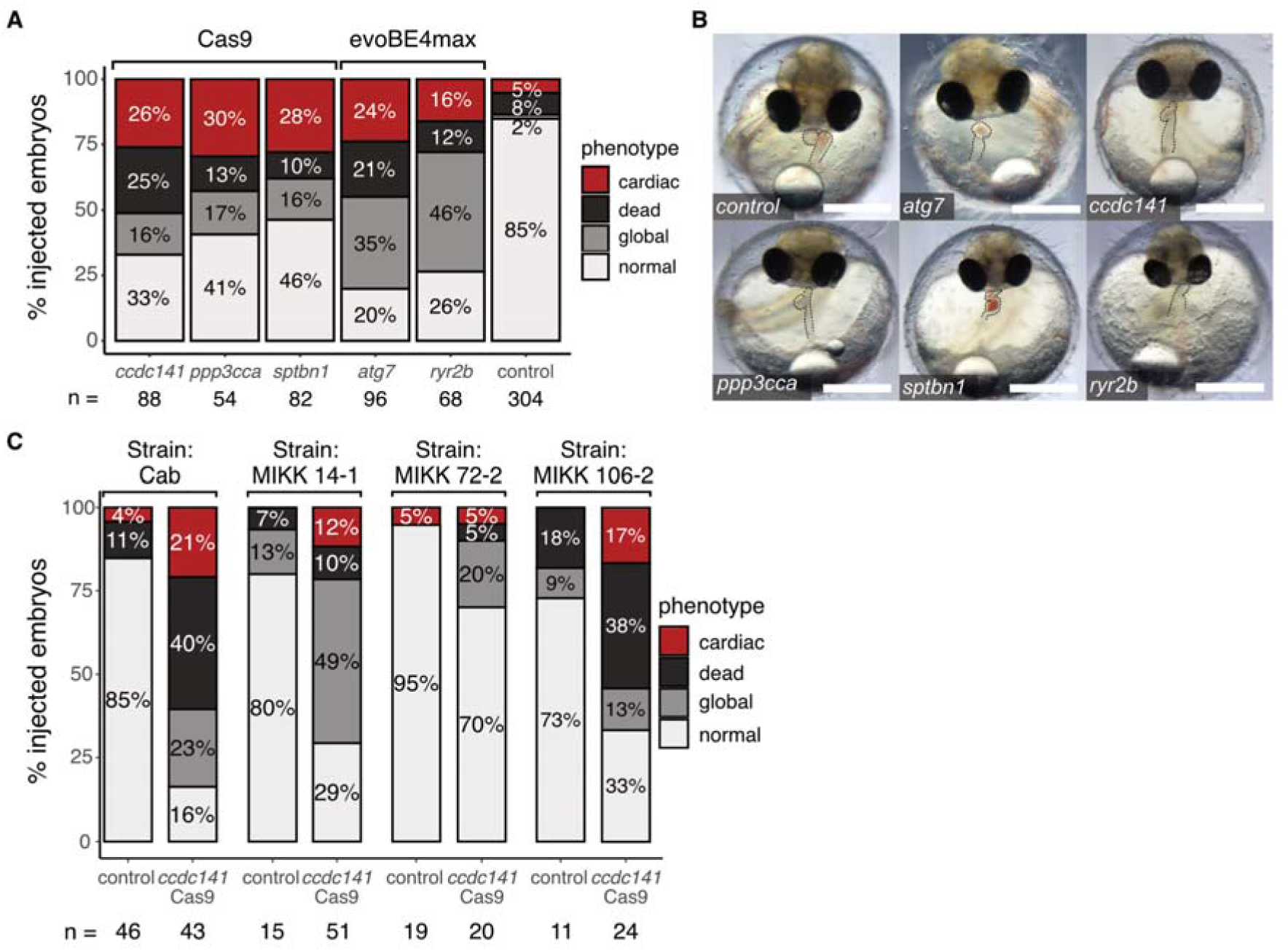
Phenotype proportions after gene editing of candidate genes. **A:** Phenotype proportions after CRISPR/Cas9- and base editor - mediated gene editing of five selected candidate genes reveals increased numbers of cardiac affected embryos 4 days post injection compared to the mock-injected control. **B:** Representative cardiac affected crispants and editants in comparison to normal developed control embryo (scale bar: 500µm). **C:** Phenotype proportions after CRISPR/Cas9-mediated genome targeting of *cccdc141* in four different medaka strains in comparison to mock-injected controls. n: number of injected embryos.

**Figure S5.**
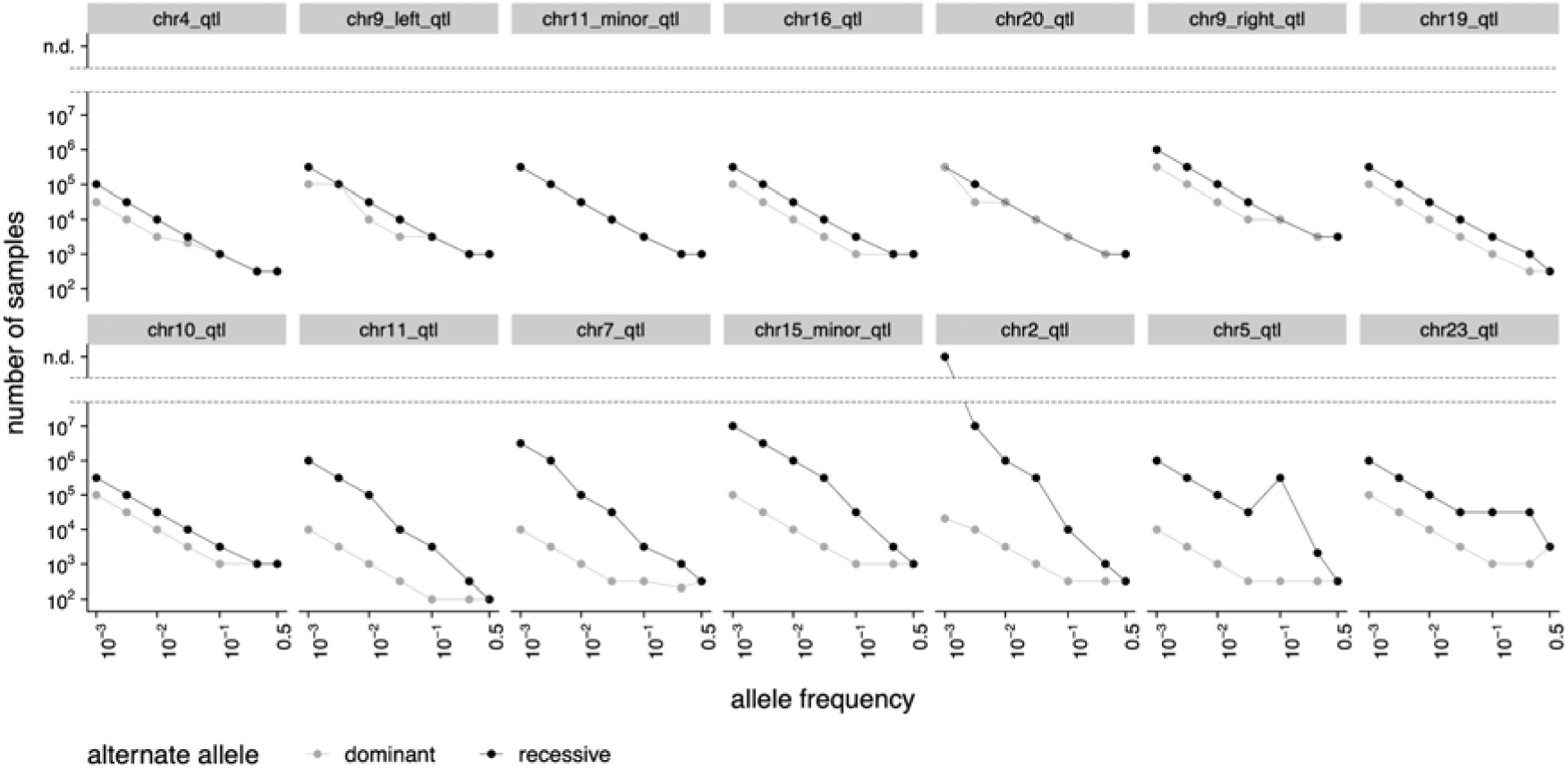
Simulation of the effect of dominance directionality. Simulation results analogous to the ones in **Figure 6**, but faceted by simulated QTL. These results use a G + E discovery model, with access to the causal variant, and are generated under a model that does not account for GxG effects. The QTLs are ordered from least to most dominant. In black the estimated sample size requirement for when the alternate allele at the locus is recessive, and in grey the estimated sample size requirement for when the alternate allele is dominant. Notice how the sample size required for discovery is systematically higher when the alternate allele is recessive (the most likely scenario in real populations where high effect recessive alleles are selected against).

**Figure S6.**
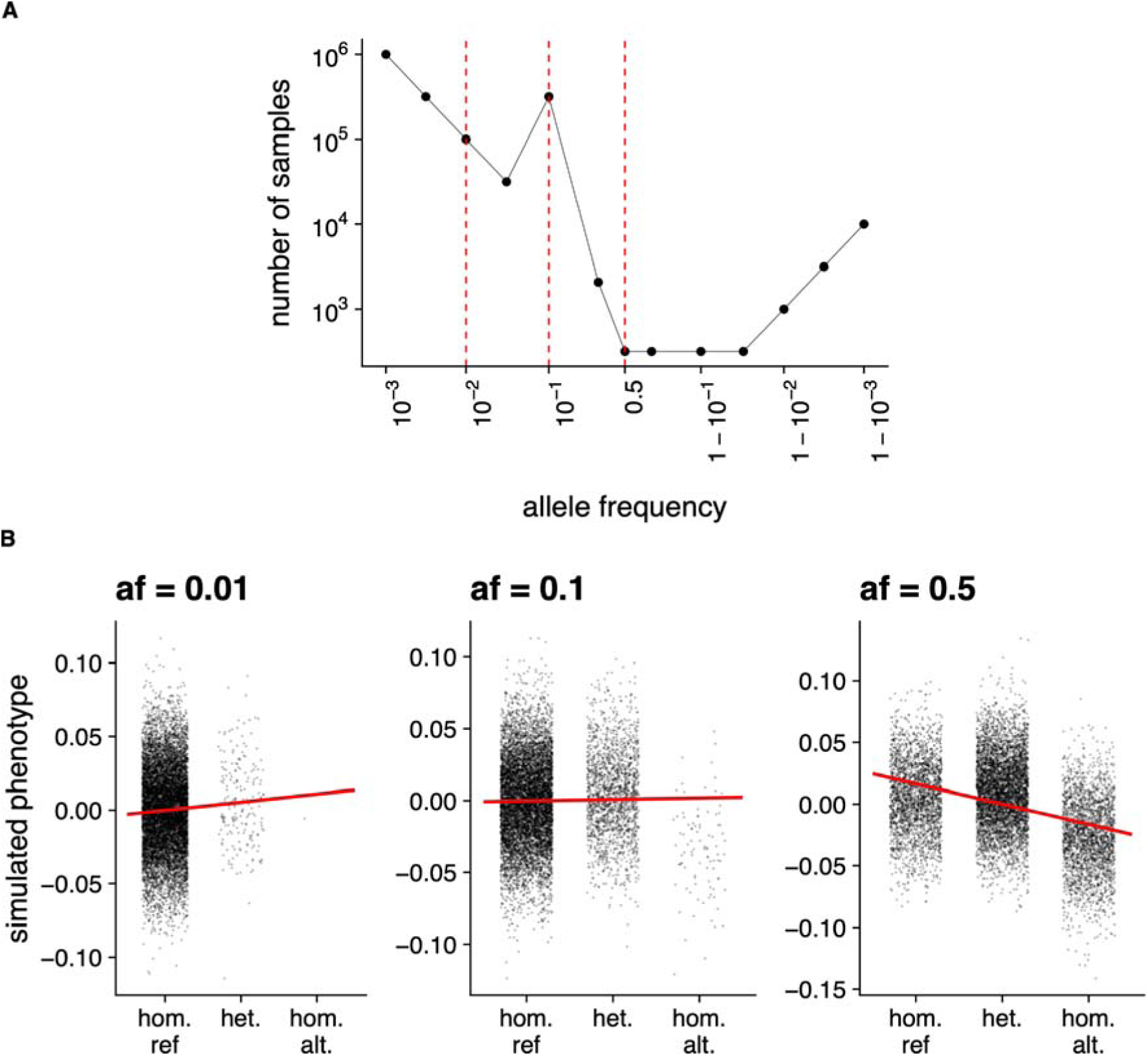
Consequences of overdominance. **A.** Extract from the simulation described in **Figure 6** showcasing an overdominant QTL (chr5_qtl). The results shown use a G + E discovery model, with access to the causal variant, and are generated under a model that does not account for GxG effects. Allele frequencies of 0.01, 0.1, and 0.5 are highlighted with red dashed vertical lines. Notice how at an allele frequency of 0.1 the required sample size increases sharply. **B.** Detailed plot of the sample-level simulation results for the allele frequencies (af) highlighted in panel A. Notice how the effect size estimate (red slope) depends on allele frequency. Going from an allele frequency of 0.01 to 0.5 the slope reverses direction. At an allele frequency of 0.1 this locus is not discoverable under a linear model because the effect of the heterozygous and homozygous samples cancels out almost exactly, leading to the flat slope in panel B and the spike in sample size in panel A.

**Figure S7.**
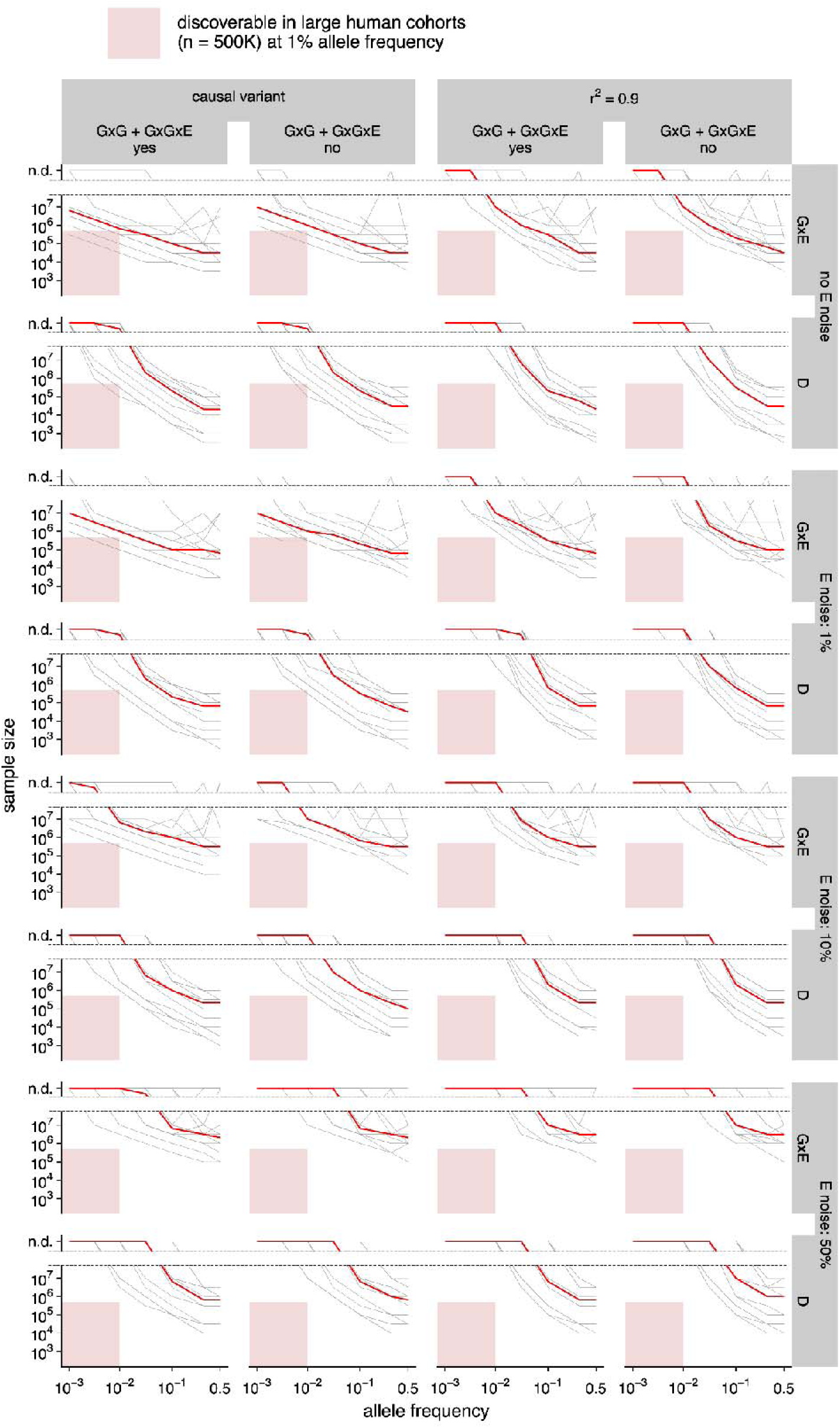
Simulation of the discoverability of non-additive effects. Simulation results analogous to the ones in **Figure 6**, but where the discoverability is evaluated against an additive G + E null model and not an E only model. Thus, the minimum sample size estimates reported here refer to what would be required to discover a significant non-additive effect (D: dominance, and GxE: gene-by-environment) when the additive effect is already accounted for.

**Figure S8.**
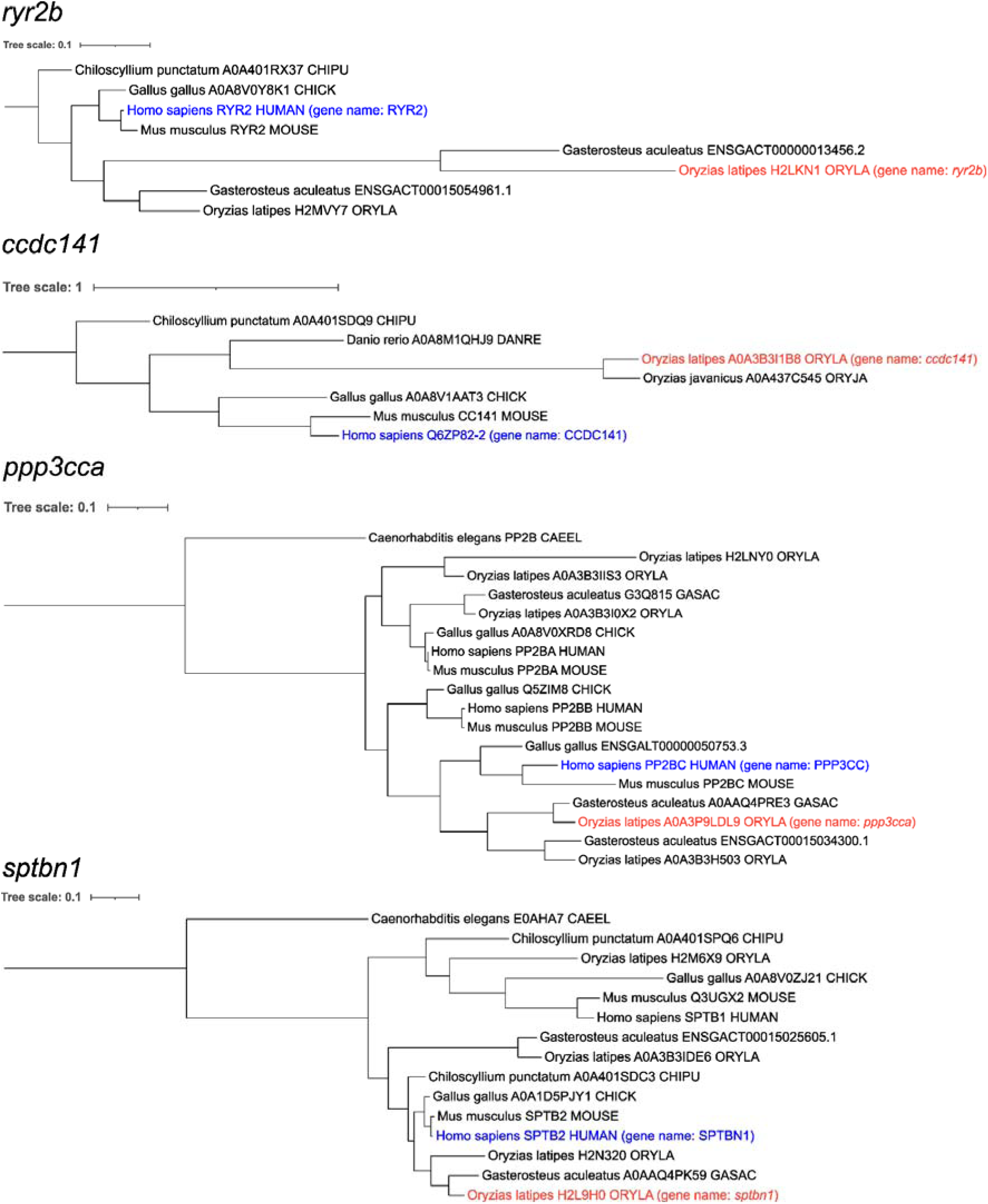
Gene trees for experimentally confirmed candidate genes. Gene trees for candidate medaka genes identified via QTL mapping and confirmed by gene editing (in red). Some of the confirmed candidate genes have a complex evolutionary history with multiple paralogues in different species, but in each case it was possible to identify a likely human orthologue or co-orthologue (in blue). Note that the tree identifiers refer to the Uniprot or ENSEMBL IDs used to obtain the protein sequence. The gene names corresponding to the confirmed medaka genes and the corresponding human orthologs or co-orthologs are written in parentheses.

## Supplementary Data

**Data S1. Putative causal variants.** List of putative causal variants detected for the genes *ryr2b*, *ccdc141*, and *atg7*.

**Data S2. Number of embryos from the MIKK panel inbred strains used for heart rate measurements.** Data underlying the plots in Figure 1A, 1D and S1.

**Data S3. Heart rate measurements of MIKK panel inbred strains.** Data underlying the plots in Figure 1A, 1D and S1.

**Data S4. Heart rate measurements of F2 embryos.** Data underlying the plots in Figure 1D and S1.

**Data S5. Numbers of cripsants, editants, and mock-injected controls used for heart rate measurements.** Data underlying the plots in Figure 4B. Only samples with a normal morphological phenotype (see Figure S4) are shown.

**Data S6. Heart rate measurements for the gene editing experiments.** Data underlying Figure 4B with heart rate measurements for control and gene-edited medaka embryos. Only samples with a normal morphological phenotype (see Figure S4) are shown.

**Data S7. Statistical test results and variance explained for GxE and dominance effects.** Data underlying the plots in Figure 3C, 3D, and 3E. Note that the GxE and DxE tests have 2 degrees of freedom because the temperature was treated as a categorical variable with 3 levels.

**Data S8. Statistical test results for GxG effects.** Data underlying the plot in Figure 5A. Note that the GxGxE test has 2 degrees of freedom because the temperature was treated as a categorical variable with 3 levels.

**Data S9. Statistical tests for reciprocal cross effect.** Test for the presence of an interaction effect between the reciprocal cross status (72-2x139-4 or 139-4x72-2) and the genotype at the QTLs determined to be polymorphic within the reciprocal cross (see methods for more details).

## References

Abdellaoui, A., Dolan, C. V., Verweij, K. J. H., & Nivard, M. G. (2022). Gene–environment correlations across geographic regions affect genome-wide association studies. Nature Genetics, 54(9), 1345–1354. 10.1038/s41588-022-01158-0

Abdellaoui, A., Yengo, L., Verweij, K. J. H., & Visscher, P. M. (2023). 15 years of GWAS discovery: Realizing the promise. The American Journal of Human Genetics, 110(2), 179–194. 10.1016/j.ajhg.2022.12.011

Aida, T. (1921). ON THE INHERITANCE OF COLOR IN A FRESH-WATER FISH, *APLOCHEILUS LATIPES* TEMMICK AND SCHLEGEL, WITH SPECIAL REFERENCE TO SEX-LINKED INHERITANCE. Genetics, 6(6), 554–573. 10.1093/genetics/6.6.554

Anshori, M. F., Musa, Y., Farid, M., Jayadi, M., Padjung, R., Kaimuddin, K., Huang, Y. C., Casimero, M., Bogayong, I., Suwarno, W. B., Sembiring, H., Purwoko, B. S., Nur, A., Wahyuni, W., Wasonga, D. O., & Seleiman, M. F. (2024). A comprehensive multivariate approach for GxE interaction analysis in early maturing rice varieties. Frontiers in Plant Science, 15. 10.3389/fpls.2024.1462981

Ashbrook, D. G., Arends, D., Prins, P., Mulligan, M. K., Roy, S., Williams, E. G., Lutz, C. M., Valenzuela, A., Bohl, C. J., Ingels, J. F., McCarty, M. S., Centeno, A. G., Hager, R., Auwerx, J., Lu, L., & Williams, R. W. (2021). A platform for experimental precision medicine: The extended BXD mouse family. Cell Systems, 12(3), 235–247.e9. 10.1016/j.cels.2020.12.002

Bakermans-Kranenburg, M. J., & IJzendoorn, M. H. van. (2015). The Hidden Efficacy of Interventions: Gene×Environment Experiments from a Differential Susceptibility Perspective. Annual Review of Psychology, 66(Volume 66, 2015), 381–409. 10.1146/annurev-psych-010814-015407

Baud, A., Hermsen, R., Guryev, V., Stridh, P., Graham, D., McBride, M. W., Foroud, T., Calderari, S., Diez, M., Ockinger, J., Beyeen, A. D., Gillett, A., Abdelmagid, N., Guerreiro-Cacais, A. O., Jagodic, M., Tuncel, J., Norin, U., Beattie, E., Huynh, N., … Rat Genome Sequencing and Mapping Consortium. (2013). Combined sequence-based and genetic mapping analysis of complex traits in outbred rats. Nature Genetics, 45(7), 767–775. 10.1038/ng.2644

Baud, A., Mulligan, M. K., Casale, F. P., Ingels, J. F., Bohl, C. J., Callebert, J., Launay, J.-M., Krohn, J., Legarra, A., Williams, R. W., & Stegle, O. (2017). Genetic Variation in the Social Environment Contributes to Health and Disease. PLOS Genetics, 13(1), e1006498. 10.1371/journal.pgen.1006498

Bert, B., Chmielewska, J., Bergmann, S., Busch, M., Driever, W., FingerOBaier, K., Hößler, J., Köhler, A., Leich, N., Misgeld, T., Nöldner, T., Reiher, A., Schartl, M., Seebach-Sproedt, A., Thumberger, T., Schönfelder, G., & Grune, B. (2016). Considerations for a European animal welfare standard to evaluate adverse phenotypes in teleost fish. The EMBO Journal, 35(11), 1151–1154. 10.15252/embj.201694448

Bick, A. G., Metcalf, G. A., Mayo, K. R., Lichtenstein, L., Rura, S., Carroll, R. J., Musick, A., Linder, J. E., Jordan, I. K., Nagar, S. D., Sharma, S., Meller, R., Basford, M., Boerwinkle, E., Cicek, M. S., Doheny, K. F., Eichler, E. E., Gabriel, S., Gibbs, R. A., … NIH All of Us Research Program Staff. (2024). Genomic data in the All of Us Research Program. Nature, 627(8003), 340–346. 10.1038/s41586-023-06957-x

Birker, K., Ge, S., Kirkland, N. J., Theis, J. L., Marchant, J., Fogarty, Z. C., Missinato, M. A., Kalvakuri, S., Grossfeld, P., Engler, A. J., Ocorr, K., Nelson, T. J., Colas, A. R., Olson, T. M., Vogler, G., & Bodmer, R. (2023). Mitochondrial MICOS complex genes, implicated in hypoplastic left heart syndrome, maintain cardiac contractility and actomyosin integrity. eLife, 12, e83385. 10.7554/eLife.83385

Bloom, J. S., Ehrenreich, I. M., Loo, W. T., Lite, T.-L. V., & Kruglyak, L. (2013). Finding the sources of missing heritability in a yeast cross. Nature, 494(7436), 234–237. 10.1038/nature11867

Bomba, L., Walter, K., & Soranzo, N. (2017). The impact of rare and low-frequency genetic variants in common disease. Genome Biology, 18(1), 77. 10.1186/s13059-017-1212-4

Bononi, A., Goto, K., Ak, G., Yoshikawa, Y., Emi, M., Pastorino, S., Carparelli, L., Ferro, A., Nasu, M., Kim, J.-H., Suarez, J. S., Xu, R., Tanji, M., Takinishi, Y., Minaai, M., Novelli, F., Pagano, I., Gaudino, G., Pass, H. I., … Carbone, M. (2020). Heterozygous germline BLM mutations increase susceptibility to asbestos and mesothelioma. Proceedings of the National Academy of Sciences, 117(52), 33466–33473. 10.1073/pnas.2019652117

Brem, R. B., Yvert, G., Clinton, R., & Kruglyak, L. (2002). Genetic Dissection of Transcriptional Regulation in Budding Yeast. Science, 296(5568), 752–755. 10.1126/science.1069516

Bycroft, C., Freeman, C., Petkova, D., Band, G., Elliott, L. T., Sharp, K., Motyer, A., Vukcevic, D., Delaneau, O., O’Connell, J., Cortes, A., Welsh, S., Young, A., Effingham, M., McVean, G., Leslie, S., Allen, N., Donnelly, P., & Marchini, J. (2018). The UK Biobank resource with deep phenotyping and genomic data. Nature, 562(7726), 203–209. 10.1038/s41586-018-0579-z

Campbell, M. J., Czosek, R. J., Hinton, R. B., & Miller, E. M. (2015). Exon 3 deletion of ryanodine receptor causes left ventricular noncompaction, worsening catecholaminergic polymorphic ventricular tachycardia, and sudden cardiac arrest. American Journal of Medical Genetics Part A, 167(9), 2197–2200. 10.1002/ajmg.a.37140

Caspi, A., Sugden, K., Moffitt, T. E., Taylor, A., Craig, I. W., Harrington, H., McClay, J., Mill, J., Martin, J., Braithwaite, A., & Poulton, R. (2003). Influence of Life Stress on Depression: Moderation by a Polymorphism in the 5-HTT Gene. Science, 301(5631), 386–389. 10.1126/science.1083968

Chen, S.-A. A., Kern, A. F., Ang, R. M. L., Xie, Y., & Fraser, H. B. (2023). Gene-by-environment interactions are pervasive among natural genetic variants. Cell Genomics, 3(4), 100273. 10.1016/j.xgen.2023.100273

Churchill, G. A., Airey, D. C., Allayee, H., Angel, J. M., Attie, A. D., Beatty, J., Beavis, W. D., Belknap, J. K., Bennett, B., Berrettini, W., Bleich, A., Bogue, M., Broman, K. W., Buck, K. J., Buckler, E., Burmeister, M., Chesler, E. J., Cheverud, J. M., Clapcote, S., … The Complex Trait Consortium. (2004). The Collaborative Cross, a community resource for the genetic analysis of complex traits. Nature Genetics, 36(11), 1133–1137. 10.1038/ng1104-1133

Claire, D.-R., & Hervé, P. (2018). 46th european mathematical genetics meeting (EMGM) 2018, cagliari, italy, april 18-20, 2018: Abstracts. Human Heredity, 83(1), 1–29. 10.1159/000488519

Collaborative Cross Consortium. (2012). The Genome Architecture of the Collaborative Cross Mouse Genetic Reference Population. Genetics, 190(2), 389–401. 10.1534/genetics.111.132639

Cooper, M., & Messina, C. D. (2021). Can We Harness “Enviromics” to Accelerate Crop Improvement by Integrating Breeding and Agronomy? Frontiers in Plant Science, 12, 735143. 10.3389/fpls.2021.735143

Copley, J. P., Hayes, B. J., Ross, E. M., Speight, S., Fordyce, G., Wood, B. J., & Engle, B. N. (2024). Investigating genotype by environment interaction for beef cattle fertility traits in commercial herds in northern Australia with multi-trait analysis. Genetics Selection Evolution, 56(1), 70. 10.1186/s12711-024-00936-0

Cornean, A., Gierten, J., Welz, B., Mateo, J. L., Thumberger, T., & Wittbrodt, J. (2022). Precise in vivo functional analysis of DNA variants with base editing using ACEofBASEs target prediction. eLife, 11, e72124. 10.7554/eLife.72124

Costanzo, M., Hou, J., Messier, V., Nelson, J., Rahman, M., VanderSluis, B., Wang, W., Pons, C., Ross, C., Ušaj, M., San Luis, B.-J., Shuteriqi, E., Koch, E. N., Aloy, P., Myers, C. L., Boone, C., & Andrews, B. (2021). Environmental robustness of the global yeast genetic interaction network. Science, 372(6542), eabf8424. 10.1126/science.abf8424

Cunningham, F., Allen, J. E., Allen, J., Alvarez-Jarreta, J., Amode, M. R., Armean, I. M., Austine-Orimoloye, O., Azov, A. G., Barnes, I., Bennett, R., Berry, A., Bhai, J., Bignell, A., Billis, K., Boddu, S., Brooks, L., Charkhchi, M., Cummins, C., Fioretto, L. D. R., … Flicek, P. (2021). Ensembl 2022. Nucleic Acids Research, 50(D1), D988–D995. 10.1093/nar/gkab1049

Currant, H., Fitzgerald, T. W., Patel, P. J., Khawaja, A. P., Consortium, U. B. E. and V., Webster, A. R., Mahroo, O. A., & Birney, E. (2023). Sub-cellular level resolution of common genetic variation in the photoreceptor layer identifies continuum between rare disease and common variation. PLOS Genetics, 19(2), e1010587. 10.1371/journal.pgen.1010587

Danecek, P., Bonfield, J. K., Liddle, J., Marshall, J., Ohan, V., Pollard, M. O., Whitwham, A., Keane, T., McCarthy, S. A., Davies, R. M., & Li, H. (2021). Twelve years of SAMtools and BCFtools. GigaScience, 10(2). 10.1093/gigascience/giab008

den Hoed, M., Eijgelsheim, M., Esko, T., Brundel, B. J. J. M., Peal, D. S., Evans, D. M., Nolte, I. M., Segrè, A. V., Holm, H., Handsaker, R. E., Westra, H.-J., Johnson, T., Isaacs, A., Yang, J., Lundby, A., Zhao, J. H., Kim, Y. J., Go, M. J., Almgren, P., … Loos, R. J. F. (2013). Identification of heart rate–associated loci and their effects on cardiac conduction and rhythm disorders. Nature Genetics, 45(6), 621–631. 10.1038/ng.2610

Di Tommaso, P., Chatzou, M., Floden, E. W., Barja, P. P., Palumbo, E., & Notredame, C. (2017). Nextflow enables reproducible computational workflows. Nature Biotechnology, 35(4), 316–319. 10.1038/nbt.3820

Dumont, B. L., Gatti, D. M., Ballinger, M. A., Lin, D., Phifer-Rixey, M., Sheehan, M. J., Suzuki, T. A., Wooldridge, L. K., Frempong, H. O., Lawal, R. A., Churchill, G. A., Lutz, C., Rosenthal, N., White, J. K., & Nachman, M. W. (2024). Into the Wild: A novel wild-derived inbred strain resource expands the genomic and phenotypic diversity of laboratory mouse models. PLoS Genetics, 20(4), e1011228. 10.1371/journal.pgen.1011228

Evans, K. S., van Wijk, M. H., McGrath, P. T., Andersen, E. C., & Sterken, M. G. (2021). From QTL to gene: *C. elegans* facilitates discoveries of the genetic mechanisms underlying natural variation. Trends in Genetics, 37(10), 933–947. 10.1016/j.tig.2021.06.005

Fennewald, D. J., Weaber, R. L., & Lamberson, W. R. (2018). Genotype by environment interaction for stayability of Red Angus in the United States. Journal of Animal Science, 96(2), 422–429. 10.1093/jas/skx080

Ferreira, M. S., Stricker, S., Fitzgerald, T., Monahan, J., Defranoux, F., Watson, P., Welz, B., Hammouda, O., Wittbrodt, J., & Birney, E. (2024). FEHAT: Efficient, large scale and automated heartbeat detection in Medaka fish embryos. Bioinformatics, 40(12), btae664. 10.1093/bioinformatics/btae664

Fisher, R. A. (1919). XV.—The Correlation between Relatives on the Supposition of Mendelian Inheritance. Earth and Environmental Science Transactions of The Royal Society of Edinburgh, 52(2), 399–433. 10.1017/S0080456800012163

Fitzgerald, T., Brettell, I., Leger, A., Wolf, N., Kusminski, N., Monahan, J., Barton, C., Herder, C., Aadepu, N., Gierten, J., Becker, C., Hammouda, O. T., Hasel, E., Lischik, C., Lust, K., Sokolova, N., Suzuki, R., Tsingos, E., Tavhelidse, T., … Loosli, F. (2022). The Medaka Inbred Kiyosu-Karlsruhe (MIKK) panel. Genome Biology, 23(1), 59. 10.1186/s13059-022-02623-z

Fukuda, T., Sugita, S., Inatome, R., & Yanagi, S. (2010). CAMDI, a Novel Disrupted in Schizophrenia 1 (DISC1)-binding Protein, Is Required for Radial Migration*. Journal of Biological Chemistry, 285(52), 40554–40561. 10.1074/jbc.M110.179481

Furutani-Seiki, M., & Wittbrodt, J. (2004). Medaka and zebrafish, an evolutionary twin study. Mechanisms of Development, 121(7–8), 629–637. 10.1016/j.mod.2004.05.010

Gaziano, J. M., Concato, J., Brophy, M., Fiore, L., Pyarajan, S., Breeling, J., Whitbourne, S., Deen, J., Shannon, C., Humphries, D., Guarino, P., Aslan, M., Anderson, D., LaFleur, R., Hammond, T., Schaa, K., Moser, J., Huang, G., Muralidhar, S., … O’Leary, T. J. (2016). Million Veteran Program: A mega-biobank to study genetic influences on health and disease. Journal of Clinical Epidemiology, 70, 214–223. 10.1016/j.jclinepi.2015.09.016

Gierten, J., Pylatiuk, C., Hammouda, O. T., Schock, C., Stegmaier, J., Wittbrodt, J., Gehrig, J., & Loosli, F. (2020). Automated high-throughput heartbeat quantification in medaka and zebrafish embryos under physiological conditions. Scientific Reports, 10(1), 2046. 10.1038/s41598-020-58563-w

Gierten, J., Welz, B., Fitzgerald, T., Thumberger, T., Agarwal, R., Hummel, O., Leger, A., Weber, P., Naruse, K., Hassel, D., Hübner, N., Birney, E., & Wittbrodt, J. (2025). Natural genetic variation quantitatively regulates heart rate and dimension. Nature Communications, 16(1), 4062. 10.1038/s41467-025-59425-7

Hammond, J. (1947). Animal Breeding in Relation to Nutrition and Environmental Conditions. Biological Reviews, 22(3), 195–213. 10.1111/j.1469-185X.1947.tb00330.x

Hammouda, O. T., Wu, M. Y., Kaul, V., Gierten, J., Thumberger, T., & Wittbrodt, J. (2021). In vivo identification and validation of novel potential predictors for human cardiovascular diseases. PLOS ONE, 16(12), e0261572. 10.1371/journal.pone.0261572

Hanssen, F., Garcia, M. U., Folkersen, L., Pedersen, A. S., Lescai, F., Jodoin, S., Miller, E., Wacker, O., Smith, N., Community, N.-C., Gabernet, G., & Nahnsen, S. (2023). Scalable and efficient DNA sequencing analysis on different compute infrastructures aiding variant discovery (p. 2023.07.19.549462). bioRxiv. 10.1101/2023.07.19.549462

Hasan, M. M., Thomson, P. C., Raadsma, H. W., & Khatkar, M. S. (2024). A Review and Meta-Analysis of Genotype by Environment Interaction in Commercial Shrimp Breeding. Genes, 15(9), Article 9. 10.3390/genes15091222

Hawn, S. E., Sheerin, C. M., Webb, B. T., Peterson, R. E., Do, E. K., Dick, D., Kendler, K. S., Bacanu, S.-A., & Amstadter, A. B. (2018). Replication of the Interaction of PRKG1 and Trauma Exposure on Alcohol Misuse in an Independent African American Sample. Journal of Traumatic Stress, 31(6), 927–932. 10.1002/jts.22339

Hayes, B. J., Daetwyler, H. D., & Goddard, M. E. (2016). Models for Genome × Environment Interaction: Examples in Livestock. Crop Science, 56(5), 2251–2259. 10.2135/cropsci2015.07.0451

Herrera-Luis, E., Benke, K., Volk, H., Ladd-Acosta, C., & Wojcik, G. L. (2024). Gene–environment interactions in human health. Nature Reviews Genetics, 25(11), 768–784. 10.1038/s41576-024-00731-z

Hill, W. G., Goddard, M. E., & Visscher, P. M. (2008). Data and Theory Point to Mainly Additive Genetic Variance for Complex Traits. PLOS Genetics, 4(2), e1000008. 10.1371/journal.pgen.1000008

Hivert, V., Sidorenko, J., Rohart, F., Goddard, M. E., Yang, J., Wray, N. R., Yengo, L., & Visscher, P. M. (2021). Estimation of non-additive genetic variance in human complex traits from a large sample of unrelated individuals. The American Journal of Human Genetics, 108(5), 786–798. 10.1016/j.ajhg.2021.02.014

Ho, W.-C., & Zhang, J. (2014). The Genotype–Phenotype Map of Yeast Complex Traits: Basic Parameters and the Role of Natural Selection. Molecular Biology and Evolution, 31(6), 1568–1580. 10.1093/molbev/msu131

Hu, S., Ferreira, L. A. F., Shi, S., Hellenthal, G., Marchini, J., Lawson, D. J., & Myers, S. R. (2025). Fine-scale population structure and widespread conservation of genetic effect sizes between human groups across traits. Nature Genetics, 57(2), 379–389. 10.1038/s41588-024-02035-8

Huang, W., Carbone, M. A., Lyman, R. F., Anholt, R. R. H., & Mackay, T. F. C. (2020a). Genotype by environment interaction for gene expression in Drosophila melanogaster. Nature Communications, 11(1), 5451. 10.1038/s41467-020-19131-y

Huang, W., Carbone, M. A., Lyman, R. F., Anholt, R. R. H., & Mackay, T. F. C. (2020b). Genotype by environment interaction for gene expression in Drosophila melanogaster. Nature Communications, 11(1), 5451. 10.1038/s41467-020-19131-y

Iwamatsu, T. (2004). Stages of normal development in the medaka Oryzias latipes. Mechanisms of Development, 121(7–8), 605–618. 10.1016/j.mod.2004.03.012

Jelier, R., Semple, J. I., Garcia-Verdugo, R., & Lehner, B. (2011). Predicting phenotypic variation in yeast from individual genome sequences. Nature Genetics, 43(12), 1270–1274. 10.1038/ng.1007

Jensen, B., Wang, T., Christoffels, V. M., & Moorman, A. F. M. (2013). Evolution and development of the building plan of the vertebrate heart. Biochimica et Biophysica Acta (BBA) - Molecular Cell Research, 1833(4), 783–794. 10.1016/j.bbamcr.2012.10.004

Johnson, E. O., Chen, L.-S., Breslau, N., Hatsukami, D., Robbins, T., Saccone, N. L., Grucza, R. A., & Bierut, L. J. (2010). Peer smoking and the nicotinic receptor genes: An examination of genetic and environmental risks for nicotine dependence. Addiction, 105(11), 2014–2022. 10.1111/j.1360-0443.2010.03074.x

Karlsson, M., Zhang, C., Méar, L., Zhong, W., Digre, A., Katona, B., Sjöstedt, E., Butler, L., Odeberg, J., Dusart, P., Edfors, F., Oksvold, P., Von Feilitzen, K., Zwahlen, M., Arif, M., Altay, O., Li, X., Ozcan, M., Mardinoglu, A., … Lindskog, C. (2021). A single–cell type transcriptomics map of human tissues. Science Advances, 7(31), eabh2169. 10.1126/sciadv.abh2169

Kasahara, M., Naruse, K., Sasaki, S., Nakatani, Y., Qu, W., Ahsan, B., Yamada, T., Nagayasu, Y., Doi, K., Kasai, Y., Jindo, T., Kobayashi, D., Shimada, A., Toyoda, A., Kuroki, Y., Fujiyama, A., Sasaki, T., Shimizu, A., Asakawa, S., … Kohara, Y. (2007). The medaka draft genome and insights into vertebrate genome evolution. Nature, 447(7145), 714–719. 10.1038/nature05846

Keaton, J. M., Kamali, Z., Xie, T., Vaez, A., Williams, A., Goleva, S. B., Ani, A., Evangelou, E., Hellwege, J. N., Yengo, L., Young, W. J., Traylor, M., Giri, A., Zheng, Z., Zeng, J., Chasman, D. I., Morris, A. P., Caulfield, M. J., Hwang, S.-J., … Warren, H. R. (2024). Genome-wide analysis in over 1 million individuals of European ancestry yields improved polygenic risk scores for blood pressure traits. Nature Genetics, 56(5), 778–791. 10.1038/s41588-024-01714-w

Kebede, F. G., Komen, H., Dessie, T., Hanotte, O., Kemp, S., Alemu, S. W., & Bastiaansen, J. W. M. (2024). Multi-environment performance analysis identifies more productive and widely adapted chicken breeds for smallholder farmers. Frontiers in Sustainable Food Systems, 8. 10.3389/fsufs.2024.1441295

Kichaev, G., Bhatia, G., Loh, P.-R., Gazal, S., Burch, K., Freund, M. K., Schoech, A., Pasaniuc, B., & Price, A. L. (2019). Leveraging Polygenic Functional Enrichment to Improve GWAS Power. The American Journal of Human Genetics, 104(1), 65–75. 10.1016/j.ajhg.2018.11.008

Kirchmaier, S., Naruse, K., Wittbrodt, J., & Loosli, F. (2015). The Genomic and Genetic Toolbox of the Teleost Medaka (*Oryzias latipes*). Genetics, 199(4), 905–918. 10.1534/genetics.114.173849

Kover, P. X., Valdar, W., Trakalo, J., Scarcelli, N., Ehrenreich, I. M., Purugganan, M. D., Durrant, C., & Mott, R. (2009). A Multiparent Advanced Generation Inter-Cross to Fine-Map Quantitative Traits in Arabidopsis thaliana. PLOS Genetics, 5(7), e1000551. 10.1371/journal.pgen.1000551

Kurki, M. I., Karjalainen, J., Palta, P., Sipilä, T. P., Kristiansson, K., Donner, K. M., Reeve, M. P., Laivuori, H., Aavikko, M., Kaunisto, M. A., Loukola, A., Lahtela, E., Mattsson, H., Laiho, P., Della Briotta Parolo, P., Lehisto, A. A., Kanai, M., Mars, N., Rämö, J., … Palotie, A. (2023). FinnGen provides genetic insights from a well-phenotyped isolated population. Nature, 613(7944), 508–518. 10.1038/s41586-022-05473-8

Kuznetsova, A., Brockhoff, P. B., & Christensen, R. H. B. (2017). lmerTest Package: Tests in Linear Mixed Effects Models. Journal of Statistical Software, 82, 1–26. 10.18637/jss.v082.i13

Lachance, J., Jung, L., & True, J. R. (2013). Genetic Background and GxE Interactions Modulate the Penetrance of a Naturally Occurring Wing Mutation in Drosophila melanogaster. G3 Genes|Genomes|Genetics, 3(11), 1893–1901. 10.1534/g3.113.007831

Leitsalu, L., & Metspalu, A. (2017). Chapter 8 - From Biobanking to Precision Medicine: The Estonian Experience. In G. S. Ginsburg & H. F. Willard (Eds.), Genomic and Precision Medicine (Third Edition) (pp. 119–129). Academic Press. 10.1016/B978-0-12-800681-8.00008-6

Letunic, I., & Bork, P. (2024). Interactive Tree of Life (iTOL) v6: Recent updates to the phylogenetic tree display and annotation tool. Nucleic Acids Research, 52(W1), W78–W82. 10.1093/nar/gkae268

Loosli, F., Köster, R. W., Carl, M., Krone, A., & Wittbrodt, J. (1998). Six3, a medaka homologue of the Drosophila homeobox gene sine oculis is expressed in the anterior embryonic shield and the developing eye. Mechanisms of Development, 74(1–2), 159–164. 10.1016/S0925-4773(98)00055-0

Loosli, F., Köster, R. W., Carl, M., Kühnlein, R., Henrich, T., Mücke, M., Krone, A., & Wittbrodt, J. (2000). A genetic screen for mutations affecting embryonic development in medaka fish (Oryzias latipes). Mechanisms of Development, 97(1–2), 133–139. 10.1016/S0925-4773(00)00406-8

Maazi, H., Hartiala, J. A., Suzuki, Y., Crow, A. L., Shafiei Jahani, P., Lam, J., Patel, N., Rigas, D., Han, Y., Huang, P., Eskin, E., Lusis, Aldons. J., Gilliland, F. D., Akbari, O., & Allayee, H. (2019). A GWAS approach identifies Dapp1 as a determinant of air pollution-induced airway hyperreactivity. PLOS Genetics, 15(12), e1008528. 10.1371/journal.pgen.1008528

Mackay, I. J., Bansept-Basler, P., Barber, T., Bentley, A. R., Cockram, J., Gosman, N., Greenland, A. J., Horsnell, R., Howells, R., O’Sullivan, D. M., Rose, G. A., & Howell, P. J. (2014). An Eight-Parent Multiparent Advanced Generation Inter-Cross Population for Winter-Sown Wheat: Creation, Properties, and Validation. G3 Genes|Genomes|Genetics, 4(9), 1603–1610. 10.1534/g3.114.012963

Mackay, T. F. C. (2014). Epistasis and quantitative traits: Using model organisms to study gene–gene interactions. Nature Reviews Genetics, 15(1), 22–33. 10.1038/nrg3627

Mackay, T. F. C., & Anholt, R. R. H. (2024). Pleiotropy, epistasis and the genetic architecture of quantitative traits. Nature Reviews Genetics, 25(9), 639–657. 10.1038/s41576-024-00711-3

Mackay, T. F. C., & Huang, W. (2018). Charting the genotype–phenotype map: Lessons from the Drosophila melanogaster Genetic Reference Panel. WIREs Developmental Biology, 7(1), e289. 10.1002/wdev.289

Mackay, T. F. C., Richards, S., Stone, E. A., Barbadilla, A., Ayroles, J. F., Zhu, D., Casillas, S., Han, Y., Magwire, M. M., Cridland, J. M., Richardson, M. F., Anholt, R. R. H., Barrón, M., Bess, C., Blankenburg, K. P., Carbone, M. A., Castellano, D., Chaboub, L., Duncan, L., … Gibbs, R. A. (2012). The Drosophila melanogaster Genetic Reference Panel. Nature, 482(7384), 173–178. 10.1038/nature10811

McLaren, W., Gil, L., Hunt, S. E., Riat, H. S., Ritchie, G. R. S., Thormann, A., Flicek, P., & Cunningham, F. (2016). The ensembl variant effect predictor. Genome Biology, 17(1). 10.1186/s13059-016-0974-4

Minh, B. Q., Schmidt, H. A., Chernomor, O., Schrempf, D., Woodhams, M. D., von Haeseler, A., & Lanfear, R. (2020). IQ-TREE 2: New Models and Efficient Methods for Phylogenetic Inference in the Genomic Era. Molecular Biology and Evolution, 37(5), 1530–1534. 10.1093/molbev/msaa015

Modafferi, S., Zhong, X., Kleensang, A., Murata, Y., Fagiani, F., Pamies, D., Hogberg, H. T., Calabrese, V., Lachman, H., Hartung, T., & Smirnova, L. (2021). Gene–Environment Interactions in Developmental Neurotoxicity: A Case Study of Synergy between Chlorpyrifos and CHD8 Knockout in Human BrainSpheres. Environmental Health Perspectives, 129(7), 077001. 10.1289/EHP8580

Morozova, T. V., Huang, W., Pray, V. A., Whitham, T., Anholt, R. R. H., & Mackay, T. F. C. (2015). Polymorphisms in early neurodevelopmental genes affect natural variation in alcohol sensitivity in adult drosophila. BMC Genomics, 16(1), 865. 10.1186/s12864-015-2064-5

Morse, H. C. I. (2012). Origins of Inbred Mice. https://shop.elsevier.com/books/origins-of-inbred-mice/morse/978-0-12-507850-4

Motsinger-Reif, A. A., Reif, D. M., Akhtari, F. S., House, J. S., Campbell, C. R., Messier, K. P., Fargo, D. C., Bowen, T. A., Nadadur, S. S., Schmitt, C. P., Pettibone, K. G., Balshaw, D. M., Lawler, C. P., Newton, S. A., Collman, G. W., Miller, A. K., Merrick, B. A., Cui, Y., Anchang, B., … Woychik, R. (2024). Gene-environment interactions within a precision environmental health framework. Cell Genomics, 4(7), 100591. 10.1016/j.xgen.2024.100591

Mott, R., Talbot, C. J., Turri, M. G., Collins, A. C., & Flint, J. (2000). A method for fine mapping quantitative trait loci in outbred animal stocks. Proceedings of the National Academy of Sciences, 97(23), 12649–12654. 10.1073/pnas.230304397

Mott, R., Yuan, W., Kaisaki, P., Gan, X., Cleak, J., Edwards, A., Baud, A., & Flint, J. (2014). The Architecture of Parent-of-Origin Effects in Mice. Cell, 156(1), 332–342. 10.1016/j.cell.2013.11.043

Nagai, A., Hirata, M., Kamatani, Y., Muto, K., Matsuda, K., Kiyohara, Y., Ninomiya, T., Tamakoshi, A., Yamagata, Z., Mushiroda, T., Murakami, Y., Yuji, K., Furukawa, Y., Zembutsu, H., Tanaka, T., Ohnishi, Y., Nakamura, Y., Shiono, M., Misumi, K., … Kubo, M. (2017). Overview of the BioBank Japan Project: Study design and profile. Journal of Epidemiology, 27(3, Supplement), S2–S8. 10.1016/j.je.2016.12.005

Nakayama, T., Tanikawa, M., Okushi, Y., Itoh, T., Shimmura, T., Maruyama, M., Yamaguchi, T., Matsumiya, A., Shinomiya, A., Guh, Y.-J., Chen, J., Naruse, K., Kudoh, H., Kondo, Y., Naoki, H., Aoki, K., Nagano, A. J., & Yoshimura, T. (2023). A transcriptional program underlying the circannual rhythms of gonadal development in medaka. Proceedings of the National Academy of Sciences, 120(52), e2313514120. 10.1073/pnas.2313514120

Napier, J. D., Heckman, R. W., & Juenger, T. E. (2023). Gene-by-environment interactions in plants: Molecular mechanisms, environmental drivers, and adaptive plasticity. The Plant Cell, 35(1), 109–124. 10.1093/plcell/koac322

Noble, L. M., Chelo, I., Guzella, T., Afonso, B., Riccardi, D. D., Ammerman, P., Dayarian, A., Carvalho, S., Crist, A., Pino-Querido, A., Shraiman, B., Rockman, M. V., & Teotónio, H. (2017). Polygenicity and Epistasis Underlie Fitness-Proximal Traits in the Caenorhabditis elegans Multiparental Experimental Evolution (CeMEE) Panel. Genetics, 207(4), 1663–1685. 10.1534/genetics.117.300406

Notredame, C., Higgins, D. G., & Heringa, J. (2000). T-coffee: A novel method for fast and accurate multiple sequence alignment1. Journal of Molecular Biology, 302(1), 205–217. 10.1006/jmbi.2000.4042

Ntalla, I., Weng, L.-C., Cartwright, J. H., Hall, A. W., Sveinbjornsson, G., Tucker, N. R., Choi, S. H., Chaffin, M. D., Roselli, C., Barnes, M. R., Mifsud, B., Warren, H. R., Hayward, C., Marten, J., Cranley, J. J., Concas, M. P., Gasparini, P., Boutin, T., Kolcic, I., … Munroe, P. B. (2020). Multi-ancestry GWAS of the electrocardiographic PR interval identifies 202 loci underlying cardiac conduction. Nature Communications, 11(1), 2542. 10.1038/s41467-020-15706-x

Ohya, Y., Sese, J., Yukawa, M., Sano, F., Nakatani, Y., Saito, T. L., Saka, A., Fukuda, T., Ishihara, S., Oka, S., Suzuki, G., Watanabe, M., Hirata, A., Ohtani, M., Sawai, H., Fraysse, N., Latgé, J.-P., François, J. M., Aebi, M., … Morishita, S. (2005). High-dimensional and large-scale phenotyping of yeast mutants. Proceedings of the National Academy of Sciences, 102(52), 19015–19020. 10.1073/pnas.0509436102

Palmer, D. S., Zhou, W., Abbott, L., Wigdor, E. M., Baya, N., Churchhouse, C., Seed, C., Poterba, T., King, D., Kanai, M., Bloemendal, A., & Neale, B. M. (2023). Analysis of genetic dominance in the UK Biobank. Science, 379(6639), 1341– 1348. 10.1126/science.abn8455

Parra, V., & Rothermel, B. A. (2017). Calcineurin signaling in the heart: The importance of time and place. Journal of Molecular and Cellular Cardiology, 103, 121–136. 10.1016/j.yjmcc.2016.12.006

Pe’er, I., Yelensky, R., Altshuler, D., & Daly, M. J. (2008). Estimation of the multiple testing burden for genomewide association studies of nearly all common variants. Genetic Epidemiology, 32(4), 381–385. 10.1002/gepi.20303

Pierotti, S., Fitzgerald, T., & Birney, E. (2025). FlexLMM: A nextflow linear mixed model framework for GWAS. Bioinformatics, btaf021. 10.1093/bioinformatics/btaf021

Pierotti, S., Welz, B., Osuna-López, M., Fitzgerald, T., Wittbrodt, J., & Birney, E. (2024). Genotype imputation in F2 crosses of inbred lines. Bioinformatics Advances, 4(1), vbae107. 10.1093/bioadv/vbae107

Polimanti, R., Kaufman, J., Zhao, H., Kranzler, H. R., Ursano, R. J., Kessler, R. C., Gelernter, J., & Stein, M. B. (2018). A genome-wide gene-by-trauma interaction study of alcohol misuse in two independent cohorts identifies PRKG1 as a risk locus. Molecular Psychiatry, 23(1), 154–160. 10.1038/mp.2017.24

Poplin, R., Ruano-Rubio, V., DePristo, M. A., Fennell, T. J., Carneiro, M. O., Auwera, G. A. V. der, Kling, D. E., Gauthier, L. D., Levy-Moonshine, A., Roazen, D., Shakir, K., Thibault, J., Chandran, S., Whelan, C., Lek, M., Gabriel, S., Daly, M. J., Neale, B., MacArthur, D. G., & Banks, E. (2018). Scaling accurate genetic variant discovery to tens of thousands of samples (p. 201178). bioRxiv. 10.1101/201178

Priori, S. G., Napolitano, C., Memmi, M., Colombi, B., Drago, F., Gasparini, M., DeSimone, L., Coltorti, F., Bloise, R., Keegan, R., Cruz Filho, F. E. S., Vignati, G., Benatar, A., & DeLogu, A. (2002). Clinical and Molecular Characterization of Patients With Catecholaminergic Polymorphic Ventricular Tachycardia. Circulation, 106(1), 69–74. 10.1161/01.CIR.0000020013.73106.D8

R Core Team. (2023). R: a language and environment for statistical computing. R Foundation for Statistical Computing. https://www.R-project.org/

Rampersaud, E., Mitchell, B. D., Pollin, T. I., Fu, M., Shen, H., O’Connell, J. R., Ducharme, J. L., Hines, S., Sack, P., Naglieri, R., Shuldiner, A. R., & Snitker, S. (2008). Physical Activity and the Association of Common FTO Gene Variants With Body Mass Index and Obesity. Archives of Internal Medicine, 168(16), 1791–1797. 10.1001/archinte.168.16.1791

Ren, J., Long, Y., Liu, R., Song, G., Li, Q., & Cui, Z. (2021). Characterization of Biological Pathways Regulating Acute Cold Resistance of Zebrafish. International Journal of Molecular Sciences, 22(6), 3028. 10.3390/ijms22063028

Robinson, J. T., Thorvaldsdóttir, H., Winckler, W., Guttman, M., Lander, E. S., Getz, G., & Mesirov, J. P. (2011). Integrative genomics viewer. Nature Biotechnology, 29(1), 24–26. 10.1038/nbt.1754

Rohde, P. D., Jensen, I. R., Sarup, P. M., Ørsted, M., Demontis, D., Sørensen, P., & Kristensen, T. N. (2019). Genetic Signatures of Drug Response Variability in Drosophila melanogaster. Genetics, 213(2), 633–650. 10.1534/genetics.119.302381

Roselli, C., Yu, M., Nauffal, V., Georges, A., Yang, Q., Love, K., Weng, L. C., Delling, F. N., Maurya, S. R., Schrölkamp, M., Tfelt-Hansen, J., Hagège, A., Jeunemaitre, X., Debette, S., Amouyel, P., Guan, W., Muehlschlegel, J. D., Body, S. C., Shah, S., … Milan, D. J. (2022). Genome-wide association study reveals novel genetic loci: A new polygenic risk score for mitral valve prolapse. European Heart Journal, 43(17), 1668–1680. 10.1093/eurheartj/ehac049

Russell, L. E., Zhou, Yitian, Almousa, Ahmed A., Sodhi, Jasleen K., Nwabufo, Chukwunonso K., & and Lauschke, V. M. (2021). Pharmacogenomics in the era of next generation sequencing – from byte to bedside. Drug Metabolism Reviews, 53(2), 253–278. 10.1080/03602532.2021.1909613

Sae-Lim, P., Boison, S. A., Brand, W., Norris, A., & Baranski, M. (2025). Genomic prediction accuracy of growth in Atlantic salmon (*Salmo salar*) when genotype-by-environment interaction is present. Aquaculture, 603, 742391. 10.1016/j.aquaculture.2025.742391

Saha, S., Spinelli, L., Castro Mondragon, J. A., Kervadec, A., Lynott, M., Kremmer, L., Roder, L., Krifa, S., Torres, M., Brun, C., Vogler, G., Bodmer, R., Colas, A. R., Ocorr, K., & Perrin, L. (2022). Genetic architecture of natural variation of cardiac performance from flies to humans. eLife, 11, e82459. 10.7554/eLife.82459

Saul, M. C., Philip, V. M., Reinholdt, L. G., & Chesler, E. J. (2019). High-Diversity Mouse Populations for Complex Traits. Trends in Genetics, 35(7), 501–514. 10.1016/j.tig.2019.04.003

Schindelin, J., Arganda-Carreras, I., Frise, E., Kaynig, V., Longair, M., Pietzsch, T., Preibisch, S., Rueden, C., Saalfeld, S., Schmid, B., Tinevez, J.-Y., White, D. J., Hartenstein, V., Eliceiri, K., Tomancak, P., & Cardona, A. (2012). Fiji: An open-source platform for biological-image analysis. Nature Methods, 9(7), 676–682. 10.1038/nmeth.2019

Scott, M. F., Fradgley, N., Bentley, A. R., Brabbs, T., Corke, F., Gardner, K. A., Horsnell, R., Howell, P., Ladejobi, O., Mackay, I. J., Mott, R., & Cockram, J. (2021). Limited haplotype diversity underlies polygenic trait architecture across 70Oyears of wheat breeding. Genome Biology, 22(1), 137. 10.1186/s13059-021-02354-7

Scott, M. F., Ladejobi, O., Amer, S., Bentley, A. R., Biernaskie, J., Boden, S. A., Clark, M., Dell’Acqua, M., Dixon, L. E., Filippi, C. V., Fradgley, N., Gardner, K. A., Mackay, I. J., O’Sullivan, D., Percival-Alwyn, L., Roorkiwal, M., Singh, R. K., Thudi, M., Varshney, R. K., … Mott, R. (2020). Multi-parent populations in crops: A toolbox integrating genomics and genetic mapping with breeding. Heredity, 125(6), 396–416. 10.1038/s41437-020-0336-6

Shah, M., de A. Inácio, M. H., Lu, C., Schiratti, P.-R., Zheng, S. L., Clement, A., de Marvao, A., Bai, W., King, A. P., Ware, J. S., Wilkins, M. R., Mielke, J., Elci, E., Kryukov, I., McGurk, K. A., Bender, C., Freitag, D. F., & O’Regan, D. P. (2023). Environmental and genetic predictors of human cardiovascular ageing. Nature Communications, 14(1), 4941. 10.1038/s41467-023-40566-6

Shiels, H. A., & Sitsapesan, R. (2015). Is there something fishy about the regulation of the ryanodine receptor in the fish heart? Experimental Physiology, 100(12), 1412–1420. 10.1113/EP085136

Sijtsma, A., Rienks, J., van der Harst, P., Navis, G., Rosmalen, J. G. M., & Dotinga, A. (2022). Cohort Profile Update: Lifelines, a three-generation cohort study and biobank. International Journal of Epidemiology, 51(5), e295–e302. 10.1093/ije/dyab257

Sitsapesan, R., Montgomery, R. A., MacLeod, K. T., & Williams, A. J. (1991). Sheep cardiac sarcoplasmic reticulum calcium-release channels: Modification of conductance and gating by temperature. The Journal of Physiology, 434, 469–488. 10.1113/jphysiol.1991.sp018481

Sleiman, Y., Lacampagne, A., & Meli, A. C. (2021). “Ryanopathies” and RyR2 dysfunctions: Can we further decipher them using in vitro human disease models? Cell Death & Disease, 12(11), 1041. 10.1038/s41419-021-04337-9

Smith, E. N., & Kruglyak, L. (2008). Gene–Environment Interaction in Yeast Gene Expression. PLOS Biology, 6(4), e83. 10.1371/journal.pbio.0060083

Snoek, B. L., Volkers, R. J. M., Nijveen, H., Petersen, C., Dirksen, P., Sterken, M. G., Nakad, R., Riksen, J. A. G., Rosenstiel, P., Stastna, J. J., Braeckman, B. P., Harvey, S. C., Schulenburg, H., & Kammenga, J. E. (2019). A multi-parent recombinant inbred line population of C. elegans allows identification of novel QTLs for complex life history traits. BMC Biology, 17(1), 24. 10.1186/s12915-019-0642-8

Souren, M., Martinez-Morales, J. R., Makri, P., Wittbrodt, B., & Wittbrodt, J. (2009). A global survey identifies novel upstream components of the Ath5 neurogenic network. Genome Biology, 10(9), R92. 10.1186/gb-2009-10-9-r92

Spivakov, M., Auer, T. O., Peravali, R., Dunham, I., Dolle, D., Fujiyama, A., Toyoda, A., Aizu, T., Minakuchi, Y., Loosli, F., Naruse, K., Birney, E., & Wittbrodt, J. (2014). Genomic and Phenotypic Characterization of a Wild Medaka Population: Towards the Establishment of an Isogenic Population Genetic Resource in Fish. G3 Genes|Genomes|Genetics, 4(3), 433–445. 10.1534/g3.113.008722

Srivastava, A., Morgan, A. P., Najarian, M. L., Sarsani, V. K., Sigmon, J. S., Shorter, J. R., Kashfeen, A., McMullan, R. C., Williams, L. H., Giusti-Rodríguez, P., Ferris, M. T., Sullivan, P., Hock, P., Miller, D. R., Bell, T. A., McMillan, L., Churchill, G. A., & de Villena, F. P.-M. (2017). Genomes of the Mouse Collaborative Cross. Genetics, 206(2), 537–556. 10.1534/genetics.116.198838

Steinberg, C., Roston, T. M., van der Werf, C., Sanatani, S., Chen, S. R. W., Wilde, A. A. M., & Krahn, A. D. (2023). RYR2-ryanodinopathies: From calcium overload to calcium deficiency. EP Europace, 25(6), euad156. 10.1093/europace/euad156

Stemmer, M., Thumberger, T., Del Sol Keyer, M., Wittbrodt, J., & Mateo, J. L. (2017). Correction: CCTop: An Intuitive, Flexible and Reliable CRISPR/Cas9 Target Prediction Tool. PLOS ONE, 12(4), e0176619. 10.1371/journal.pone.0176619

Sul, J. H., Martin, L. S., & Eskin, E. (2018). Population structure in genetic studies: Confounding factors and mixed models. PLOS Genetics, 14(12), e1007309. 10.1371/journal.pgen.1007309

Svenson, K. L., Gatti, D. M., Valdar, W., Welsh, C. E., Cheng, R., Chesler, E. J., Palmer, A. A., McMillan, L., & Churchill, G. A. (2012). High-Resolution Genetic Mapping Using the Mouse Diversity Outbred Population. Genetics, 190(2), 437–447. 10.1534/genetics.111.132597

Tegenfeldt, F., Kuznetsov, D., Manni, M., Berkeley, M., Zdobnov, E. M., & Kriventseva, E. V. (2025). OrthoDB and BUSCO update: Annotation of orthologs with wider sampling of genomes. Nucleic Acids Research, 53(D1), D516–D522. 10.1093/nar/gkae987

The Complex Trait Consortium. (2004). The Collaborative Cross, a community resource for the genetic analysis of complex traits. Nature Genetics, 36(11), 1133–1137. 10.1038/ng1104-1133

The Human Protein Atlas. (n.d.). Retrieved 1 April 2025, from https://www.proteinatlas.org/

Thumberger, T., Tavhelidse-Suck, T., Gutierrez-Triana, J. A., Cornean, A., Medert, R., Welz, B., Freichel, M., & Wittbrodt, J. (2022). Boosting targeted genome editing using the hei-tag. eLife, 11, e70558. 10.7554/eLife.70558

Tonnelé, H., & Baud, A. (2022). Genetc Effects: Accounting for diet and age. eLife, 11, e80890. 10.7554/eLife.80890

Tukey, J. W. (with Internet Archive). (1977). Exploratory data analysis. Reading, Mass.□: Addison-Wesley Pub. Co. http://archive.org/details/exploratorydataa00tuke_0

van de Vegte, Y. J., Eppinga, R. N., van der Ende, M. Y., Hagemeijer, Y. P., Mahendran, Y., Salfati, E., Smith, A. V., Tan, V. Y., Arking, D. E., Ntalla, I., Appel, E. V., Schurmann, C., Brody, J. A., Rueedi, R., Polasek, O., Sveinbjornsson, G., Lecoeur, C., Ladenvall, C., Zhao, J. H., … van der Harst, P. (2023). Genetic insights into resting heart rate and its role in cardiovascular disease. Nature Communications, 14(1), 4646. 10.1038/s41467-023-39521-2

van den Berg, M. E., Warren, H. R., Cabrera, C. P., Verweij, N., Mifsud, B., Haessler, J., Bihlmeyer, N. A., Fu, Y.-P., Weiss, S., Lin, H. J., Grarup, N., Li-Gao, R., Pistis, G., Shah, N., Brody, J. A., Müller-Nurasyid, M., Lin, H., Mei, H., Smith, A. V., … Munroe, P. B. (2017). Discovery of novel heart rate-associated loci using the Exome Chip. Human Molecular Genetics, 26(12), 2346–2363. 10.1093/hmg/ddx113

Van der Auwera, G. A., & O’Connor, B. D. (2020). Genomics in the cloud. O’Reilly Media, Inc. https://www.oreilly.com/library/view/genomics-in-the/9781491975183/

VanderWeele, T. J., & Knol, M. J. (2014). A Tutorial on Interaction. Epidemiologic Methods, 3(1), 33–72. 10.1515/em-2013-0005

Vasimuddin, Md., Misra, S., Li, H., & Aluru, S. (2019). Efficient Architecture-Aware Acceleration of BWA-MEM for Multicore Systems. 2019 IEEE International Parallel and Distributed Processing Symposium (IPDPS), 314–324. 10.1109/IPDPS.2019.00041

Verweij, N., Benjamins, J.-W., Morley, M. P., van de Vegte, Y. J., Teumer, A., Trenkwalder, T., Reinhard, W., Cappola, T. P., & van der Harst, P. (2020). The Genetic Makeup of the Electrocardiogram. Cell Systems, 11(3), 229–238.e5. 10.1016/j.cels.2020.08.005

Vilhjálmsson, B. J., & Nordborg, M. (2013). The nature of confounding in genome-wide association studies. Nature Reviews Genetics, 14(1), 1–2. 10.1038/nrg3382

Wade, C. M., & Daly, M. J. (2005). Genetic variation in laboratory mice. Nature Genetics, 37(11), 1175–1180. 10.1038/ng1666

Westerman, K. E., Majarian, T. D., Giulianini, F., Jang, D.-K., Miao, J., Florez, J. C., Chen, H., Chasman, D. I., Udler, M. S., Manning, A. K., & Cole, J. B. (2022). Variance-quantitative trait loci enable systematic discovery of gene-environment interactions for cardiometabolic serum biomarkers. Nature Communications, 13(1), 3993. 10.1038/s41467-022-31625-5

Westerman, K. E., & Sofer, T. (2024). Many roads to a gene-environment interaction. The American Journal of Human Genetics, 111(4), 626–635. 10.1016/j.ajhg.2024.03.002

Widmayer, S. J., Evans, K. S., Zdraljevic, S., & Andersen, E. C. (2022). Evaluating the power and limitations of genome-wide association studies in Caenorhabditis elegans. G3 Genes|Genomes|Genetics, 12(7), jkac114. 10.1093/g3journal/jkac114

Willis-Owen, S. A. G., & Valdar, W. (2009). Deciphering gene-environment interactions through mouse models of allergic asthma. Journal of Allergy and Clinical Immunology, 123(1), 14–23. 10.1016/j.jaci.2008.09.016

Wittbrodt, J., Shima, A., & Schartl, M. (2002). Medaka—A model organism from the far east. Nature Reviews Genetics, 3(1), 53–64. 10.1038/nrg704

Wright, K. M., Deighan, A. G., Di Francesco, A., Freund, A., Jojic, V., Churchill, G. A., & Raj, A. (2022). Age and diet shape the genetic architecture of body weight in diversity outbred mice. eLife, 11, e64329. 10.7554/eLife.64329

Xhaard, C., Dandine-Roulland, C., Villemereuil, P. de, Floch, E. L., Bacq-Daian, D., Machu, J.-L., Ferreira, J. P., Deleuze, J.-F., Zannad, F., Rossignol, P., & Girerd, N. (2021). Heritability of a resting heart rate in a 20-year follow-up family cohort with GWAS data: Insights from the STANISLAS cohort. European Journal of Preventive Cardiology, 28(12), 1334– 1341. 10.1177/2047487319890763

Yang, J., Benyamin, B., McEvoy, B. P., Gordon, S., Henders, A. K., Nyholt, D. R., Madden, P. A., Heath, A. C., Martin, N. G., Montgomery, G. W., Goddard, M. E., & Visscher, P. M. (2010). Common SNPs explain a large proportion of the heritability for human height. Nature Genetics, 42(7), 565–569. 10.1038/ng.608

Yengo, L., Vedantam, S., Marouli, E., Sidorenko, J., Bartell, E., Sakaue, S., Graff, M., Eliasen, A. U., Jiang, Y., Raghavan, S., Miao, J., Arias, J. D., Graham, S. E., Mukamel, R. E., Spracklen, C. N., Yin, X., Chen, S.-H., Ferreira, T., Highland, H. H., … Hirschhorn, J. N. (2022). A saturated map of common genetic variants associated with human height. Nature, 610(7933), 704–712. 10.1038/s41586-022-05275-y

